# Chemical phylogenetics of the staphylococcal quorum sensing landscape

**DOI:** 10.1101/2021.04.18.440348

**Authors:** Bengt H. Gless, Benjamin S. Bejder, Ludovica Vitolo, Leonor Marques, Paal S. Andersen, Martin S. Bojer, Hanne Ingmer, Christian A. Olsen

## Abstract

Staphylococci utilize secreted autoinducing peptides (AIPs) to regulate group behaviour through a process called quorum sensing (QS). Here, we survey the QS interaction landscape within the *Staphylococcus* genus by assembling a unique compound collection, comprising all the currently known AIPs. These ribosomally synthesized and posttranslationally modified peptides (RiPPs) were obtained by chemical synthesis and mapping of their ability to modulate QS was evaluated using reporter strains of common human and animal colonizing pathogens (*S. aureus, S. epidermidis, S. lugdunensis*). The resulting map of >200 native QS interactions provides a holistic view of nodes that contribute to the complex signalling network within the *Staphylococcus* genus. This overview reveals surprising cross-species QS induction and identify the first pan-inhibitory AIP, which is then shown to attenuate MRSA induced skin infection in a mouse model. Our results expose a complex universe of possible staphylococcal interactions and provide further impetus for development of therapeutics based on QS modulators targeting antibiotic resistant pathogens.

---

Staphylococci are common colonizers of humans and animals with a genus currently consisting of 57 species and 26 subspecies^1^. The genus represents some of the most abundant microbes found in the human microbiota, where the bacteria are involved in complex interactions with the host and other bacteria^2^. A prominent example is the pathogenic species *Staphylococcus aureus* (SA), which is carried asymptomatically by 30% of the human population causing 76% of skin and soft-tissue infections, despite being a relatively poor colonizer of the skin^3^. Emerging methicillin-resistant strains of SA (MRSA) are causing pressure on the public health sector^4,5^ and heavily contribute to the threat of a post-antibiotic era^6^. While SA is mainly colonizing the nasal cavity, the human skin is highly populated by coagulase-negative staphylococci (CoNS)^3,7,8^ of which the most abundant species, *S. epidermidis* (SE), can be found on the skin of nearly all humans^9^. Its roles as symbiont are manifold^10^ with recent studies showing a beneficial role for the host^11,12^, while at the same time causing medical device infections^9^. Other CoNS, such as *S. hominis*, *S. haemolyticus* and *S. lugdunensis* (SL),^3,7,8^ are also common commensal bacteria, which contribute to the complexity of interactions that shape the skin microbiome^13^. Quorum sensing (QS) plays an important role in the transition from harmless skin colonizer to invasive pathogen^14,15^ and this is regulated through the secretion and detection of autoinducing peptides (AIPs), containing a thiolactone (lactone for *S. intermedius* group)^16^. The AIP-mediated QS machinery is encoded by a chromosomal locus termed *accessory gene regulator* (*agr*), which controls the expression of virulence factors, Agr proteins, and AIPs (Extended Data Fig. 1a)^14,15^. Cross-species QS interference in human microbiota has received therapeutic interest as it can repress virulence gene expression in pathogenic staphylococci including *Staphylococcus aureus*^17,18^. However, the specific interactions that occur between co-inhabiting species have only been sporadically investigated^19–24^. Studies have investigated the AIPs of SA^25–27^as well as non-aureus staphylococci^20–24^as QS inhibitors and a recent pioneering study even investigated the disease outcome of SA infection upon co-colonization with the AIP producing commensal bacterium *S. hominis* in a human clinical trial^28^. Another skin commensal bacterium *S. lugdunensis* (SL) has proven pathogenic by causing severe endocarditis^29^ and has been targeted in one previous study^30^.

However, the effect of non-cognate AIPs, except those from *S. aureus*, on different *agr* systems, including that of the common skin colonizer *S: epidermidis* (SE) has received surprisingly limited attention despite its abundance on the human skin^31–33^. The growing number of identified AIPs and the lack of information about bacterial crosstalk therefore encouraged us to investigate interactions of all known AIPs (**1**–**26**; Extended Fig. 2) with SA *agr*-I–IV, SE *agr*-I–III, and SL *agr*-I

## Identification of new autoinducing peptides using an improved trapping protocol

We recently developed a method for the rapid identification of AIPs from bacterial supernatants (see Extended Fig. 3), which led to a substantial increase in the number of known AIPs^34^. The most time-consuming step in this procedure is the lyophilization of the supernatant. Therefore we optimized the protocol to allow trapping of the AIPs directly from the bacterial supernatant by adjusting the pH of the supernatant and adding reducing reagent (Supplementary Figs. 1–4)^17,35^. We identified three new AIPs, namely *S. pasteuri* AIP-I (**15**), *S. succinus* AIP-I (**16**), and *S. cohnii* AIP-I (**17**) (Fig. 1a, Supplementary Fig. 5–7), which brings the number of known staphylococcal AIPs to 28 [26 unique structures (**1**–**26**)], originating from 19 species and covering 5 of the 6 phylogenetic species groups as classified trough multi-locus analysis (Extended Data Fig. 2)^36^.

**Fig. 1.**
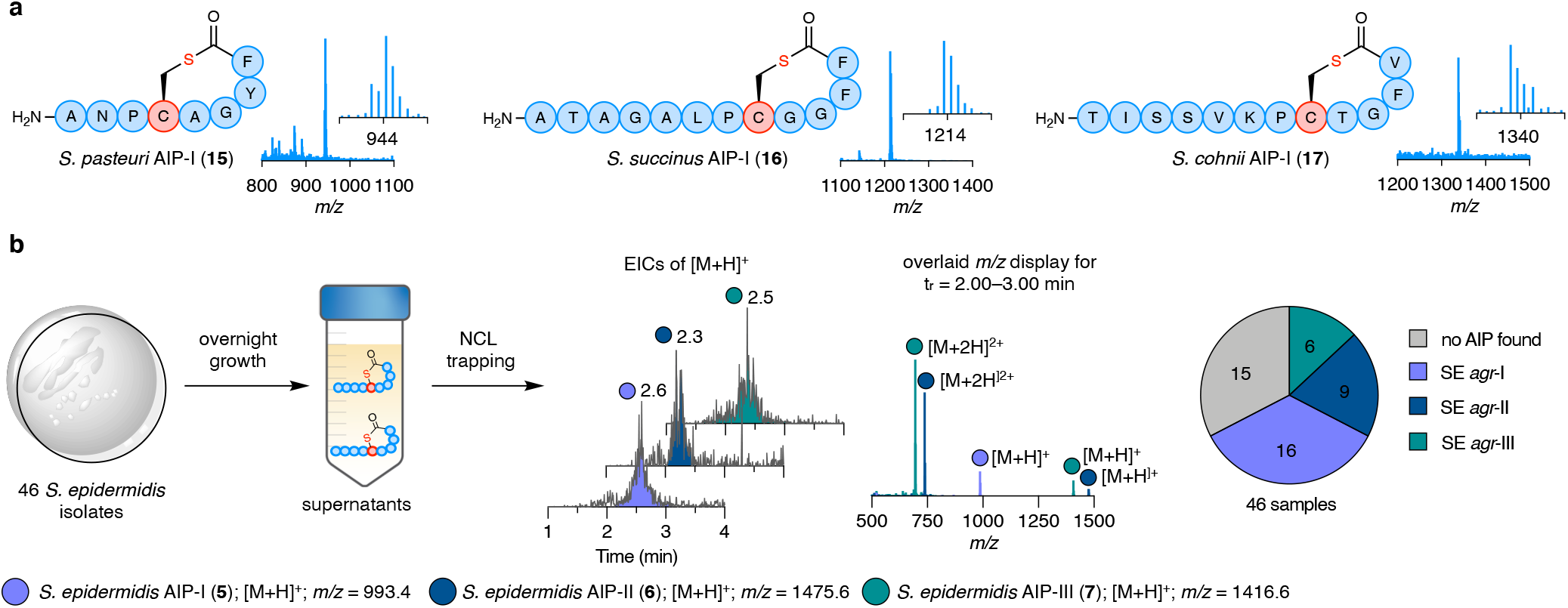
Identification of AIPs through native chemical ligation (NCL) trapping. **a**, Newly identified AIPs of three CoNS species; *S. pasteuri* AIP-I (**15**), *S. succinus* AIP-I (**16**), and *S. cohnii* AIP-I (**17**). **b**, NCL trapping performed on bacterial supernatants of 46 isolates of SE from 10 human nasal swabs. The three trapped *Se*-AIPs (**5**–**7**) in an analytic LC-MS run eluted between 2.00–3.00 min, as shown for the extracted ion chromatograms (EICs) of *m/z* [M+H]^+^ (t_r_ = 2.6, 2.3 and 2.5 min). The nasal swab strains were analyzed through combined *m/z* display (t_r_ = 2.00–3.00 min) and inspection for [M+H]^+^ and [M+2H]^2+^ of the corresponding trapped *Se*-AIPs.

The substantially less time-consuming identification of AIPs also enabled rapid parallel analysis of the *agr* groups of 46 SE isolates from a collection of 10 human nasal swabs (Fig. 1b, Supplementary Fig. 8–10), furnishing an identification frequency of close to 70% (31 of 46). The SE *agr*-I group was the most prevalent variant in the ensemble of isolates (16), followed by *agr*-II (9) and *agr*-III (6). None of the analyzed strains secreted AIPs corresponding to the predicted AIP of SE *agr*-IV, which remains unidentified^37^. Interestingly, several nasal swaps contained more than one *agr* variant of SE, which was also previously found by PCR analysis of healthy human nasal swabs^38^, underscoring the complex composition of the staphylococcal colonization.

The substantially less time-consuming identification of AIPs also enabled rapid parallel analysis of the *agr* groups of 46 SE isolates from a collection of 10 human nasal swabs (Fig. 1b, Supplementary Fig. 8–10), furnishing an identification frequency of close to 70% (31 of 46). The SE *agr*-I group was the most prevalent variant in the ensemble of isolates (16), followed by *agr*-II (9) and *agr*-III (6). None of the analyzed strains secreted AIPs corresponding to the predicted AIP of SE *agr*-IV, which remains unidentified^37^. Interestingly, several nasal swaps contained more than one *agr* variant of SE, which was also previously found by PCR analysis of healthy human nasal swabs^38^, underscoring the complex composition of the staphylococcal colonization.

## Quorum sensing interaction map of all known AIPs

Having compiled all 26 currently known AIPs (see the supplementary information for synthetic details), we first determined the half maximal inhibitory concentrations (IC_50_) for QS inhibition against SA *agr*-I–IV of the 12 newly synthesized AIPs in the previously applied *β*-lactamase reporter strain assay (Extended Data Table 1; Supplementary Fig. 11–14) to compare their potencies to the previously tested AIPs^34^. However, to include assessment of the QS modulation properties against SE and SL, we turned to fluorescent reporter strains. Thus, screening of the 26 AIPs and *Sa*-AIP-III D4A (**27**), a known pan-inhibtior of SA *agr-*I–IV,^26^ against 8 GFP/YFP-producing reporter strains of SA, SE, and SL provides a comparable data set across all tested strains (Fig 2a, Supplementary Figs. 15–20), which correlated well with data points from previously reported QS interference studies (Supplementary Table 1 and 2).

**Fig. 2.**
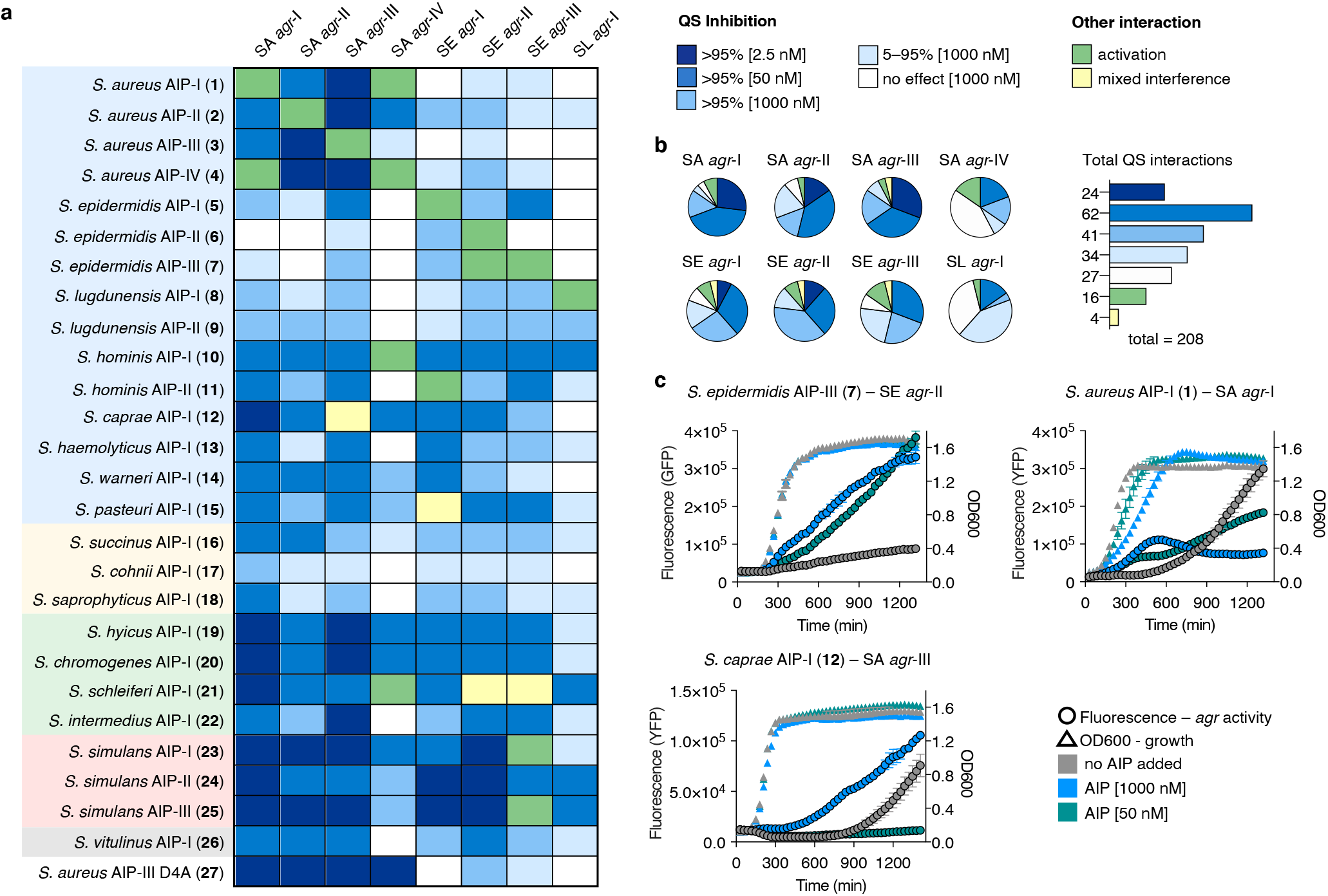
Quorum sensing (QS) interaction map. **a**, 27 synthetic AIPs were tested at several concentration (1000 nM, 50 nM, 2.5 nM) against fluorescent reporter strains of *S. aureus* (SA) *agr*-I–IV, *S. epidermidis* (SE) *agr*-I–III and *S. lugdunensis* (SL) *agr*-I to assess their QS modulation abilities. Blue shading of boxes represents different potencies of QS inhibition (>95% at 2.5 nM, 50 nM and 1000 nM or 5–95% at 1000 nM), white boxes represent no interaction at 1000 nM, green boxes represent QS activation and yellow boxes represent mixed interference (activation at 1000 nM and inhibition at 50 nM). Shaded boxes represent the phylogenetic species groups as defined by Lamers *et al.* ^36^ with following colors: Epidermidis-Aureus, blue; Saprophyticus, tan; Simulans, red; Hyicus-Intermedius, green; Sciuri, gray. All assays were performed as technical triplicates in three individual biological assays (n = 3). **b**, Pie chart representing the summary of QS interaction for each species and *agr* group with the 26 native AIPs (excluding **27**) as well as the overall summary of QS interactions following the heat map color scheme. **c**, Overnight growth (OD_600_) and fluorescence curves recorded of bacterial cultures treated with selected AIPs at 1000 nM or 50 nM and without addition of AIPs. The graphs represent single experiments in technical duplicate.

The 208 native QS interactions were predominantly inhibitory (161 of 208), with only 27 out of 208 (13%) showing no effect at 1 M (Fig. 2b). The SA *agr*-I–III variants were highly susceptible to most tested AIPs also representing the majority of interactions reaching >95% inhibition at 2.5 nM (19 of 24). In agreement with our-lactamase data set (Extended Data Table 1), SA *agr*-IV is significantly less susceptible to inhibition by other AIPs compared to SA *agr*-I–III and only the synthetic inhibitor **27** was able to fully suppress *agr* activity of that particular strain at 2.5 nM. The SE *agr*-I–III variants were nearly equally susceptible to the inhibition by AIPs of other species, as observed for as SA *agr*-I–III, but the majority of inhibitory interactions required higher AIP concentrations to achieve full inhibition. SL *agr*-I was the least affected by AIPs from other staphylococci with only 4 of 26 interactions being fully inhibitory at 50 nM. The AIPs of SA (**1**–**4**) showed surprisingly little effect against SE *agr*-I–III and vice versa with the exception of *Se*-AIP-I (**5**) for SA *agr*-III. This stands in contrast to the majority of AIPs, which showed no strong tendency to inhibit only SA or SE but rather showed similar inhibition profiles against both, although generally with increased potencies against SA (Fig. 2a).

The screening revealed that several AIPs increased the fluorescence readout compared to the control wells (Fig. 2a, shown in green) and we therefore monitored growth and fluorescence continuously overnight. Surprisingly, we found *Se*-AIP-III (**7**) to be an activator of SE *agr*-II, in contrast to previous experiments with bacterial supernatant where the AIP had no effect (Fig. 2c)^35^. Further, several cross-species activators were discovered: *S. hominis* AIP-II (**11**) activated SE *agr*-I and *S. simulans* AIP-I (**23**) and AIP-III (**25**) activated SE *agr*-III (Supplementary Fig. 22). In addition, we monitored all known activators in the same manner and observed earlier increase in fluorescence development, which was often accompanied by a delay in growth as determined by OD_600_ values (Fig. 2c and Supplementary Figs. 21 and 22). The previously inconsistent inhibition behavior of *S. caprae* AIP-I (**12**) against SA *agr*-III (Extended Data Table 1, Supplementary Fig. 13) was also observed with the fluorescent reporter strain, where activation occurred at 1 μM while lower concentrations of the AIP led to inhibition (Fig. 2c and Supplementary Fig. 17). The same mixed interference was seen for *S. pasteuri* AIP-I (**15**) against SE *agr*-I (Supplementary Fig. 17) and for *S. schleiferi* AIP-I (**21**), which showed 70% inhibition of SE *agr*-II and III from 2.5–1000 nM. The non-natural SA pan-inhibitor **27** was a highly potent inhibitor of SA *agr-*I–IV, as expected, but showed low levels of inhibition against SE *agr*-I–III and SL *agr*-I. In contrast, the QS interaction map revealed pan-inhibitors of all 8 staphylococcal strains, namely *S. hyicus* AIP-I (**19**) and *S. chromogenes* AIP-I (**20**), which potently inhibited all SA and SE *agr* variants but only weakly inhibited SL *agr*-I as well as *S. simulans* AIP-II (**24**) that inhibited all 8 *agr* systems (Fig. 2a).

## Quorum sensing interactions in the context of phylogeny of *agr* systems

The diversity of AIPs in the *Staphylococcus* genus can be estimated by the number of deposited AgrD sequences in genomic databases. We collected the majority of available AgrD sequences and set as a requirement for having a unique *agr* group that they possessed a unique AIP-containing sequence (Extended Data Fig. 1b). A total of 125 AgrD sequences were found, belonging to 55 species, which were used to construct a maximum likelihood tree (Fig. 4a). AgrD sequences with substitutions in the C-terminal recognition sequences and the *N*-terminal leader peptide were not considered as a unique *agr* group and were therefore not included. The *agr* groups of identified AIPs were named in their chronological order of their discovery and *agr* groups with so far unknown AIPs were ranked in order of their number of accessions in the Protein Database (Supplementary Table 3). The AgrD sequences clustered to large extent into the 6 species groups that were previously defined by multi-locus analysis; albeit, with some exceptions^36^. Firstly, AgrD of SA *agr*-II is highly divergent from the other SA *agr* groups and is located on a single clade close to the Saprophyticus species group as found in a previous analysis of AgrD sequences^39^. The AgrD peptides of SL *agr*-I–II and *S. caprae agr*-I–II are phylogenetically divergent from the remaining Epidermidis-Aureus species group and show closer sequence similarity to the Hyicus-Intermedius and Simulans species groups, respectively. All AgrD sequences belonging to the Hyicus-Intermedius, the Simulans, and the Sciuri species groups clustered together and diverged early from the Epidermidis-Aureus and the Saprophyticus groups (Fig. 4a). The largest coverage of presently identified AIPs is within the Epidermidis-Aureus groups (17 of 45 unique *agr* groups), showing that most common species that colonize humans belong to this group. The QS interactions found in this group are relatively diverse, as it contains the three species against which the QS interaction screen was performed (Fig. 4b). The majority of AIPs that exhibit a lack of QS interference with SL *agr*-I belong to this species group.

The second largest species group, Saprophyticus, contains the smallest number of identified AIPs (3 of 37 unique *agr* groups). Two of these were identified in this study and all three AIPs proved to exhibit relatively weak interactions with SA, SE and SL. The third largest species group, Hyicus-Intermedius, contains several species that are often isolated from animal hosts, such as *S. hyicus, S. schleiferi,* and *S. chromogenes*^40^. To date, 4 AIPs have been identified from the 25 unique *agr* groups, which are all potent interactors with the *agr* systems of SA, SE, and SL. The three known AIPs from the smaller Simulans species group (just 7 unique *agr* groups from 4 species) belong to *S. simulans* and show exceptionally strong QS interference with all 8 tested reporter strains.

Finally, the only discovered AIP of the Sciuri species group (7 *agr* groups from 5 species) exhibited moderate activity against most of the tested strains and none of the two *agr* groups in the Auricularis species group have had their AIPs identified yet.

To further examine the diversity and relatedness of secreted AIPs, we created a maximum likelihood tree of the 12 amino acid-long AIP-containing sequences of all found AgrD peptides (Extended Data Fig. 4). Not unexpectedly, the clustering into species groups became less pronounced but several interesting observations surfaced. Notably, *S. schleiferi agr*-I and *S. hominis agr*-I, which both secrete activators of SA *agr*-IV, show high levels of relatedness and appear in close proximity in the cladogram. Also, high sequence similarity was revealed for SA *agr*-I, III, and IV to the Simulans species group, which diverge directly from the same clade. The most striking difference was the clustering of SE *agr*-II–IV into the Simulans species group. Particularly when considering that all interactions with full inhibition of SE at 2.5 nM concentration of tested AIPs were elicited by *S. simulans* AIPs and the cross-species activation of SE *agr*-III was caused by two of the *S. simulans* AIPs.

## The autoinducing peptides from *S. simulans* and chemical phylogenetics

The most potent group of QS inhibitors revealed through our QS interaction mapping were the three AIPs produced by *S. simulans* (Figs. 2a and 4a). We determined the IC_50_ values of *S. simulans* AIP-I–III (**23**–**25**) by dose-response experiments against SA *agr*-I–IV, SE *agr*-I–III, and SL *agr*-I, which further confirmed their potent inhibitory properties (Extended Data Table 2 and Supplementary Figs. 23–25). The *Ss*-AIP-I (**23**) and *Ss*-AIP-III (**25**) have similar QS interaction profiles, where both show high potencies against all SA *agr*-I–III groups (IC_50_ <1 nM) and activate SE *agr*-III. The *Ss*-AIP-II (**24**), on the other hand, inhibited all 8 *agr* groups (SA, SE, SL); albeit, with slightly lower potencies against SA *agr*-II and SA *agr*-III compared to the other two *S. simulans* AIPs (Extended Data Table 2). The *Ss*-AIP-I (**23**) and *Ss*-AIP-III (**25**) have the same exotail (Fig. 4a), which is also shared by *Se*-AIP-III (**7**) and could therefore be the determining feature for the activation of SE *agr*-III. That *Ss*-AIP-II (**24**) has a single amino acid substitution (N3Y) and acts as potent inhibitor of this system (IC_50_ = 0.6 nM) supports that hypothesis further. Similarly, it is tempting to speculate that the identical macrocycle sequence of *Ss*-AIP-II (**24**) and *Ss*-AIP-III (**25**) may be important for the potent inhibition of SL *agr*-I by both AIPs. The potent inhibition of SE *agr*-I–II by *Ss*-AIP-III (**25**) at subnanomolar IC_50_ values (0.14 nM and 0.11 nM) represent the most potent native QS interactions between staphylococci reported, to the best of our knowledge. In addition, the IC_50_ value of *Ss*-AIP-III (**25**) against SL *agr*-I (0.48 nM) represents a more than 400-fold increase in potency compared to previously reported inhibitors of this species group^30^.

We then performed a chemical phylogenetic analysis by combining the phylogenetic trees of the 8 AgrC receptors (SA, SE, SL) with determined IC_50_ values for the cognate AIPs of SA *agr*-I (**1**), SE *agr*-I (**5**), and SL *agr*-I (**8**) as well as pan-inhibitor *Ss*-AIP-II (**24**) (Fig. 4b, Extended Data Table 2 and Supplementary Figs. 26–28). The AgrC receptors of SA grouped together, with *agr*-II being more distantly related to the other *agr* groups, creating a similar picture as observed in the phylogeny of the full AgrD sequences (Fig. 3a).

**Fig. 3.**
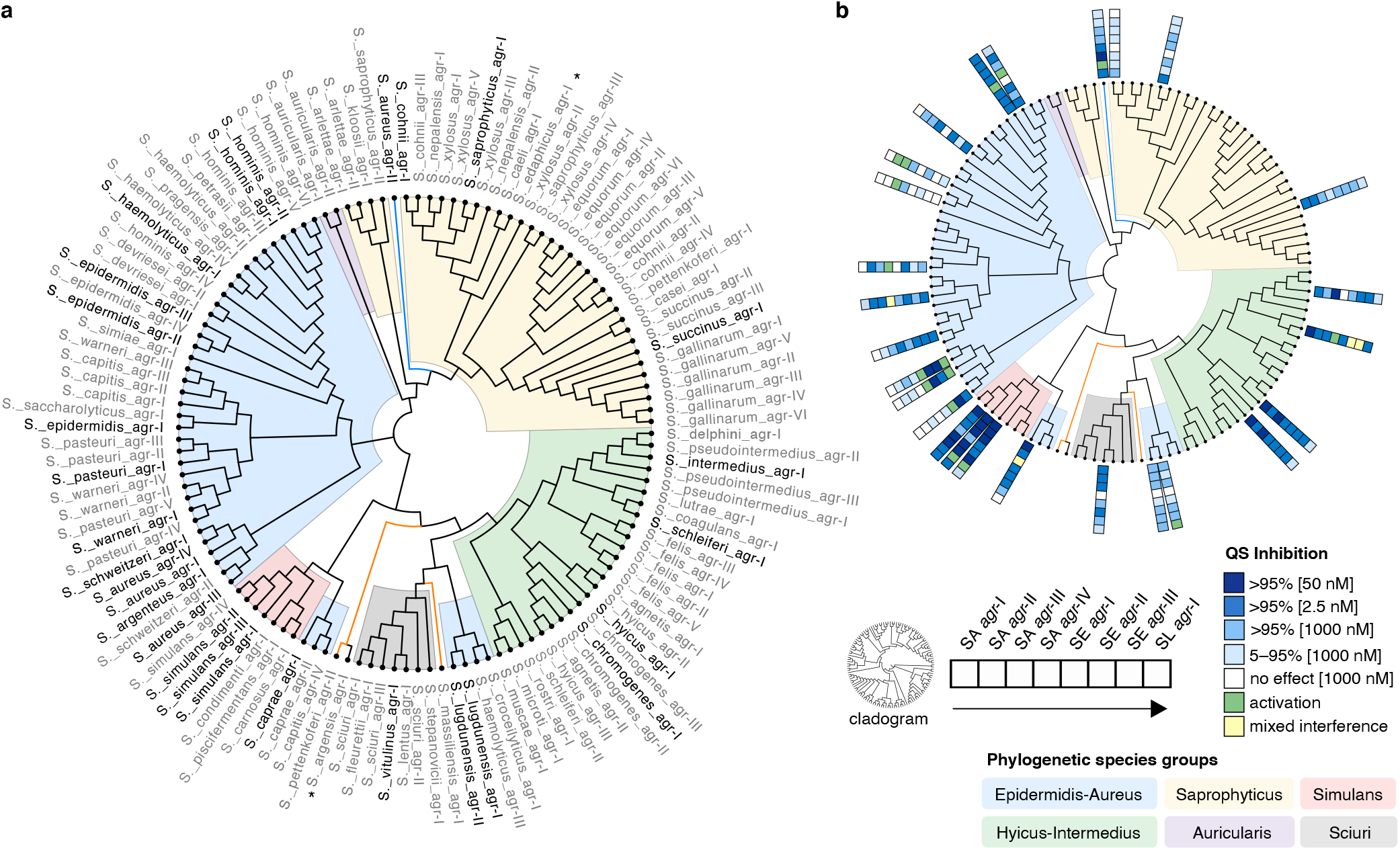
Phylogeny of the AIP-precursor peptides AgrD. **a**, Maximum likelihood tree (bootstrap n = 500) created from 125 AgrD sequences with unique AIP-containing sequences. Names of *agr* groups with known AIPs are black and grey with unknown AIPs. Phylogenetic species groups as defined by Lamers *et al.*^36^ are colored as follows: Epidermidis-Aureus, blue; Saprophyticus, tan; Simulans, red; Hyicus-Intermedius, green; Auicularis, purple; Sciuri, gray. *Species not yet assigned to species group. **b**, Cladogram overlaid with QS interactions of known AIPs against SA *agr*-I–IV, SE *agr*-I–III and SL *agr*-I. The interactions of one AIP are represented as colored boxes with SA *agr*-I closest to the cladogram as depicted in the figure.

**Fig. 4.**
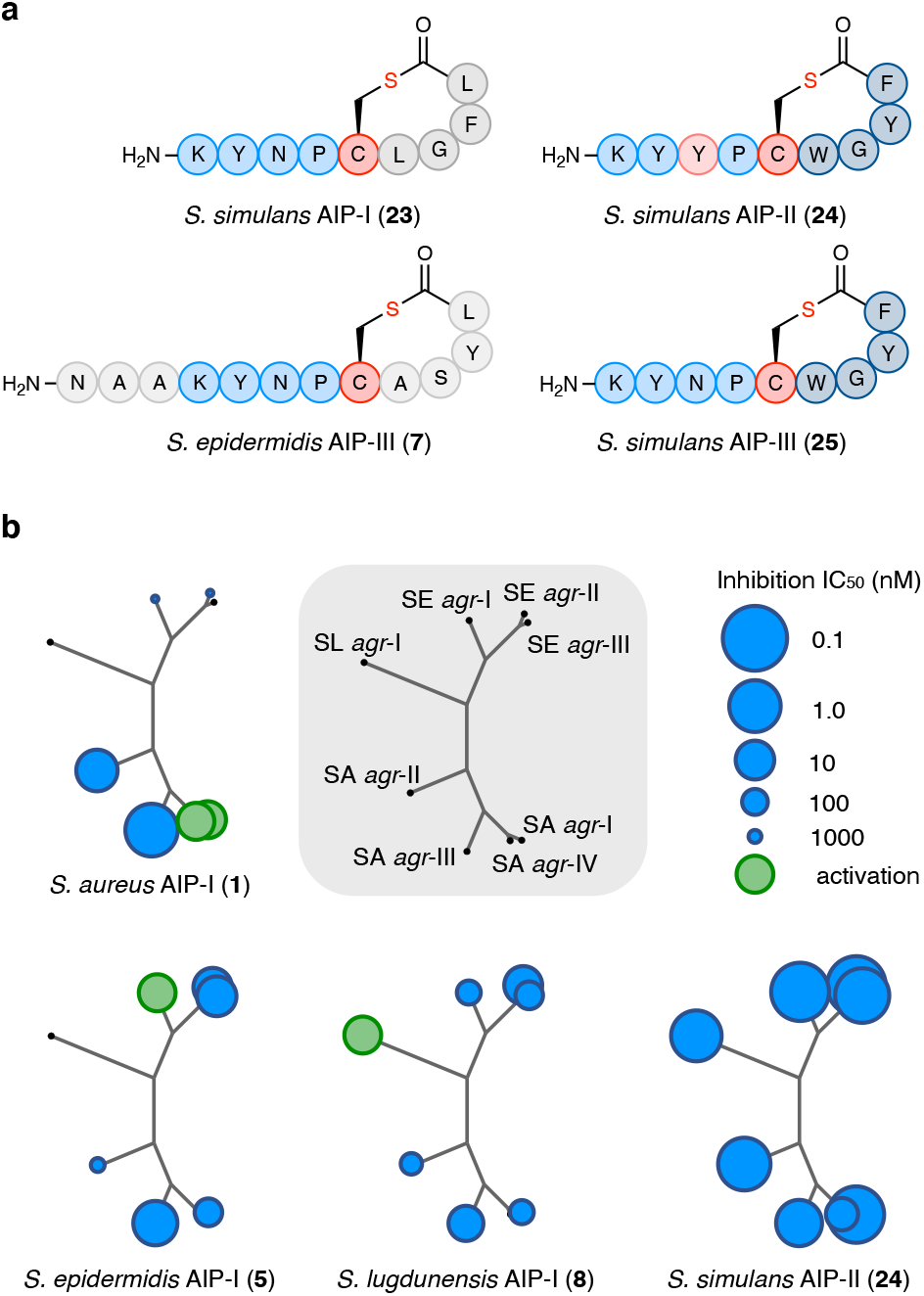
*S. simulans* AIP structures and chemical phylogenetic analyses. **a**, The structures of *S. simulans* AIP-I–III (**23**–**25**) and SE-AIP-III (**7**) show the same exotail amino acid sequence (KYNP) in **23**, **25**, and **7** as well as the same macrocyle amino acid sequence (CWGYF) for **24** and **25**. **b**, Phylogenetic trees (bootstrap n = 500) of the AgrC receptor of SA *agr*-I–IV, SE *agr*-I–III, and SL *agr*-I were overlaid with IC50 values determined through full-dose response curves (Extended Data Table 2) highlighting the different QS interference patterns of cognate AIPs (**1**, **5**, and **8**) and *S. simulans* AIP-II (**24**).

It is noteworthy that the receptors of *agr*-I and *agr*-IV share 87% sequence similarity despite their greatly differing susceptibly to QS interference (Fig. 2b)^14^. The AgrC receptors of SE also group together, with *agr*-I being more distantly related to *agr*-II/III, which parallels the AgrD clustering, while the AgrC receptor of SL is highly divergent from the other two species. The selective inhibition pattern of *Sa*-AIP-I (**1**) also observed in the QS interaction screening, appeared even more pronounced with nearly no interference with SE and SL (Fig. 4b). The *Se*-AIP-I (**5**) showed higher potency against SA *agr-*III than SE *agr*-II and III but no interference with SL *agr*-I. The cognate AIP of SL *agr*-I (**8**) was active against all other *agr* groups, except SA *agr*-IV. In contrast, *Ss*-AIP-II (**24**) showed strong inhibitory properties in nearly all AIP–AgrC interactions regardless of the receptor phylogeny.

## An autoinducing peptide from *S. simulans* attenuates MRSA in mouse skin model

Next, we tested *Ss*-AIP-II (**24**) in an *in vivo* MRSA (*agr*-I) mouse skin infection model (Fig. 5a–d). The importance of a functioning *agr* system for SA during infection to evade the immune response has been established and it was shown that inhibition of *agr* during early stages of infection can lead to improved disease outcome 48–72 h after its initiation^41^. Thus, *Ss*-AIP-II (**24**) (100 μM) added to the MRSA inoculum (10^7^ CFU) applied on the skin was compared to vehicle and daily treatment with the commercial antibacterial product Fucidin^®^ (2% fusidic acid). A significant reduction in the skin lesion size was observed after 48 h and 96 h for mice treated with *Ss*-AIP-II (**24**) (*P =*0.0174 and *P =*0.0249) as well as fusidic acid (*P =*0.0108 and *P =*0.0202) compared to the vehicle control (Fig. 5a and c). Further, a significant decrease in bacterial load (~60-fold, *P =*0.009) was observed for mice treated with *Ss*-AIP-II (**24**) compared to vehicle control after 4 days, which was comparable to daily treatment with fusidic acid (~38-fold, *P =*0.0168) (Fig. 5b). No statistical difference between treatment or vehicle or time of the infection was found for the body weight, which was expected as no systemic infections were observed (Fig. 5d). These results are highly encouraging for the prospects of anti-virulence treatments of staphylococcal infections with non-antibiotic peptides. The higher extent of bacterial clearance is most likely a result of a more efficient immune response towards non-virulent MRSA bacteria.

**Fig. 5.**
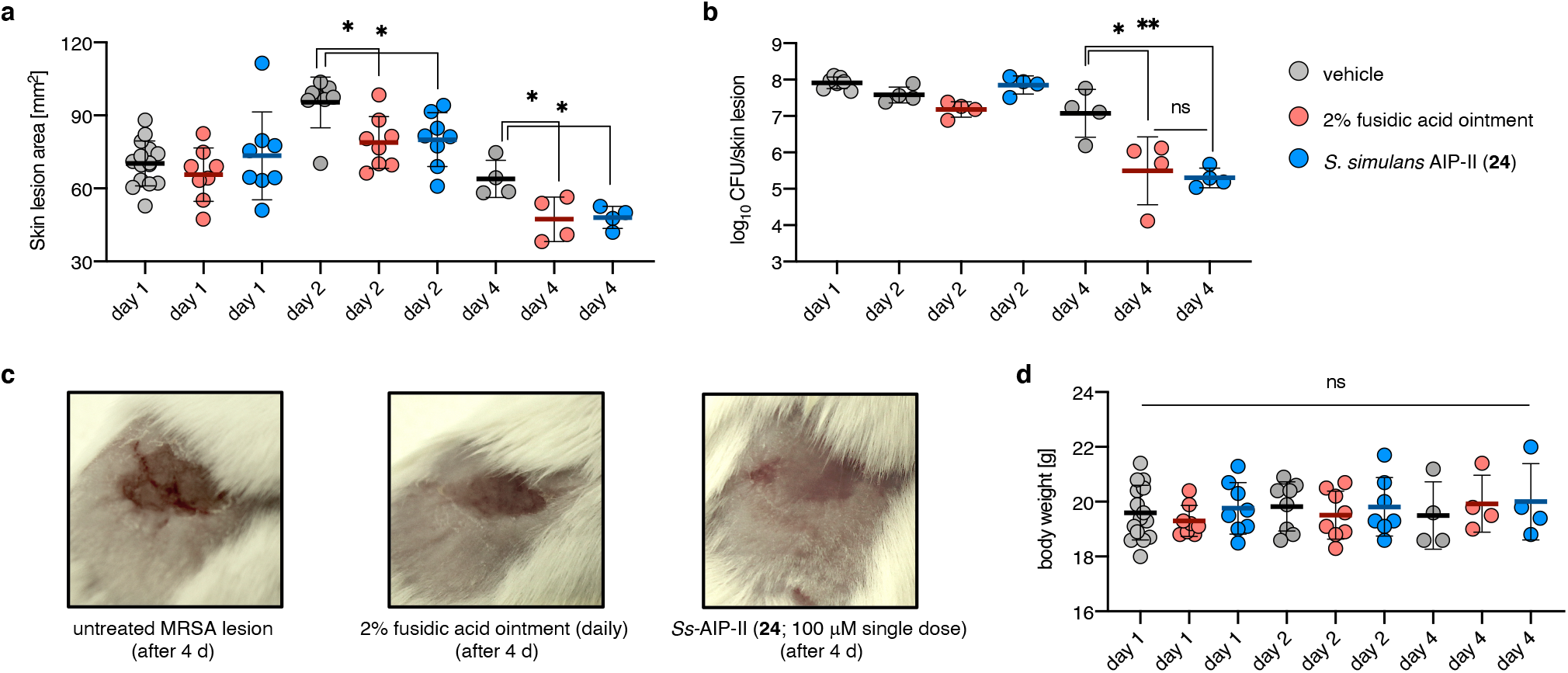
*S. simulans* AIP-II attenuates MRSA infection in murine skin model. **a**, Murine skin infection model performed with vehicle control group [day 1 (n = 16), day 2 (n = 8), day 4 (n = 4)], fusidic acid (daily application of a 38.7 mM ointment (Fucidin) [day 2 (n = 8) and 4 (n = 4)] and 24 (single treatment at day 0 at 100 μM [day 2 (n = 8) and 4 (n = 4)]. MRSA inoculum: 10^7^ CFU. Skin lesions measured on day 1, 2, and 4 showing significant reduction in lesion size for mice treated with fusidic acid and 24. **b**, CFU count determined after day 1, 2, and 4 showing significant reduction in bacterial load per skin lesion. **c**, Representative pictures taken of MRSA lesions at day 4 for untreated and treated mice. **d**, Body weight measured at day 1, 2, and 4 showing no statistically significant differences. Data is presented as mean and error bars represent the standard deviation (SD) of the mean. *P* > 0.05 (ns), *P* < 0.05 (*), *P* < 0.01 (**).

## Discussion

Altering gene expression of competing staphylococci through secreted AIPs represents a powerful ability of the *agr* system next to its cognate function of “sensing” cell density. Even though *agr* loci and AgrD peptides can be found in the genomes of all species it is not known if and to what extent these systems are functional. However, 21 AIPs from 15 CoNS have now been identified from bacterial supernatants pointing towards a broad utilization of the *agr* system, including secretion and detection of AIPs as well as potential QS interference. The colonization of humans and animals by a wide range of staphylococcal species emphasizes that many species share the same environments, providing an arena for biologically relevant inter-staphylococcal interactions, such as QS interference. We report herein an improvement of our previously developed method for AIP identification, which allows direct trapping of AIPs in untreated supernatants, which will further help to elucidate the complex network of QS interactions. The faster analysis rate enabled us to rapidly screen 46 SE samples from human nasal swabs with an identification frequency of more than 70% of the samples. In addition, the method was used to identify 3 new AIPs, *S. pasteuri* AIP-I (**15**), *S. succinus* AIP-I (**16**), and *S. cohnii* AIP-I (**17**) that helped improve coverage of the various species groups. Combined with the previously identified AIPs, we determined QS modulating properties against fluorescent reporter strains of SA *agr*-I–IV, SE *agr*-I–III and SL *agr*-I to provide a map of >200 QS interactions. This represents the largest systematic analysis of the currently known staphylococcal QS landscape and reveals extensive promiscuity of staphylococcal crosstalk, leading to the discovery of several cross-species activators as well as pan-inhibitors against all the 8 tested *agr* systems. To further elucidate the diversity of AIPs from the *Staphylococcus* genus and the phylogenetic relatedness between *agr* groups, we collected 125 AgrD sequences of 55 species and created cladograms that together with the QS interaction data showed intriguing correlations between sequence similarity and bioactivity. For example, the interactions between the AIPs of *S. simulans* and the *agr* system of SE were highly sequence sensitive with a single substitution deciding between potent inhibitor and activator of SE *agr*-III. We could further show a significant effect of an inhibitory AIP, produced by *S. simulans* (**24**), on the colonization and pathogenesis of MRSA, highlighting the power of gene repression through QS inhibition^21,23,24,28^. In addition, the interactions of *S. simulans* AIPs with SL *agr*-I represents the first highly potent inhibitors of QS in SL with a 400-fold increase in potency compared to the previously reported compounds.

The impact of the substantial native QS interference among commensal staphylococci in human microbiota may reflect the considerable overlap in living space but this remains to be explored further. However, our results highlight the potential importance of the *agr* system and cross-species interference on the colonization of commensal staphylococci and on the pathogenesis of for example SA.

It is our hope that the mapping of cross-species QS interactions initiated in the present work will help provide insight into the roles of *agr* systems in future investigations. Furthermore, our findings highlight the potential utility of natural scaffolds as a promising platform for the development of inhibitors for anti-virulence treatment of *Staphylococcus* infections.

## Methods

The chemical synthesis details, characterization data and copies of NMR spectra are provided in the Supplementary Information.

### Reagents and materials

Bacteria were cultured in tryptic soya broth (TSB) medium and on tryptic soya agar (TSA) plates (both Oxoid) supplemented with 10 μg/ml chloramphenicol (CAM) or erythromycin (ERM) (Sigma-Aldrich) when appropriate. *Nα*-Fmoc protected and*α-N*-Boc protected amino acids, 2-(1*H*-Benzotriazol-1-yl)-1,1,3,3-tetramethyluronoium hexafluorophosphate (HBTU) and 2-(1*H*-7-Azabenzotriazol-1-yl)-1,1,3,3-tetramethyluronoium hexafluorphosphate (HATU) were obtained from ChemImpex and PepChem. Aminomethyl ChemMatrix resin was obtained from PCAS BioMatrix. Aminomethyl PEGA resin, *N*,*N*-diisopropylethylamine, 4-nitrophenylchloroformate, maleic acid (qNMR grade) and nitrocefin were obtained from Sigma-Aldrich. All other chemicals used were obtained in the highest available purity from CombiBlocks or IrisBiotech. All solvents used were of analytical grade and purchased from Fisher Scientific. Manual solid-phase peptide synthesis was performed in polypropylene syringes equipped with fritted disks, purchased from Torviq.

### Compound analysis and purification

Analytical ultra-performance liquid chromatography (UPLC) analyses were performed on a C18 Agilent InfinityLab Poroshell 120 column (2.7 μm, 100 × 3.0 mm) using an Agilent 1260 Infinity II series system equipped with a diode array UV detector. A gradient with eluent A (water–MeCN– TFA, 95:5:0.1, v/v/v) and eluent B (0.1% TFA in MeCN) rising linearly from 0 to 50% of B over 10.0 min at a flow rate of 1.2 mL min^−1^ was applied to determine the purity of peptides (λ = 215 nm). UPLC-mass spectrometry (MS) analyses were performed on a Phenomenex Kinetex column (1.7 m, 100 Å, 50 × 2.10 mm) using a Waters Acquity system. A gradient with eluent C (0.1% HCOOH in water) and eluent D (0.1% HCOOH in MeCN) rising linearly from 0 to 95% of D over 5.20 min at a flow rate of 0.6 mL min^−1^ was applied to analyse reaction mixtures. Preparative high-performance liquid chromatography (HPLC) purification was performed on a C18 Phenomenex Luna column (5 m, 100 Å, 250 × 20 mm) using an Agilent 1260 LC system equipped with a diode array ultraviolet detector. Various gradients with eluent A and eluent B at a flow rate of 20 mL min^−1^ were applied for the purification. Fractions containing the purified target peptide were identified using UPLC-MS or matrix assisted laser desorption ionisation–time of flight mass spectrometry (MALDI-TOF MS). MALDI-TOF mass spectra were recorded with a Bruker microflex bench-top MALDI using a matrix of alpha-cyano-4-hydroxycinnamic acid or 2,5-dihydroxybenzoic acid in water/MeCN (1:1, v/v) containing 0.1% TFA. The observed *m/z* corresponded to the monoisotopic ions, unless otherwise stated. Nuclear magnetic resonance (NMR) spectra were recorded at 298 K using a Bruker Avance III HD (^1^H NMR and ^13^C NMR recorded at 600 MHz and 150 MHz, respectively). Chemical shifts are reported in parts per million (ppm) relative to the deuterated solvent peak of DMSO-*d*_6_ (δ_H_ = 2.50 ppm; δ_C_ = 39.52 ppm) or CDCl_3_ (δ_H_ = 7.26 ppm; δ_C_ = 77.16 ppm) as internal standard.

### NCL trapping of AIPs from bacterial supernatants

Bacterial isolates were streaked on agar plates and grown over night at 37 °C. Single colonies were then inoculated in 50 mL TSB media overnight at 37 °C in an incubator at 200 rpm shaking. Overnight cultures were centrifuged at 8000 rpm at 4 °C and supernatants filtered through a sterile filter (0.22 μm) and stored at 4 °C for direct use or frozen and stored at −20 °C until use. *Resin preparation:* Fmoc-Cys(S*t*-Bu)-Rink-PEGA resin (50 mg) or Fmoc-Cys(STmp)-Rink-PEGA resin (50 mg) was placed in a 2.0 mL polypropylene syringe equipped with a fritted disk, swelled in DMF for 15 min and washed with DMF (5 × 1 min). The resin was treated with piperidine in DMF (1:4, v/v, 2.0 mL) (1 × 2 min, 1 × 20 min) and washed with DMF (5 × 1 min). The resin was then treated with a solution of *β*-mercatoethanol (BME) in DMF (1:4, v/v, 2.0 mL) containing *N*-methyl morpholine (NMM) (0.1 M) or DL-dithiothreitol (DTT) in DMF (0.05:0.95, w/v, 2.0 mL) containing NMM (0.1 M) (3 10 min) and subsequently washed with DMF (3 × 1 min), MeOH (3 × 1 min) and H_2_O (3 × 1 min). *NCL trapping:* The sterile and filtered bacterial supernatant (~50 mL) was added to 50 mL centrifugal tube and the pH adjusted to pH = ~7.0 using aqueous NaOH (1.0 M). An aqueous tris(2-carboxyethyl)phosphine hydrochloride (TCEP) solution (1.0 mL, 0.5 M, pH = 7.0; final conc. = 10.0 mM) was added to the supernatant followed by the deprotected Cys-Rink-PEGA-resin and the centrifugal tube containing the trapping mixture was agitated at 37 °C overnight. The next day, the resin was separated from the supernatant through filtration using a 10 mL polypropylene syringe equipped with a fritted disk under suction and washed with DMF (3 × 1 min), H_2_O (3 × 1 min), and DMF (3 × 1 min). A solution of DTT in DMF (0.05:0.95, w/v, 2.0 mL) containing NMM (0.1 M) was added to the resin and the resin was agitated at 37 °C. After 30 min, the resin was washed with DMF (3 1 min), MeOH (3 × 1 min), and CH_2_Cl_2_ (3 × 1 min) and dried under suction for 15 min. The dried resin was treated with a cleavage cocktail (2.0 mL, TFA–MilliQ water, 97:3, v/v) for 2 h at room temperature. The peptide containing cleavage solution was removed from the resin, collected and the resin rinsed with neat TFA (1.0 mL). The combined TFA fractions were evaporated under N_2_ stream to near dryness, redissolved in a solution of MeCN in H_2_O (100 μL, 1:1, v/v) and filtered (0.22 μm). *LC-MS analysis*: The filtered TFA cleavage solution was analysed using an Waters Acquity system equipped with a Phenomenex Kinetex column (1.7 μm, 100 Å, 50 2.10 mm) applying a gradient with eluent C (0.1% HCOOH in water) and eluent D (0.1% HCOOH in MeCN) rising linearly from 0 to 50% of D over 10.0 min at a flow rate of 0.6 mL min^1^ and an injection volume of 40 μL. The total ion chromatograms (TIC) were analyzed by displaying extracted ion chromatograms (EIC) of *m/z* [M+H]^+^ values of the possible linear peptides with an additional *C*-terminal cysteine and amide functionality based on the AgrD sequence.

### Concentration determination of DMSO stock solutions

Concentrations of DMSO stock solutions for compounds were determined by quantitative NMR relative to the signal (H = 6.24 ppm, 2H) of the internal standard maleic acid (qNMR grade).

### *β*-Lactamase assay for IC_50_ determination against *S. aureus agr*-I–IV

The *β*-lactamase reporter strains RN10829 [(P2-*agrA*:P3-*blaZ*)]^42^, with p*agrC-*I^43^, p*agrC-*II, p*agrC-* III or p*agrC-*IV^34^ substituting the native *agr* locus with a chromosomal integration of P2-*agrA* and P3-*blaZ* and a plasmid from which a wild-type variant of the corresponding AgrC is expressed, were used to asses inhibition and activation of the AgrC receptor via *β*-lactamase activity in response to varying concentrations of the QS modulating peptides. Overnight cultures of the reporter strains in TSB medium were diluted 1:250 in fresh TSB medium and grown to OD_600_ = 0.35–0.40 (early exponential phase) at 37 °C. Peptide solutions (10 μL) in 1:10 serial dilutions from DMSO stock solutions (1 mM) in TSB medium (final concentrations = 10 μM–10 pM) were added to each well of a clear 96-well plate as well as solutions (10 μL) of cognate AIP (final concentration = 100 nM) in TSB medium followed by 80 μL of bacterial cells. Control wells for 100% *β*-lactamase activity were wells replacing peptide solution with TSB media (10 μL) and control wells for 0% *β*-lactamase activity were wells replacing both peptide and cognate AIP solutions with TSB medium (20 μL). The 96-well plates were incubated at 37 °C shaking at 200 rpm for 1 h and immediately frozen down at − 80 °C to minimize growth during nitrocefin treatment. Next, the 96-well plates were thawed and OD_600_ values were determined using a plate-reader followed by addition of 50 μL of nitrocefin solution to the wells (final concentration = 33.3 μg/mL). *β*-Lactamase activity was monitored at OD_486_ every 20 s for 10 min at 37 °C using a plate-reader. Linear nitrocefin conversion rates were plotted to obtain IC_50_ values by non-linear regression with variable slope using GraphPad Prism 8.0 software. Assays were performed at least as duplicate determinations in biological triplicate.

### Fluorescence reporter assay for screening *agr*-interference of AIPs

Peptides were evaluated for the ability to interfere with *agr-*mediated quorum sensing in *S. aureus* (AH1677, AH430, AH1747 and AH1872 for *agr*-I–IV, respectively)^44^ using reporter strains expressing yellow fluorescent protein (YFP) upon *agr* activation. Interference with *agr-*mediated quorum sensing in *S. epidermidis* (AH3408, AH3623 and AH3409 for *agr*-I–III, respectively)^35^ and *S. lugdunensis* (AH4031 for *agr*-I)^30^ was evaluated using reporter strains expressing superfolder green fluorescent protein (sGFP) upon *agr* activation. Overnight cultures of the reporter strains were grown in TSB medium containing chloramphenicol (CAM, 10 μg/mL for *S. aureus*) or erythromycin (ERM, 10 μg/mL of *S. epidermidis* and *S. lugdunensis*) and diluted 1:100 in fresh TSB medium containing the same antibiotic. Assays were performed in sterile black 96-well plates with clear bottom and the outer wells were filled with water (200 μL) to ensure consistent bacterial growth in the remaining 60 wells. All peptides were screened at concentrations of 1.0 μM and 50 nM and peptides showing at least 75% inhibition at 50 nM were screened further at 2.5 nM against the respective reporter strain, and similarly at 0.125 nM in the case of at least 75% inhibition at 2.5 nM. DMSO stock solutions of peptides (1 mM) were diluted in TSB media and added (15 μL) in technical triplicate to the 96-well plate followed by diluted bacterial overnight cultures (135 μL). Control wells for 100% *agr* acitivty were wells replacing the peptide solution with TSB medium (15 μL). Wells containing 150 μL TSB medium were used to measure background fluorescence. The 96-well plates were incubated on a shaking table at 400 rpm at 37 °C for 22–24 h and fluorescence (for GFP: extication 479 nm, emission 520 nm; for YFP: extication 500 nm, emission 541 nm; automatic gain) and OD_600_ values were subsequently measured using a plate-reader. Background fluorescence was subtracted from all wells and further normalized to the corresponding OD_600_ value of the respective wells. Average fluorescence of control wells was used as relative measure for 100% activation of the *agr*-circuit and bar graphs were generated using GraphPad Prism 8.0 software. All assays were performed in biological triplicate.

### Fluorescence reporter assay for IC_50_ determination

Overnight cultures of the reporter strains were grown in TSB medium containing chloramphenicol (CAM, 10 μg/mL for *S. aureus*) or erythromycin (ERM, 10 μg/mL of *S. epidermidis* and *S. lugdunensis*) and diluted 1:100 in fresh TSB medium containing the same anitbiotic. Assays were performed in sterile black 96-well plates with clear bottom. Peptide solutions (15 μL) in 1:5 serial dilutions from DMSO stock solutions (1 mM) in TSB medium were added to the 96-well plate in technical duplicate followed diluted bacterial overnight cultures (135 μL). Control wells for 100% *agr* acitivty were wells replacing the peptide solution with TSB medium (15 μL). Wells containing 150 μL TSB medium were used to measure background fluorescence. The 96-well plates were incubated in a humidified incubator at 37°C shaking at 1000 rpm for 22–24 h and fluorescence (for GFP: extication 479 nm, emission 520 nm; for YFP: extication 500 nm, emission 541 nm; automatic gain) and OD_600_ values were subsequently measured using a plate-reader. Background fluorescence was subtracted from all wells and further normalized to the corresponding OD_600_ value of the respective wells. Average fluorescence of control wells were used as relative measure for 100% activation of the *agr*-circuit. Relative *agr* activity was plotted to obtain IC_50_ values by non-linear regression with variable slope using GraphPad Prism 8.4 software. All assays were performed in biological triplicate.

### Maximum-likelihood tree construction for AgrD sequences

AgrD sequences (125) were retrieved from the protein database of the National Center of Biotechnology Information (NCBI). Multiple sequence alignment was perfomed using the program ClustalW^45^ in MEGA X (10.1.18) and maximum-likelihood trees were constructed through 500 bootstrap replicates using the Jones-Taylor-Thornton (JTT) model^46^ in MEGA X (10.1.18). Cladograms were visualized using the program FigTree (v1.4.3).

### In vivo MRSA skin infection model

The mouse model was performed under contract at Statens Serum Institut (DK) essentially as previously described^47^. In brief, eight to ten-week-old Balb/c female mice (Taconic Denmark) mice were used for all experiments [n = 16 for vehicle, n = 8 for Fucidin^®^ (2% fusidic acid ointment) treatment, and n = 8 for *S. simulans* AIP-II (**24**) treatment]. All animal experiments were approved by the National Committee of Animal Ethics, Denmark. Mice were anaesthetized and the hair was removed on a 2 cm^2^ skin area on the back and thereafter was the outer most layer of the skin scraped off with a dermal curette to obtain a 1 cm^2^ superficial skin lesion. For vehicle control and Fucidin^®^ treatment, 10 μL inoculum containing approximatly 10^7^ CFU of methilicin-resistant *S. aureus* (MRSA43484) were spread on the skin lesions. For *S. simulans* AIP-II (**24**) treatment, 10 μL of the same inoculum containing **24** (100 μM prepared from a 10 mM DMSO stock solution of **24.**HCl immediately before application) were spread on the skin lessions. After the applied inoculum had dried, the mice were placed in a cage and kept in a warming cabinet until fully awake. The topical skin treatment with Fucidin^®^ was initiated one day after inoculation (day 2, 3, 4) by spreading 50 μL of Fucidin^®^ on the inoculated skin area once a day. Skin lesion size and body weight was measured on day 1, 2, and 4 of all mice. Mice were sacrificed on day 1 (n = 8 vehicle), day 2 (n = 4 vehilce, Fucidin^®^, **24**) and day 4 (n = 4 vehilce, Fucidin^®^, **24**) and the infected skin area was cut out and homogenized to determine the CFU count in the skin lesions.

## Statistical analysis

All statistical analyses were performed using GraphPad Prism 8.4 software. *P* values were determined using one-way analysis of variance (ANOVA) and Dunnet’s test. *P* values < 0.05 considered significant.

## Acknowledgements

We thank Prof. Horswill (Univerity of Colorado) for his donation of fluorescent reporter strains. We thank Carina Vingsbo Lundberg and Karen Juhl from Statens Serum Institut (DK) for performing the mouse studies under contract. We thank Peter Damborg from Statens Serum Institut (DK) for contributing bacterial isolates. This work was supported by the Danish Independent Research Council–Natural Sciences (Grant No. 0135-00427B; C.A.O.) and the LEO Foundation Open Competition Grant (LF-OC-19-000039; CAO).

## Author contributions

C.A.O. and B.H.G. conceptualised the study; B.H.G., B.S.B., L.V., L.M., and M.S.B. performed the experiments; P.S.A. contributed nasal swap samples and bacterial strains; C.A.O. P.S.A and H.I. supervised the study; B.H.G. and C.A.O. wrote the manuscript with input from all authors; C.A.O. acquired funding.

## Additional information

**Supplementary information** is available for this paper.

**Correspondence and requests for materials** should be addressed to C.A.O.

**Extended Data Fig. 1.**
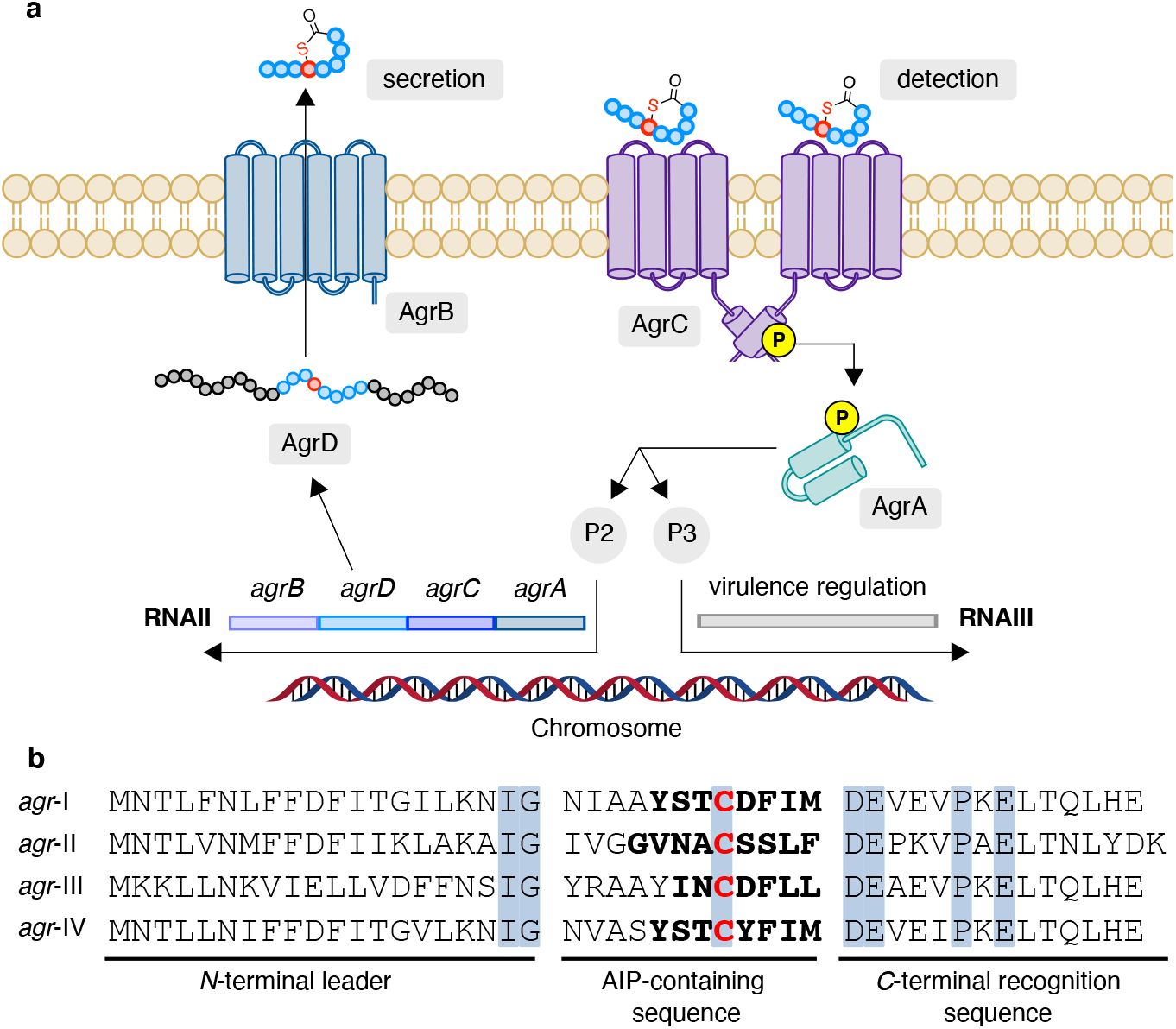
The staphylococcal *accessory gene regulator* (*agr*) system. **a**, The chromosomal *agr* locus consists of the two promoter regions P2 and P3, which initiate the transcription of RNAII and RNAIII, respectively. RNAII encodes the four protein components (AgrA, AgrB, AgrC, and AgrD) of the *agr* system, while RNAIII is a regulatory RNA controlling the expression of virulence factors. In the first step of the *agr* circuit, the AIP-precursor peptide AgrD (44–46 amino acids for SA) is processed by the membrane-embedded endopeptidase AgrB installing the thiolactone functionality. The peptide is further translocated to the extracellular space and finally cleaved to release the mature AIP. Once a concentration threshold of the AIP molecules is reached due to increased cell density, the AgrC receptor, a homodimeric membrane-bound histidine kinase, is activated. AIP-induced activation is followed by auto-phosphorylation of AgrC and subsequent phosphoryl transfer to AgrA, the response regulator of the *agr* system, making the AgrC–AgrA interaction a classical two-component regulatory system. Phosphorylated AgrA binds to the promotors P2 and P3, resulting in upregulated transcription of RNAII and RNAIII, which lead to a positive feedback loop for AgrBDCA expression as well as upregulated expression of virulence factors. **b**, AgrD peptides consist of three domains, the *C*-terminal recognition sequence, the *N*-terminal leader peptide and in between the 12 amino acid long AIP-containing sequence. Conserved residues are highlighted in blue boxes.

**Extended Data Fig. 2.**
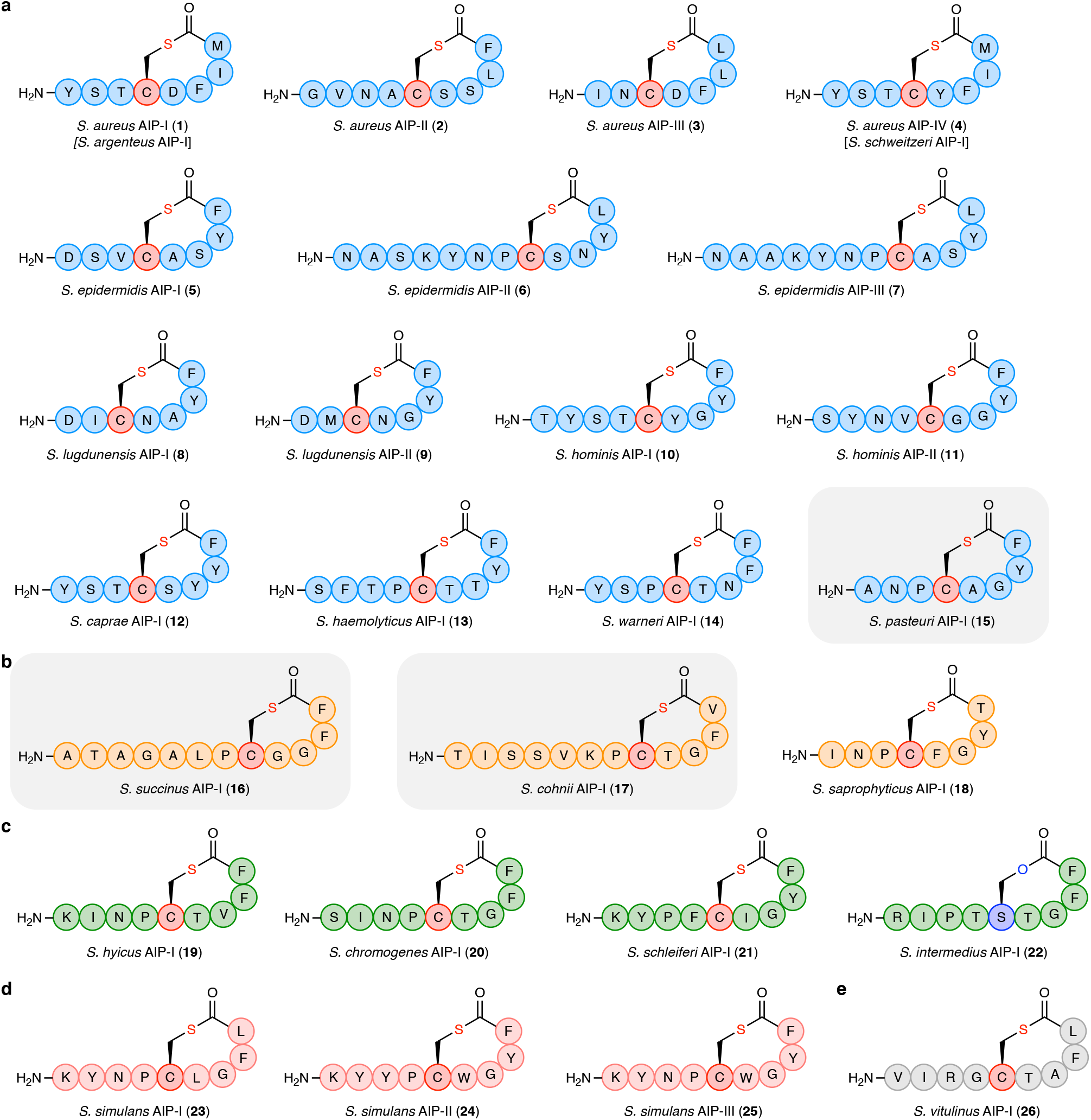
Structures of the 26 identified AIPs studied herein. **a**, 15 (17) AIPs known for the **Epidermidis-Aureus** species group [*S. aureus* AIP-I–IV (**1**–**4**), *S. epidermidis* AIP-I–III (**5**–**7**), *S. lugdunensis* AIP-I–II (**8**–**9**), *S. hominis* AIP-I–II (**10**–**11**), *S. caprae* AIP-I (**12**), *S. haemolyticus* AIP-I (**13**), *S. warneri* AIP-I (**14**), *S. pasteuri* AIP-I (**15**)]. **b**, 3 AIPs known from the **Saprophyticus** species group [*S. succinus* AIP-I (**16**), *S. cohnii* AIP-I (**17**), *S. saprophyticus* AIP-I (**18**)]. **c**, 4 AIPs known from the **Hyicus-Intermedius** species group [*S. hyicus* AIP-I (**19**), *S. chromogenes* AIP-I (**20**), *S. schleiferi* AIP-I (**21**), *S. intermedius* AIP-I (**22**)]. **d**, 3 AIPs known from the **Simulans** species group [*S. simulans* AIP-I–III (**23**–**25**)]. **e**, 1 AIP known from the **Sciuri** species group [*S. vitulinus* AIP-I (**26**)]; and no AIPs have been identified from **Auricularis**. AIPs highlighted in grey boxes were newly identified in this study.

**Extended Data Fig. 3.**
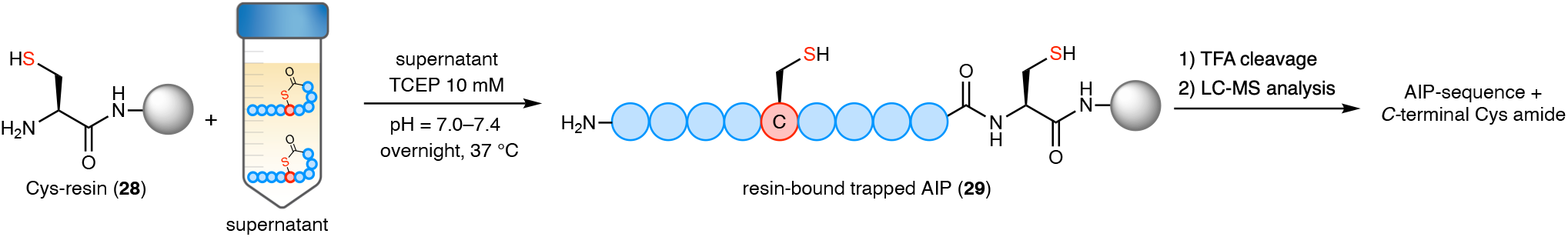
Identification of AIPs through native chemical ligation (NCL) trapping. Cys-resin (**28**) is incubated overnight in pH-adjusted bacterial supernatant containing tris(2-carboxyethyl)phosphine (TCEP, 10 mM) enabling chemoselective trapping of AIPs. The resin with the trapped AIP (**29**) is washed extensively and subsequently treated with trifluoroacetic acid (TFA) to release the trapped AIP. The concentrated cleavage solution is analyzed by liquid-chromatography mass-spectrometry (LC-MS) for the 7 possible AIPs sequences with an additional *C*-terminal Cys amide extracted from the AIP-precursor peptide AgrD.

**Extended Data Fig. 4.**
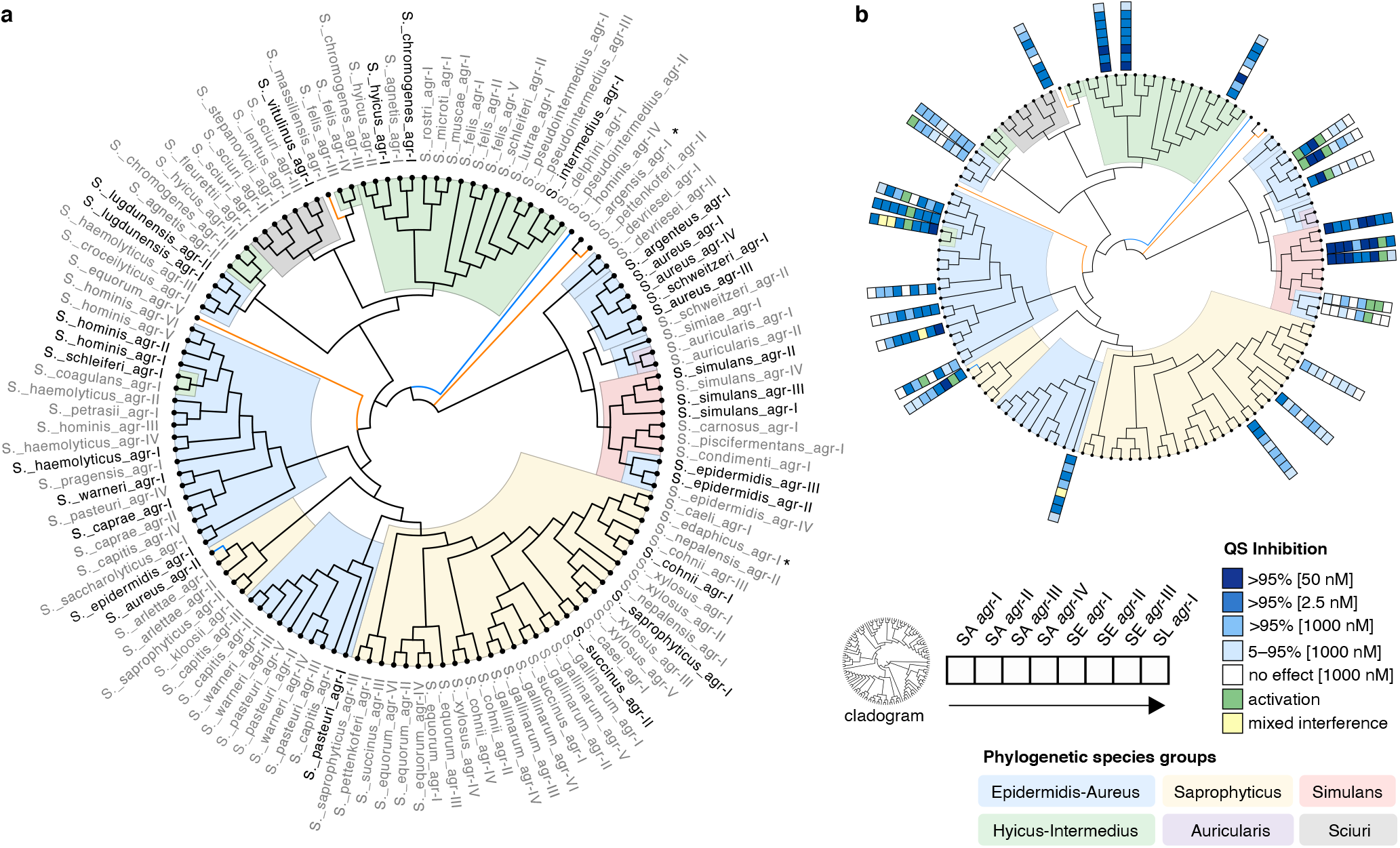
Phylogeny based on the 12 amino acid AIP-containing sequences. Maximum likelihood tree (bootstrap n = 500) created from 125 AIP-containing sequences extracted from AgrD sequences. The *agr* groups of identified AIPs were named in their chronological order of their discovery and *agr* groups with so far unknown AIPs were ranked in order of their number of accessions in the Protein Database. Names of *agr* groups with known AIPs are black and grey with unknown AIPs. Phylogenetic species groups as defined by Lamers *et al.*^36^ are colored as follows: Epidermidis-Aureus, blue; Saprophyticus, tan; Simulans, red; Hyicus-Intermedius, green; Auicularis, purple; Sciuri, gray. *Species not yet assigned to species group. **b**, Cladogram overlaid with QS interactions of known AIPs against SA *agr*-I–IV, SE *agr*-I–III and SL *agr*-I. The interactions of one AIP are represented as colored boxes with SA *agr*-I closest to the cladogram as depicted in the figure.

**Extended Data Table 1.** IC_50_ and EC_50_ values (nM) measured in *β*-lactamase assay using *S. aureus* (SA) AgrC-I–IV reporter strains. All inhibition assays were performed in the presence of 100 nM cognate AIP and the results represent means ± standard error of the mean (SEM) of at least duplicate determinations performed in biological triplicate. a Only partial inhibition observed. ^b^No inhibition recorded at the highest AIP concentration tested. Blue shaded boxes represent IC_50_ values determined in this study. White represent IC_50_/EC_50_ values determined previously^34^. The AIPs of SE (**5**–**7**) and SL-AIP-I (**8**) were weak inhibitors or displayed no significant inhibition at the highest tested concentration. The *S. hominis* AIP-II (**11**) was a moderate inhibitor of SA *agr*-I and *agr*-III with lower potency against *agr*-II and no effect on *agr*-IV. The *S. caprae* AIP-I (**12**) displayed high potencies against *agr*-I and *agr*-II and moderate potency against *agr*-IV. The AIP was not able to inhibit *agr*-III at higher concentrations but showed inhibtion of *agr* activity at lower nanomolar concentrations. The *S. pasteuri* AIP-I (**15**) and *S. succinus* AIP-I (**16**) exhibited moderate to low potencies against the four reporter strains, while the *S. cohnii* AIP-I (**17**) showed only limited inhibition of *agr*-I. The *S. intermedius* AIP (**22**) exhibited moderate and weak potency against *agr*-I and *agr*-II, respectively. Higher potency was observed against *agr*-III and no effect was seen against *agr*-IV. *S. simulans* AIP-II–III (**24**, **25**) were highly potent inhibitors of *agr*-I and *agr*-II and exhibited slightly decreased potencies against *agr*-III and moderate inhibition against *agr*-IV.

| Species AIP | SA AgrC-I | SA AgrC-II | SA AgrC-III | SA AgrC-IV |
| --- | --- | --- | --- | --- |
| *S. aureus* AIP-I (1) | **EC<sub>50</sub>: 5.4 ± 0.5** | 700 ± 165 | 80 ± 11 | **EC<sub>50</sub>: 790 ± 66** |
| *S. aureus* AIP-II (2) | 110 ± 12 | **EC<sub>50</sub>: 3.7 ± 0.5** | 40 ± 15 | 230 ± 19 |
| *S. aureus* AIP-III (3) | 120 ± 15 | 150 ± 63 | **EC<sub>50</sub>: 11 ± 1** | >1000 |
| *S. aureus* AIP-IV (4) | **EC<sub>50</sub>: 59 ± 2** | 47 ± 9 nM | 9.8 ± 0.2<sup>a</sup> | **EC<sub>50</sub>: 2.6 ± 0.2** |
| *S. epidermidis* AIP-I (5) | >1000 | ..b | >1000 | ..b |
| *S. epidermidis* AIP-II (6) | - | ..b | ..b | ..b |
| *S. epidermidis* AIP-III (7) | >10000 | ..b | >10000 | ..b |
| *S. lugdunensis* AIP-I (8) | >1000 | ..b | >1000 | ..b |
| *S. lugdunensis* AIP-II (9) | >1000 | ..b | ..b | ..b |
| *S. hominis* AIP-I (10) | 134 ± 8 | 580 ± 61 | 220 ± 46<sup>a</sup> | **EC<sub>50</sub>: 54 ± 5** |
| *S. hominis* AIP-II (11) | 224 ± 37 | >1000 | 338 ± 83 | ..b |
| *S. caprae* AIP-I (12) | 5.7 ± 0.8 | 252 ± 44 | ..a | 199 ± 19 |
| *S. haemolyticus* AIP-I (13) | 340 ± 25 | ..b | 340 ± 91 | - |
| *S. warneri* AIP-I (14) | 100 ± 10 | 1000 ± 105 | 460 ± 59 | >1000 |
| *S. pasteurii* AIP-I (15) | 155 ± 34 | ..b | 191 ± 18 | >1000 |
| *S. succinus* AIP-I (16) | 117 ± 26 | 854 ± 65 | >1000 | >1000 |
| *S. cohnii* AIP-I (17) | >10000 | ..b | ..b | ..b |
| *S. saprophyticus* AIP-I (18) | 360 ± 30 | - | ..b | ..b |
| *S. hyicus* AIP-I (19) | 3.3 ± 0.5 | 350 ± 90 | 4.0 ± 0.8 | 180 ± 33 |
| *S. chromogenes* AIP-I (20) | 15 ± 1 | 200 ± 13 | 60 ± 13 | 350 ± 85 |
| *S. schleiferi* AIP-I (21) | 2.8 ± 0.8 | 86 ± 6 | 80 ± 16<sup>a</sup> | **EC<sub>50</sub>: 31 ± 6** |
| *S. intermedius* AIP-I (22) | 87 ± 20 | >1000 | 104 ± 20 | ..b |
| *S. simulans* AIP-I (23) | 8.6 ± 0.5 | 23 ± 2 | 50 ± 11 | 280 ± 66 |
| *S. simulans* AIP-II (24) | 1.9 ± 0.3 | 80 ± 7 | 198 ± 33 | 273 ± 26 |
| *S. simulans* AIP-III (25) | 1.6 ± 0.1 | 60 ± 7 | 103 ± 5 | 351 ± 60 |
| *S. vitulinus* AIP-I (26) | 190 ± 15 | 800 ± 116 | 690 ± 46 | ..b |
| *S. aureus* AIP-III D4A (29) | 3.3 ± 0.7 | 34 ± 1 | 1.6 ± 0.2<sup>a</sup> | 10 ± 0.2 |

**Extended Data Table 2.**
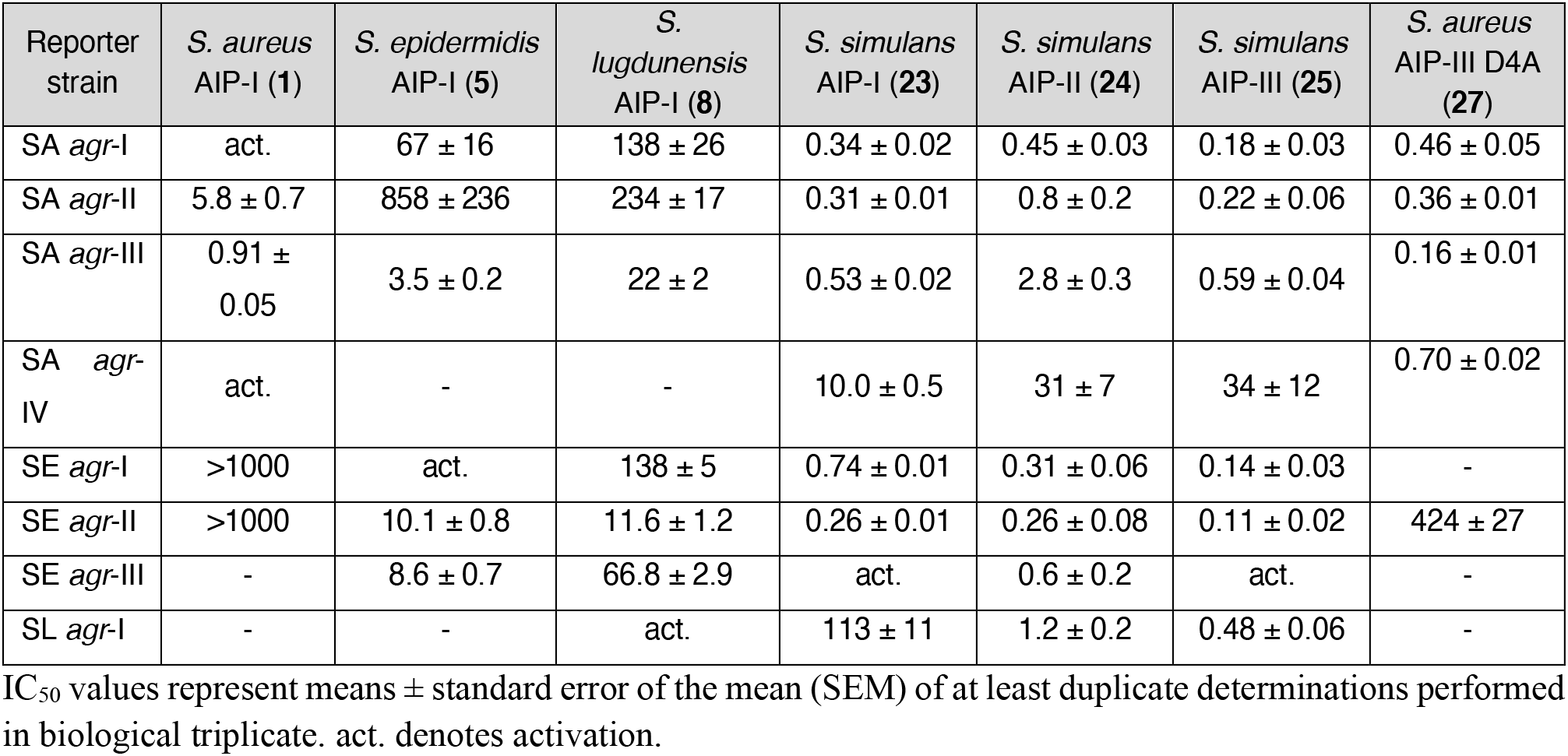
C_50_ values (nM) determined using fluorescent reporter strains of *S. aureus* (SA) *agr*-I–IV, *S. epidermidis* (SE) *agr*-I–III and *S. lugdunensis*(SL) *agr*-I. IC_50_ values represent means ± standard error of the mean (SEM) of at least duplicate determinations performed in biological triplicate. act. denotes activation.

| Reporter strain | *S. aureus* AIP-I (1) | *S. epidermidis* AIP-I (5) | *S. lugdunensis* AIP-I (8) | *S. simulans* AIP-I (23) | *S. simulans* AIP-II (24) | *S. simulans* AIP-III (25) | *S. aureus* AIP-III D4A (27) |
| --- | --- | --- | --- | --- | --- | --- | --- |
| SA *agr*-I | act. | 67 ± 16 | 138 ± 26 | 0.34 ± 0.02 | 0.45 ± 0.03 | 0.18 ± 0.03 | 0.46 ± 0.05 |
| SA *agr*-II | 5.8 ± 0.7 | 858 ± 236 | 234 ± 17 | 0.31 ± 0.01 | 0.8 ± 0.2 | 0.22 ± 0.06 | 0.36 ± 0.01 |
| SA *agr*-III | 0.91 ± 0.05 | 3.5 ± 0.2 | 22 ± 2 | 0.53 ± 0.02 | 2.8 ± 0.3 | 0.59 ± 0.04 | 0.16 ± 0.01 |
| SA *agr*-IV | act. | - | - | 10.0 ± 0.5 | 31 ± 7 | 34 ± 12 | 0.70 ± 0.02 |
| SE *agr*-I | >1000 | act. | 138 ± 5 | 0.74 ± 0.01 | 0.31 ± 0.06 | 0.14 ± 0.03 | - |
| SE *agr*-II | >1000 | 10.1 ± 0.8 | 11.6 ± 1.2 | 0.26 ± 0.01 | 0.26 ± 0.08 | 0.11 ± 0.02 | 424 ± 27 |
| SE *agr*-III | - | 8.6 ± 0.7 | 66.8 ± 2.9 | act. | 0.6 ± 0.2 | act. | - |
| SL *agr*-I | - | - | act. | 113 ± 11 | 1.2 ± 0.2 | 0.48 ± 0.06 | - |

