## Supplemental Information for "Chemical phylogenetics of the staphylococcal quorum sensing landscape"

#### 1. Supplementary figures

##### LC-MS traces of sequence-guided identifications of autoinducing peptides (AIPs)

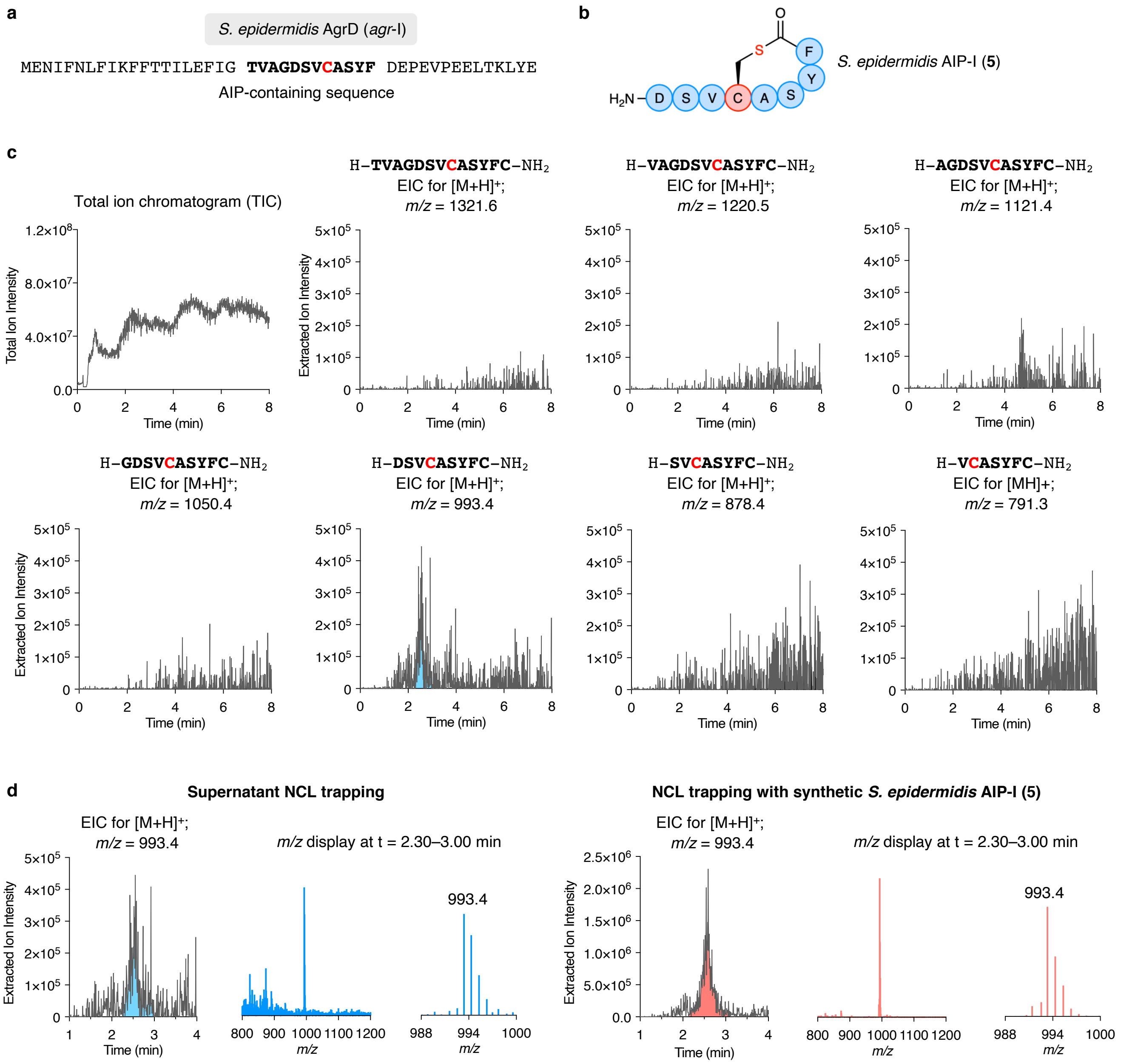

**Supplementary Figure 1. NCL trapping of *S. epidermidis* AIP-I (5). a**, AgrD sequence of *S. epidermidis agr*-I (NCBI: WP_001830021.1). **b**, Structure of *S. epidermidis* AIP-I (**5**). **c**, LC-MS analysis of the TFA cleavage solution: total ion chromatogram (TIC) and extracted ion chromatograms (EIC) of *m/z* = $[M+H]^+$ of possible linear AIP sequences including a C-terminal cysteine amide. **d**, NCL trapping of synthetic *S. epidermidis* AIP-I (**5**) confirms the identity of trapped AIP (**5**) from bacterial supernatant.

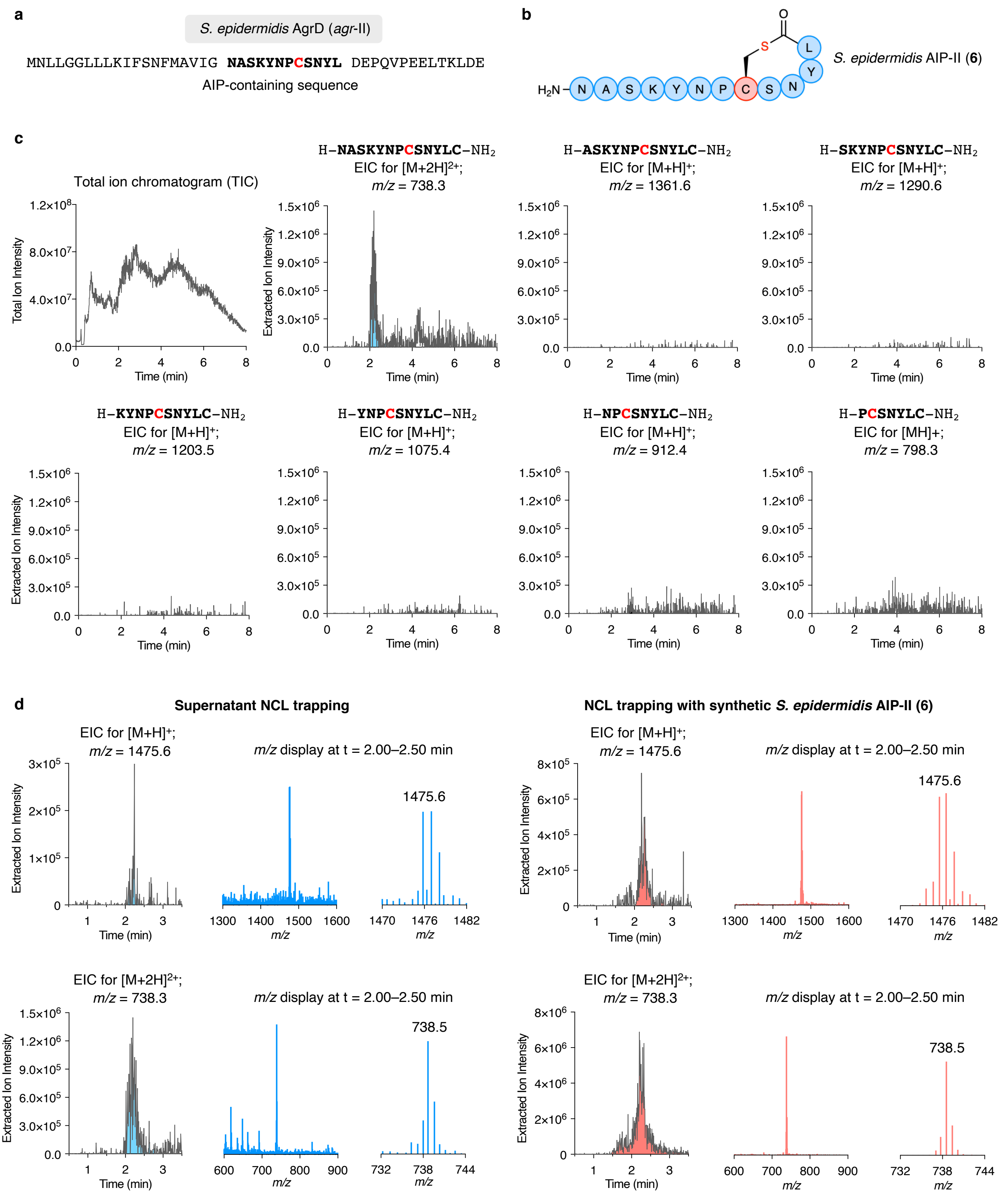

**Supplementary Figure 2. NCL trapping of *S. epidermidis* AIP-II (6). a**, AgrD sequence of *S. epidermidis agr*-II (NCBI: WP_002447513.1). **b**, Structure of *S. epidermidis* AIP-II (**6**). **c**, LC-MS analysis of the TFA cleavage solution: total ion chromatogram (TIC) and extracted ion chromatograms (EIC) of *m/z* = $[M+H]^+$ of possible linear AIP sequences including a C-terminal cysteine amide. **d**, NCL trapping of synthetic *S. epidermidis* AIP-II (**6**) confirms the identity of trapped AIP (**6**) from bacterial supernatant.

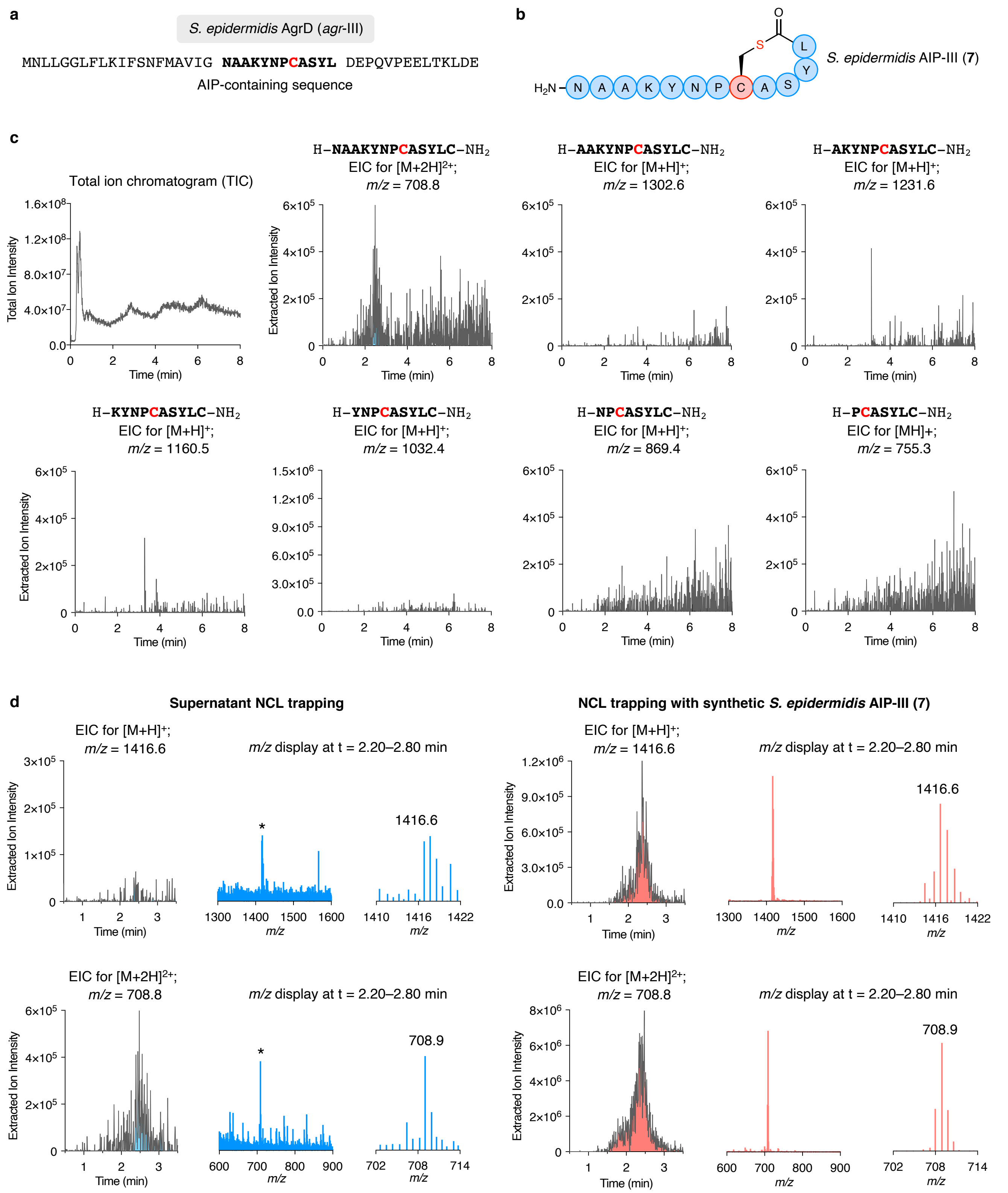

**Supplementary Figure 3. NCL trapping of *S. epidermidis* AIP-III (7). a**, AgrD sequence of *S. epidermidis agr*-III (NCBI: WP_002490348.1). **b**, Structure of *S. epidermidis* AIP-III (**7**). **c**, LC-MS analysis of the TFA cleavage solution: total ion chromatogram (TIC) and extracted ion chromatograms (EIC) of *m/z* = $[M+H]^+$ of possible linear AIP sequences including a C-terminal cysteine amide. **d**, NCL trapping of synthetic *S. epidermidis* AIP-III (**7**) confirms the identity of trapped AIP (**7**) from bacterial supernatant.

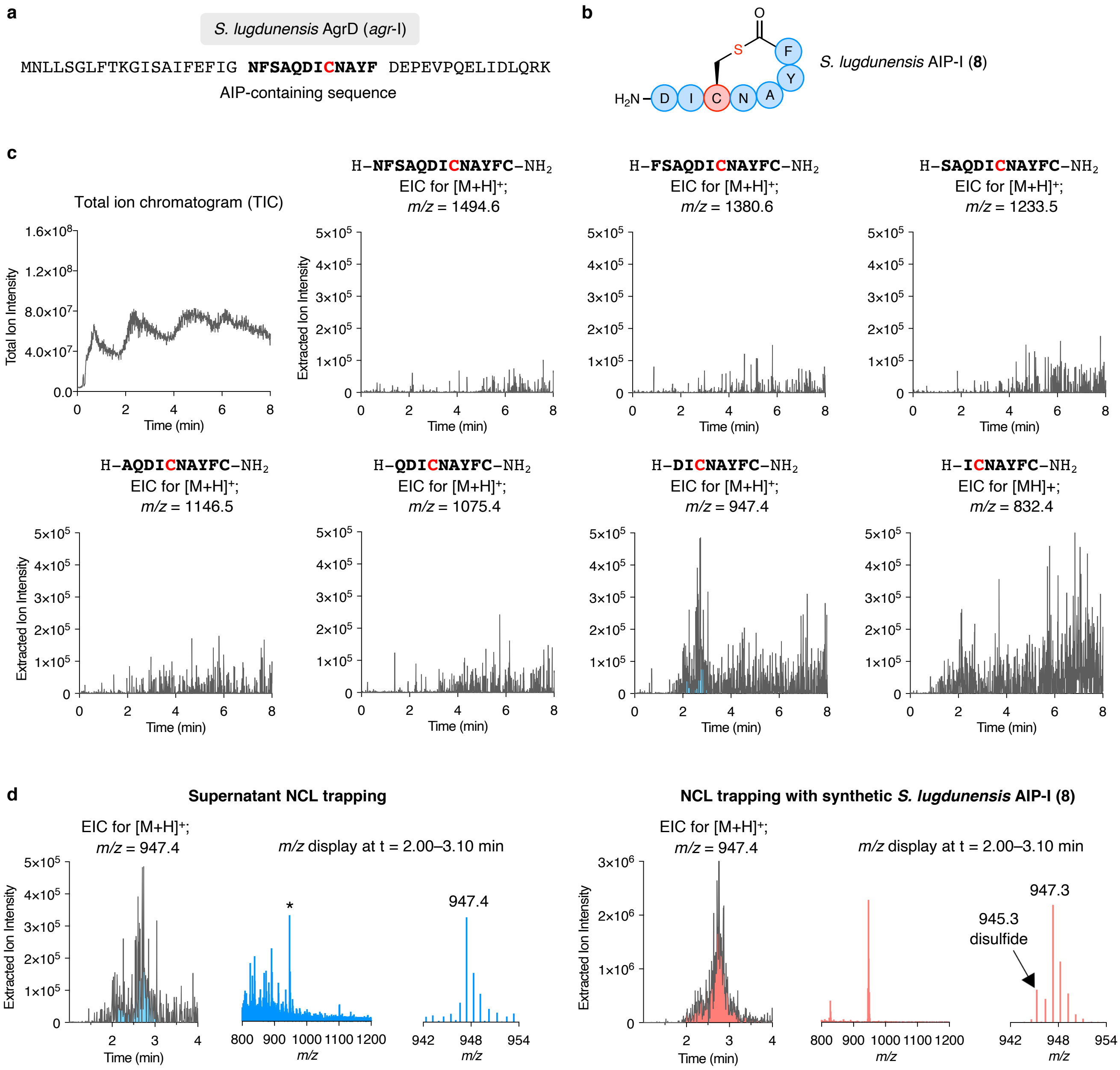

**Supplementary Figure 4. NCL trapping of *S. lugdunensis* AIP-I (8). a**, AgrD sequence of *S. lugdunensis agr*-I (NCBI: WP_002477921.1). **b**, Structure of *S. lugdunensis* AIP-I (**8**). **c**, LC-MS analysis of the TFA cleavage solution: total ion chromatogram (TIC) and extracted ion chromatograms (EIC) of *m/z* = $[M+H]^+$ of possible linear AIP sequences including a C-terminal cysteine amide. **d**, NCL trapping of synthetic *S. lugdunensis* AIP-I (**8**) confirms the identity of trapped AIP (**8**) from bacterial supernatant.

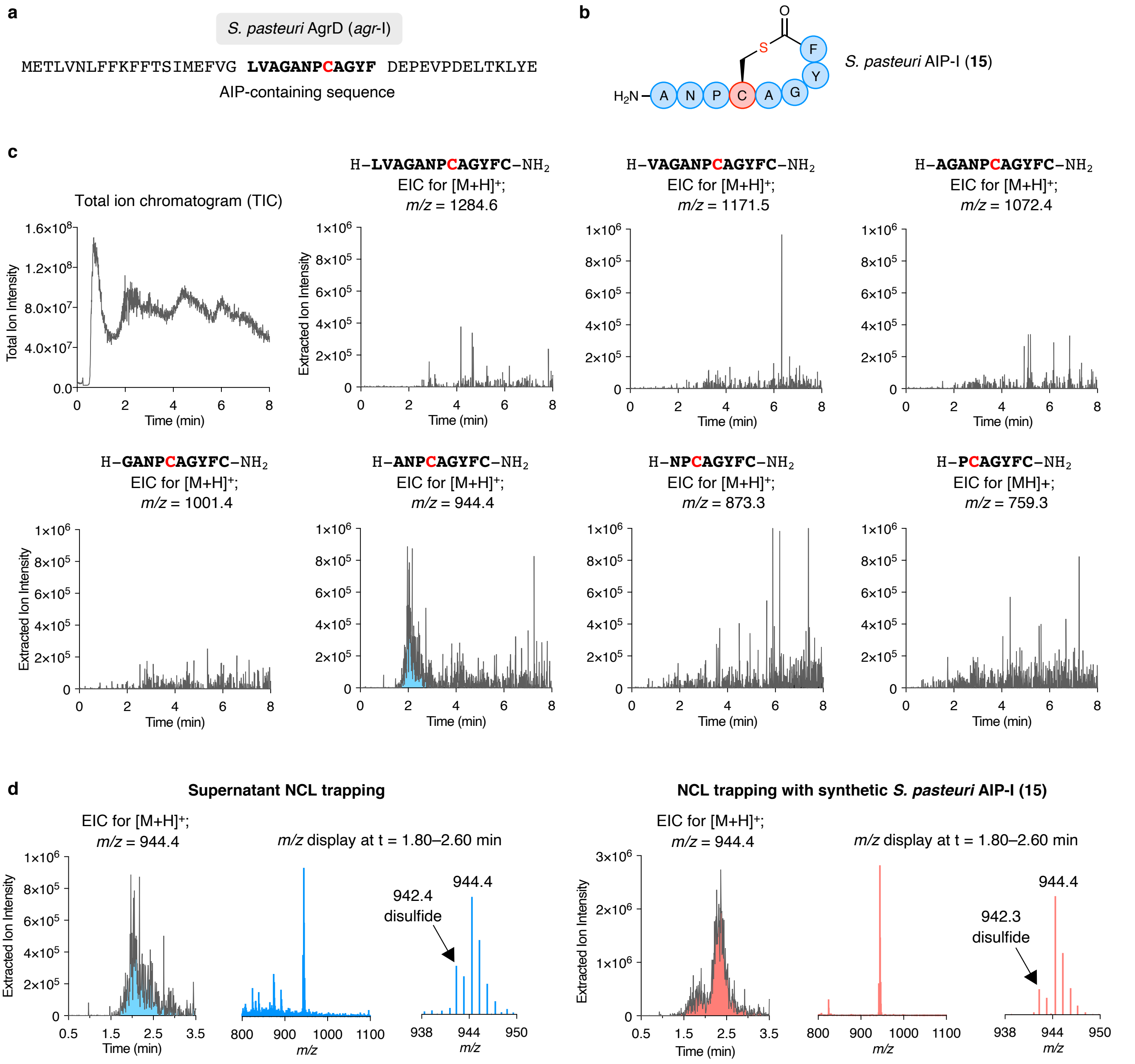

**Supplementary Figure 5. NCL trapping of *S. pasteuri* AIP-I (15). a**, AgrD sequence of *S. pasteuri agr*-I (sequenced in this study). **b**, Structure of *S. pasteuri* AIP-I (**15**). **c**, LC-MS analysis of the TFA cleavage solution: total ion chromatogram (TIC) and extracted ion chromatograms (EIC) of *m/z* = $[M+H]^+$ of possible linear AIP sequences including a C-terminal cysteine amide. **d**, NCL trapping of synthetic *S. pasteuri* AIP-I (**15**) confirms the identity of trapped AIP (**15**) from bacterial supernatant.

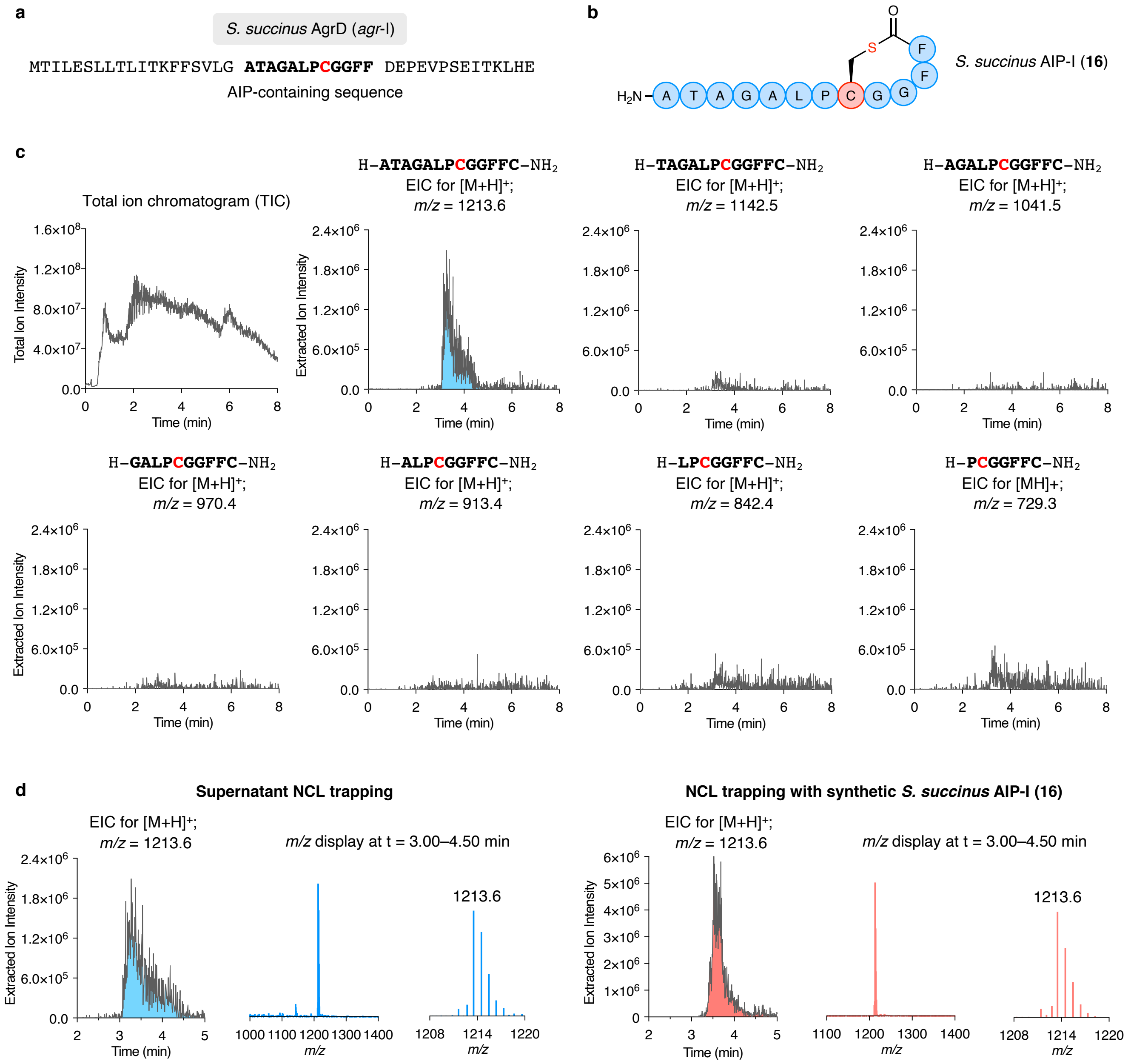

**Supplementary Figure 6. NCL trapping of *S. succinus* AIP-I (16). a**, AgrD sequence of *S. succinus agr*-I (sequenced in this study). **b**, Structure of *S. succinus* AIP-I (**16**). **c**, LC-MS analysis of the TFA cleavage solution: total ion chromatogram (TIC) and extracted ion chromatograms (EIC) of *m/z* = $[M+H]^+$ of possible linear AIP sequences including a C-terminal cysteine amide. **d**, NCL trapping of synthetic *S. succinus* AIP-I (**16**) confirms the identity of trapped AIP (**16**) from bacterial supernatant.

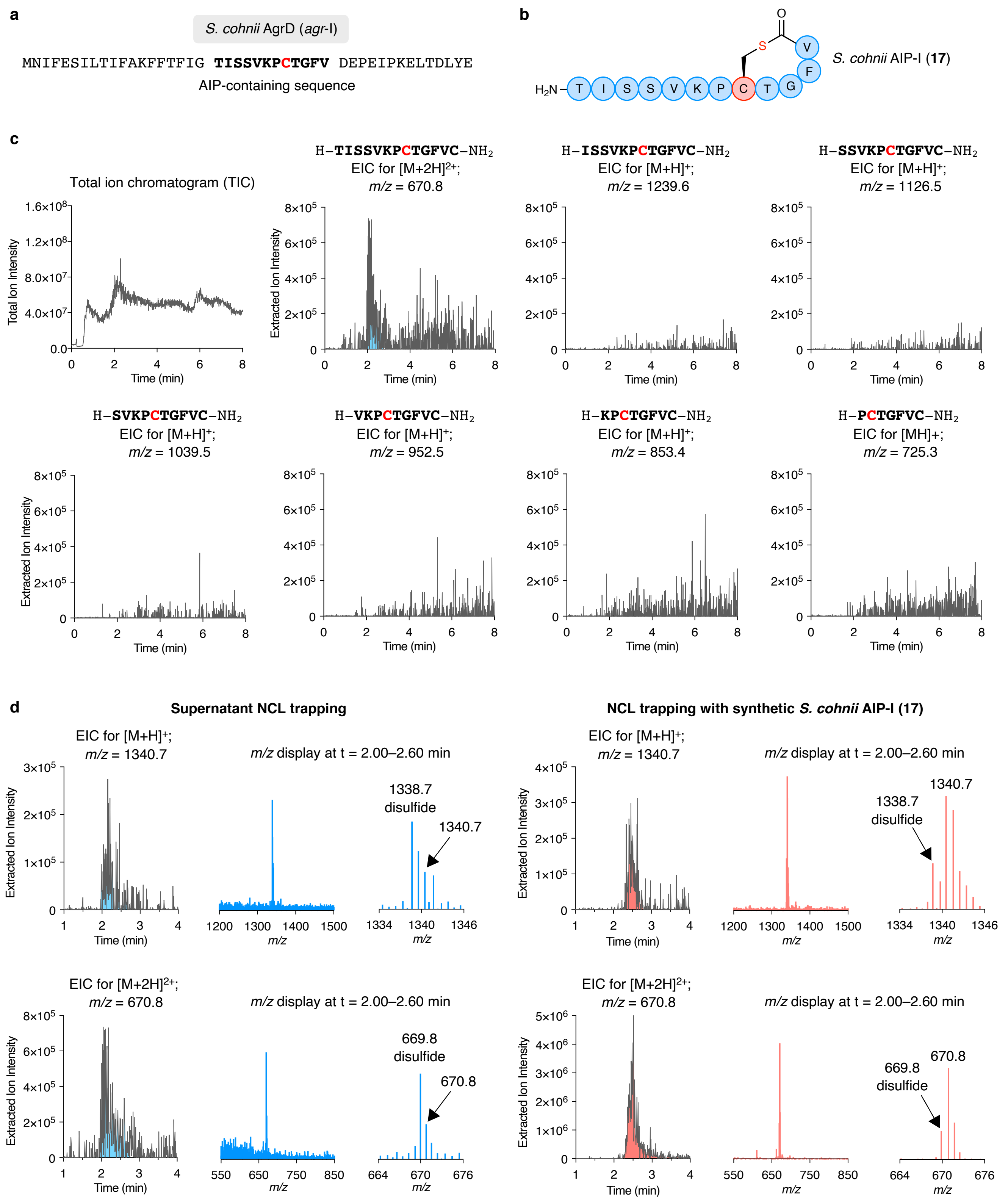

**Supplementary Figure 7. NCL trapping of *S. cohnii* AIP-I (17). a**, AgrD sequence of *S. cohnii agr*-I (sequenced in this study). **b**, Structure of *S. cohnii* AIP-I (**17**). **c**, LC-MS analysis of the TFA cleavage solution: total ion chromatogram (TIC) and extracted ion chromatograms (EIC) of *m/z* = [M+H]$^+$ of possible linear AIP sequences including a C-terminal cysteine amide. **d**, NCL trapping of synthetic *S. cohnii* AIP-I (**17**) confirms the identity of trapped AIP (**17**) from bacterial supernatant.

#### Identification of *agr* groups of *S. epidermidis* isolates from human nasal swabs

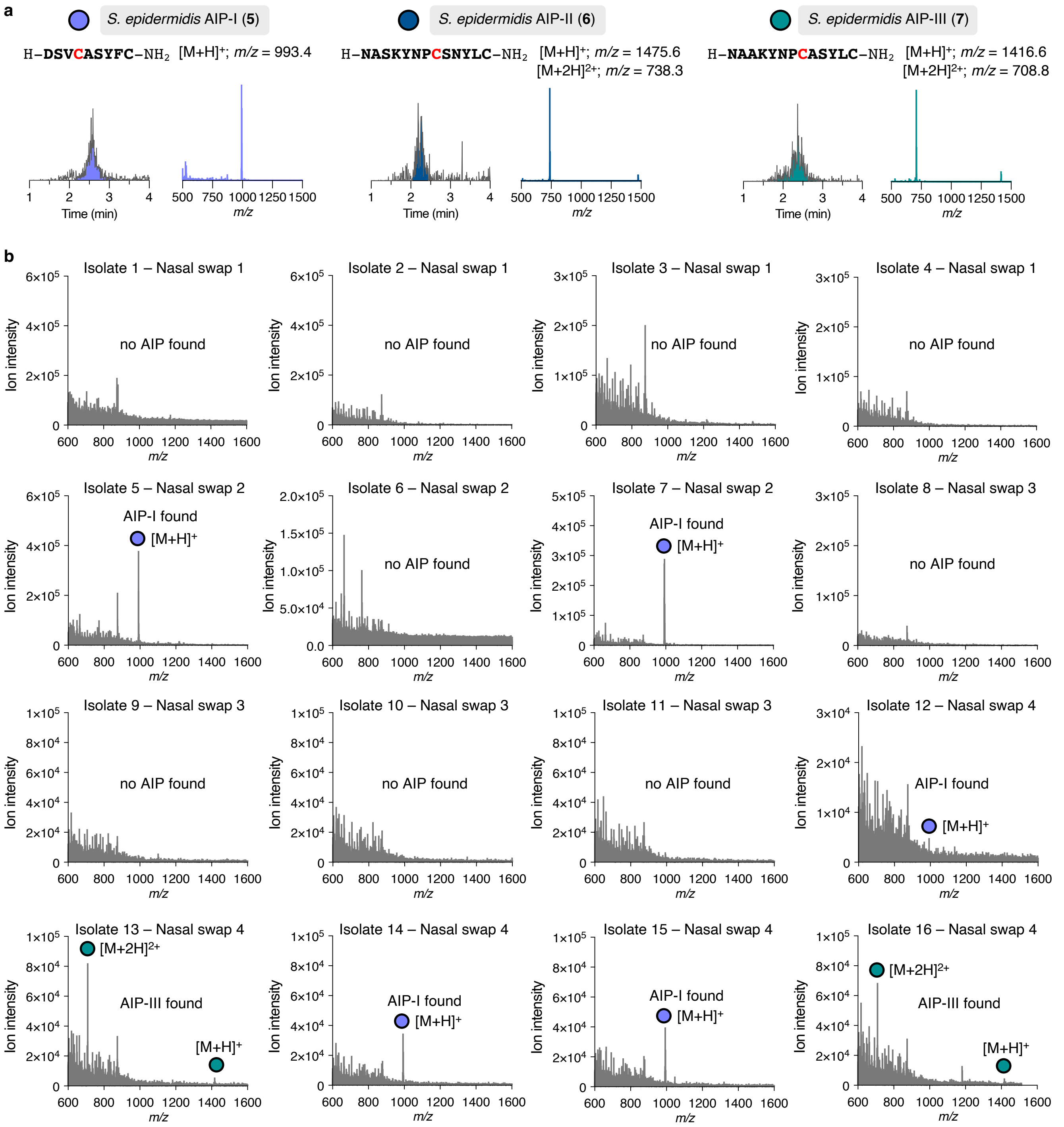

**Supplementary Figure 8. NCL trapping screen of *S. epidermidis* isolates 1–16. a**, NCL trapping of synthetic *S. epidermidis* AIP-I–III (**5**–**7**) with LC-MS retention time ($t_r$ = 2.6 min for **5**; $t_r$ = 2.3 min for **6**; $t_r$ = 2.5 min for **7**) and *m/z* of display. **b**, NCL trapping was performed on supernatants of overnight cultures from nasal swap isolates. The final trapping samples were analyzed by LC-MS through combining the *m/z* display from $t_r$ = 2.00–3.00 min.

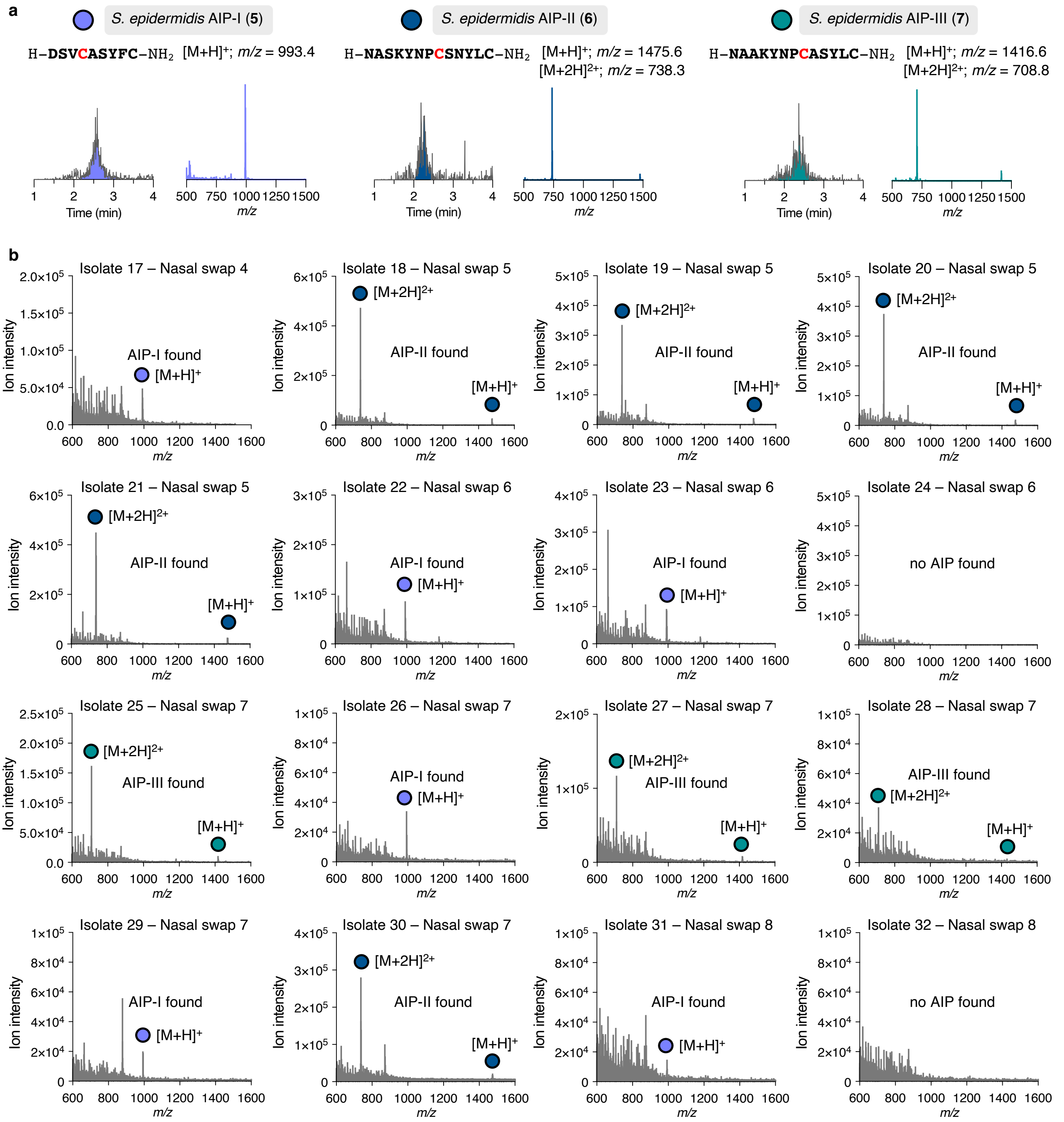

**Supplementary Figure 9. NCL trapping screen of *S. epidermidis* isolates 17–32. a**, NCL trapping of synthetic *S. epidermidis* AIP-I–III (**5**–**7**) with LC-MS retention time ($t_r$ = 2.6 min for **5**; $t_r$ = 2.3 min for **6**; $t_r$ = 2.5 min for **7**) and *m/z* of display. **b**, NCL trapping was performed on supernatants of overnight cultures from nasal swap isolates. The final trapping samples were analyzed by LC-MS through combining the *m/z* display from $t_r$ = 2.00–3.00 min.

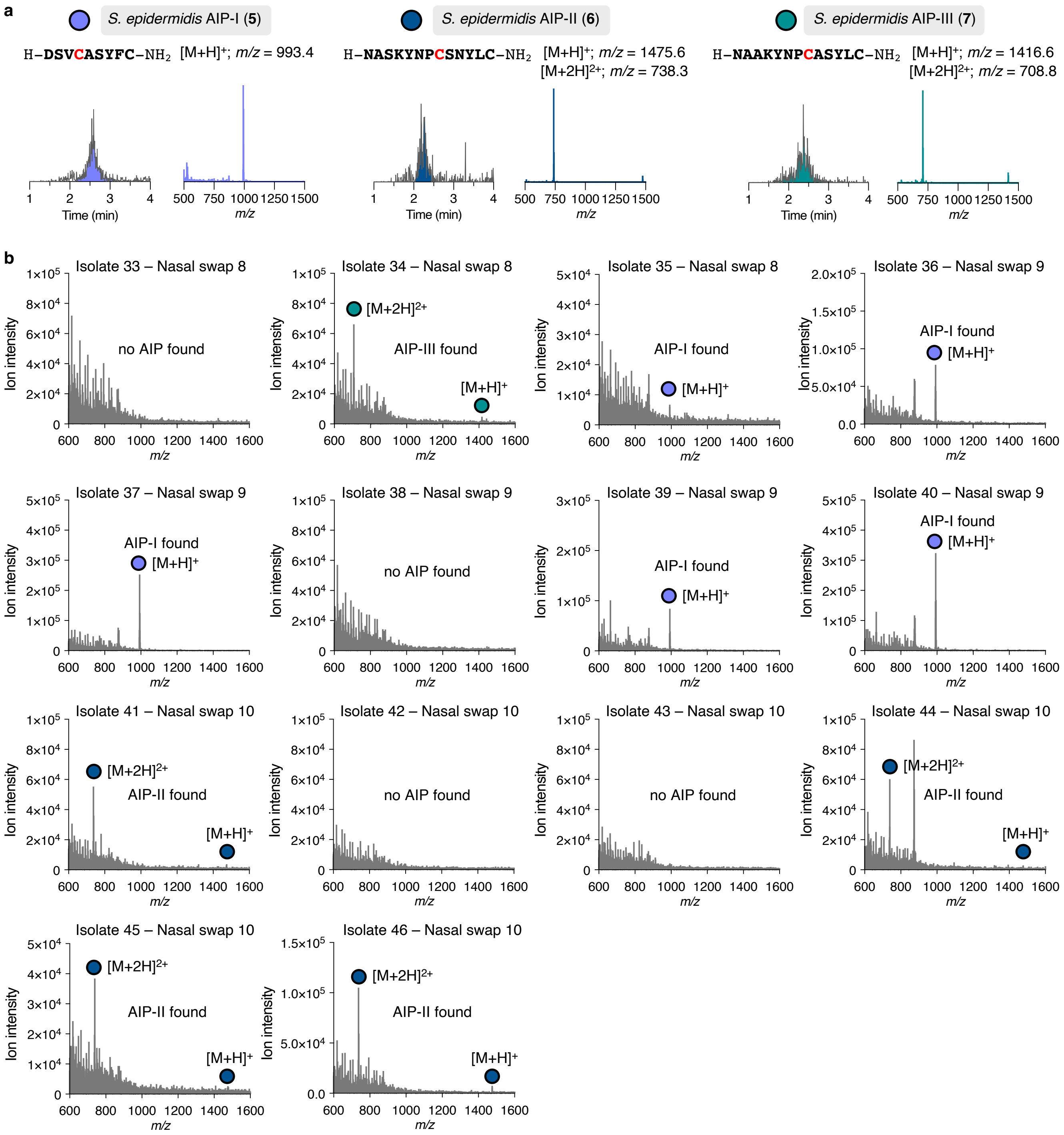

**Supplementary Figure 10. NCL trapping screen of *S. epidermidis* isolates 33–46. a**, NCL trapping of synthetic *S. epidermidis* AIP-I–III (**5**–**7**) with LC-MS retention time ($t_r$ = 2.6 min for **5**; $t_r$ = 2.3 min for **6**; $t_r$ = 2.5 min for **7**) and *m/z* of display. **b**, NCL trapping was performed on supernatants of overnight cultures from nasal swap isolates. The final trapping samples were analyzed by LC-MS through combining the *m/z* display from $t_r$ = 2.00–3.00 min.

#### Dose-response curves for AgrC interference using *β*-lactamase reporter assays

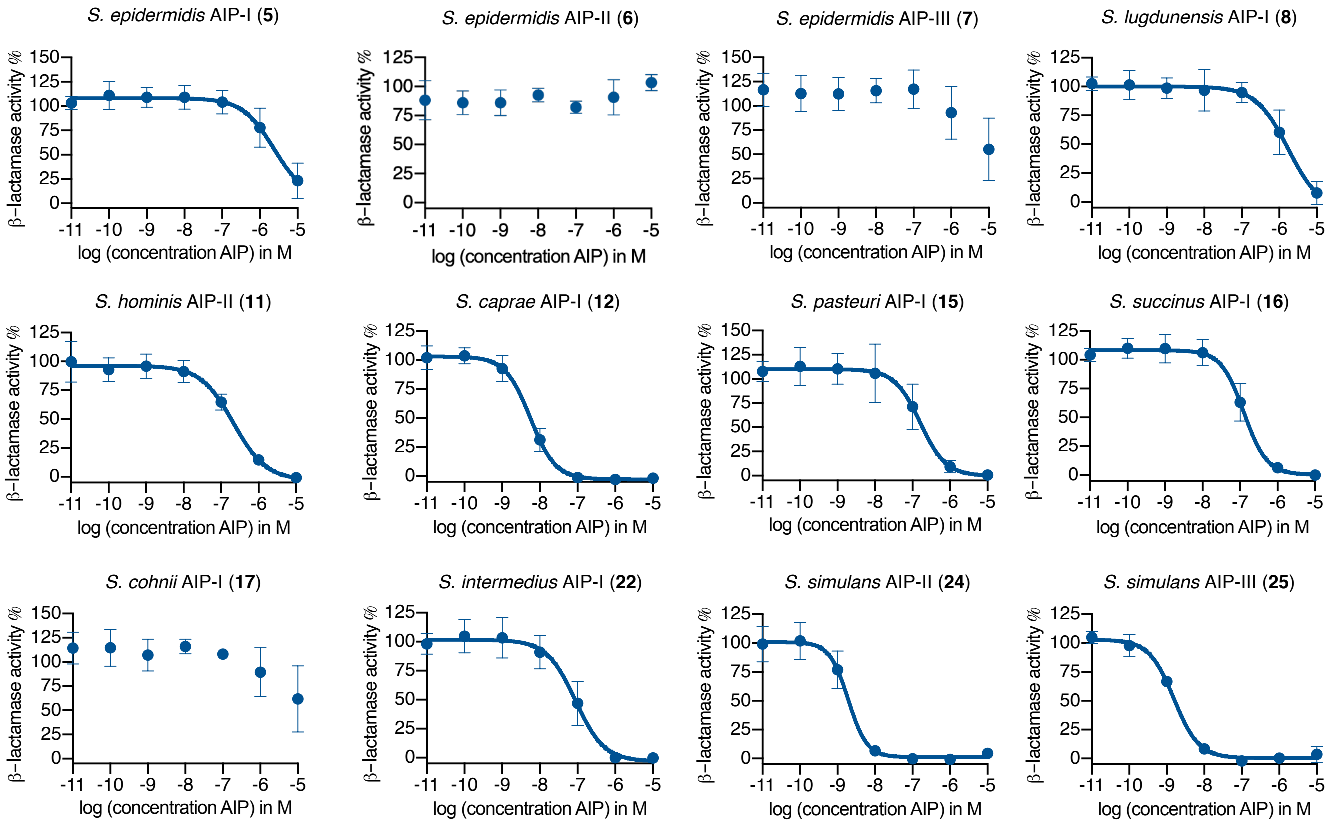

**Supplementary Figure 11. Dose-response curves of AgrC inhibition of *S. aureus agr*-I.** Inhibition properties of synthetic AIPs (10 µM to 10 pM) were determined through *β*-lactamase activity using the *S. aureus agr*-I reporter strain in the presence of 100 nM *S. aureus* AIP-I (**1**). The curves were generated from three individual assays performed in technical duplicate and shown error bars are the standard deviation of the mean (SD).

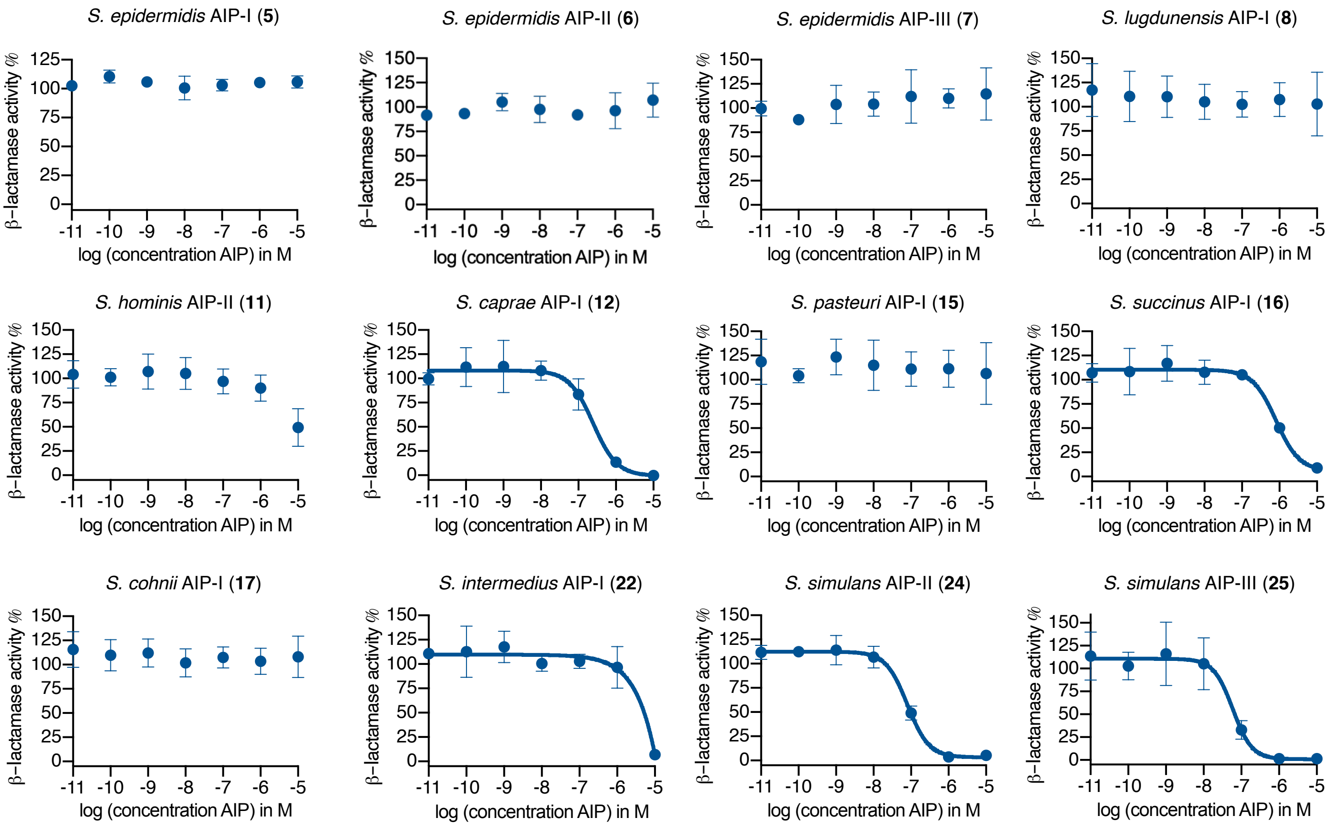

**Supplementary Figure 12. Dose-response curves of AgrC inhibition of *S. aureus agr*-II.** Inhibition properties of synthetic AIPs (10 µM to 10 pM) were determined through *β*-lactamase activity using the *S. aureus agr*-II reporter strain in the presence of 100 nM *S. aureus* AIP-II (**2**). The curves were generated from three individual assays performed in technical duplicate and shown error bars are the standard deviation of the mean (SD).

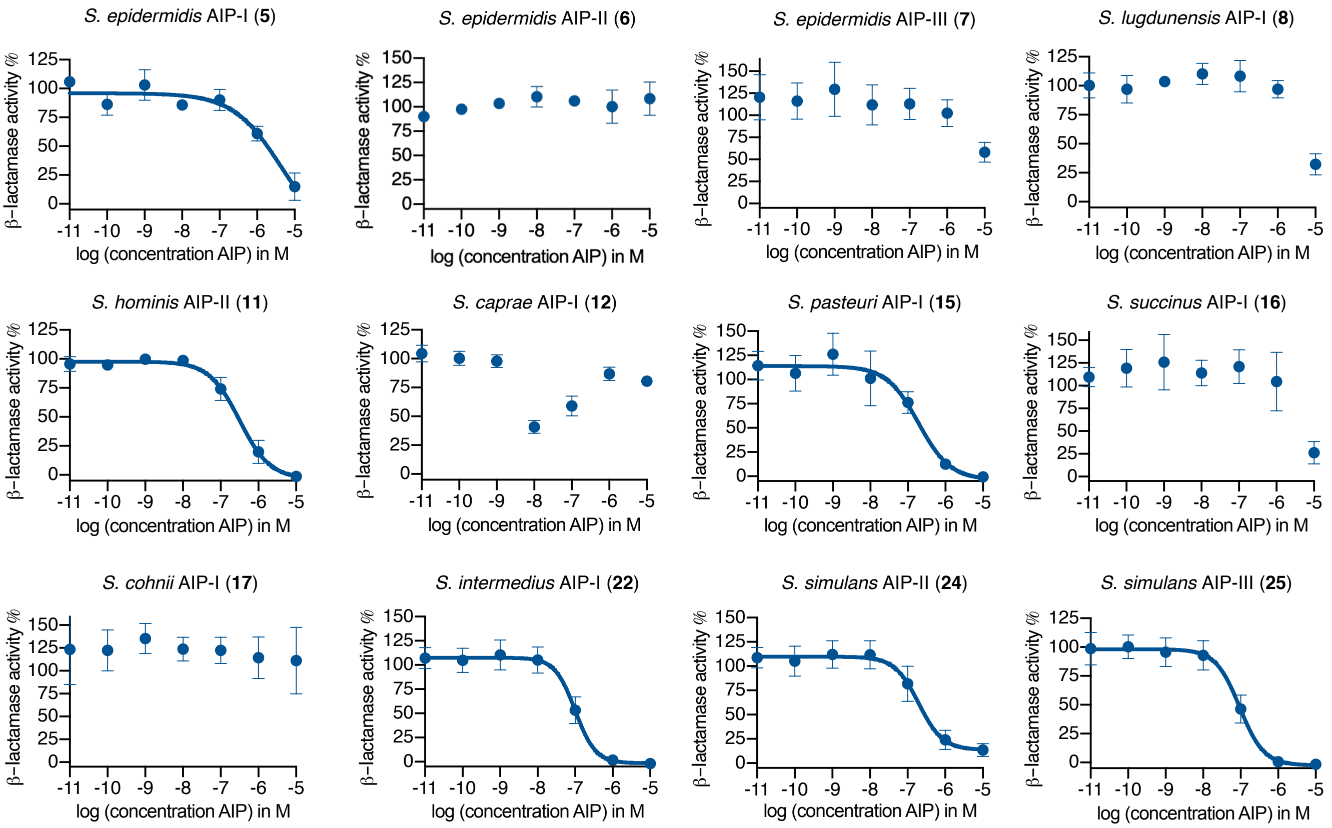

**Supplementary Figure 13. Dose-response curves of AgrC inhibition of *S. aureus agr*-III.** Inhibition properties of synthetic AIPs (10 µM to 10 pM) were determined through *β*-lactamase activity using the *S. aureus agr*-III reporter strain in the presence of 100 nM *S. aureus* AIP-III (**3**). The curves were generated from three individual assays performed in technical duplicate and shown error bars are the standard deviation of the mean (SD).

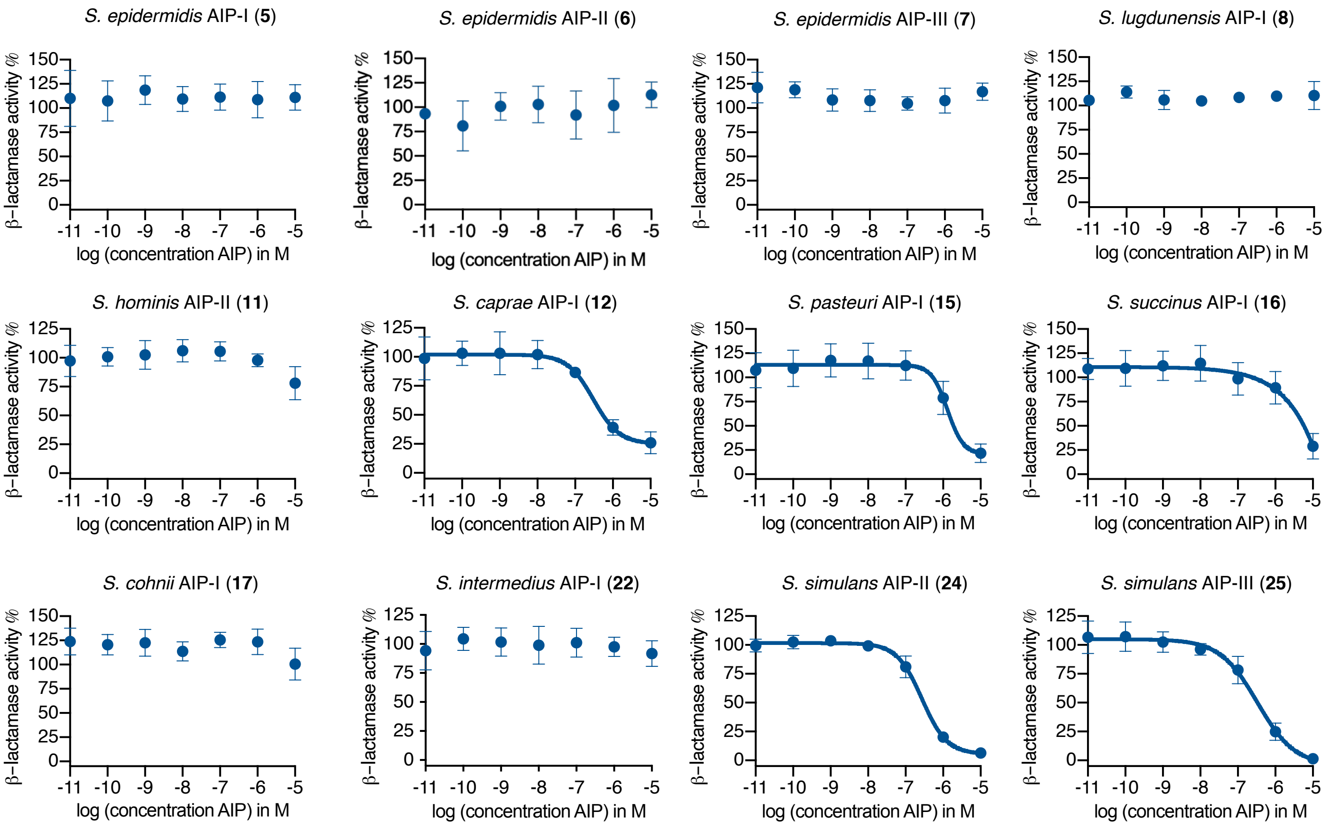

**Supplementary Figure 14. Dose-response curves of AgrC inhibition of *S. aureus agr*-IV.** Inhibition properties of synthetic AIPs (10 µM to 10 pM) were determined through *β*-lactamase activity using the *S. aureus agr*-IV reporter strain in the presence of 100 nM *S. aureus* AIP-IV (**4**). The curves were generated from three individual assays performed in technical duplicate and shown error bars are the standard deviation of the mean (SD).

#### Bar graphs for *agr* interference using fluorescence reporter assays

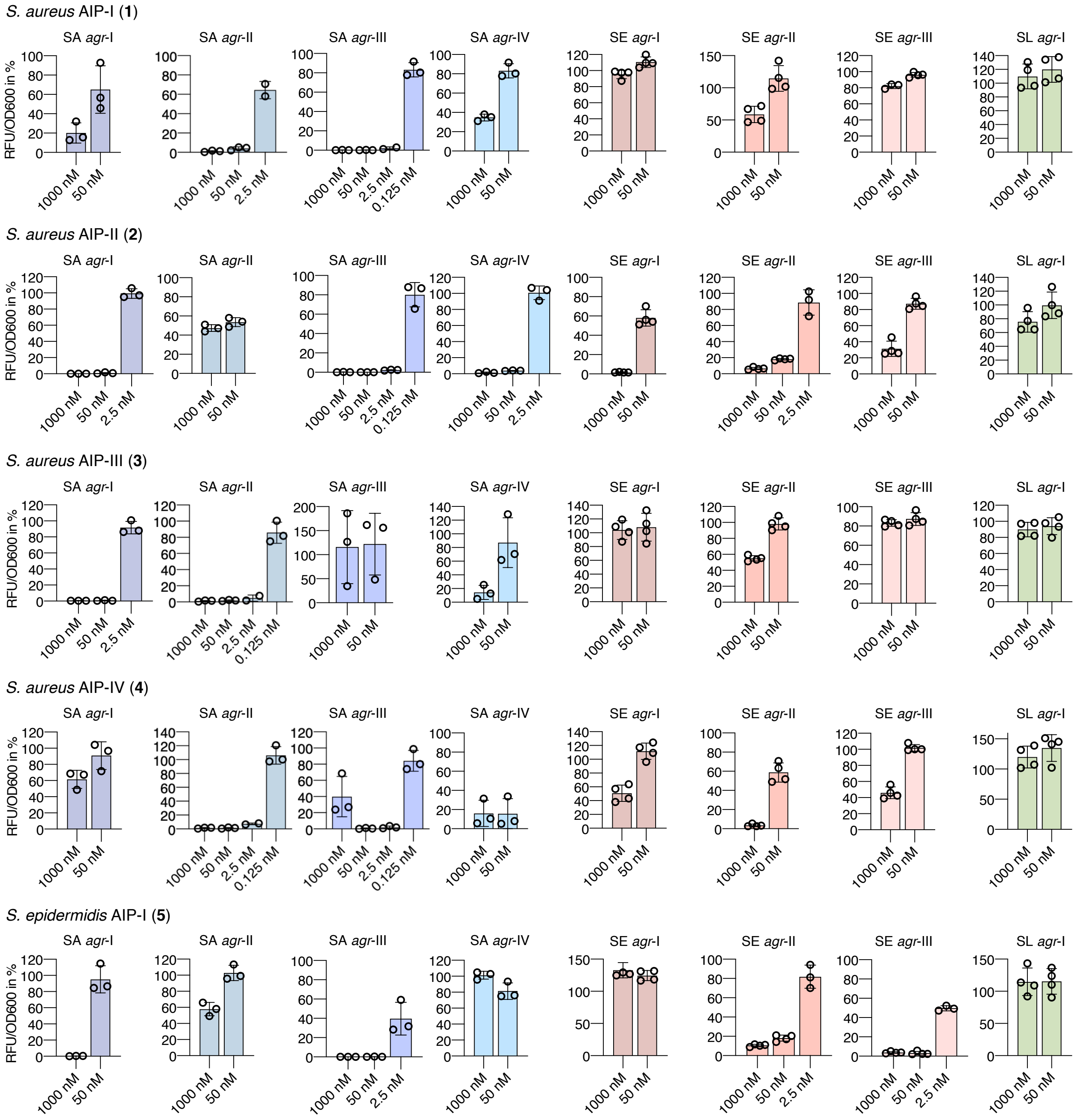

**Supplementary Figure 15. Fluorescence reporter strain assay for *agr* interference with synthetic AIPs.** Fluorescent reporter strains of *S. aureus* (SA), *S. epidermidis* (SE), S. *lugdunensis* (SL) were treated with AIPs at 1000 nM and 50 nM. AIPs were further tested at 2.5 nM and 0.125 nM in case >75% inhibition was observed at higher concentrations. Error bars are the standard error of the mean (SEM) of three individual biological assays performed in technical triplicate.

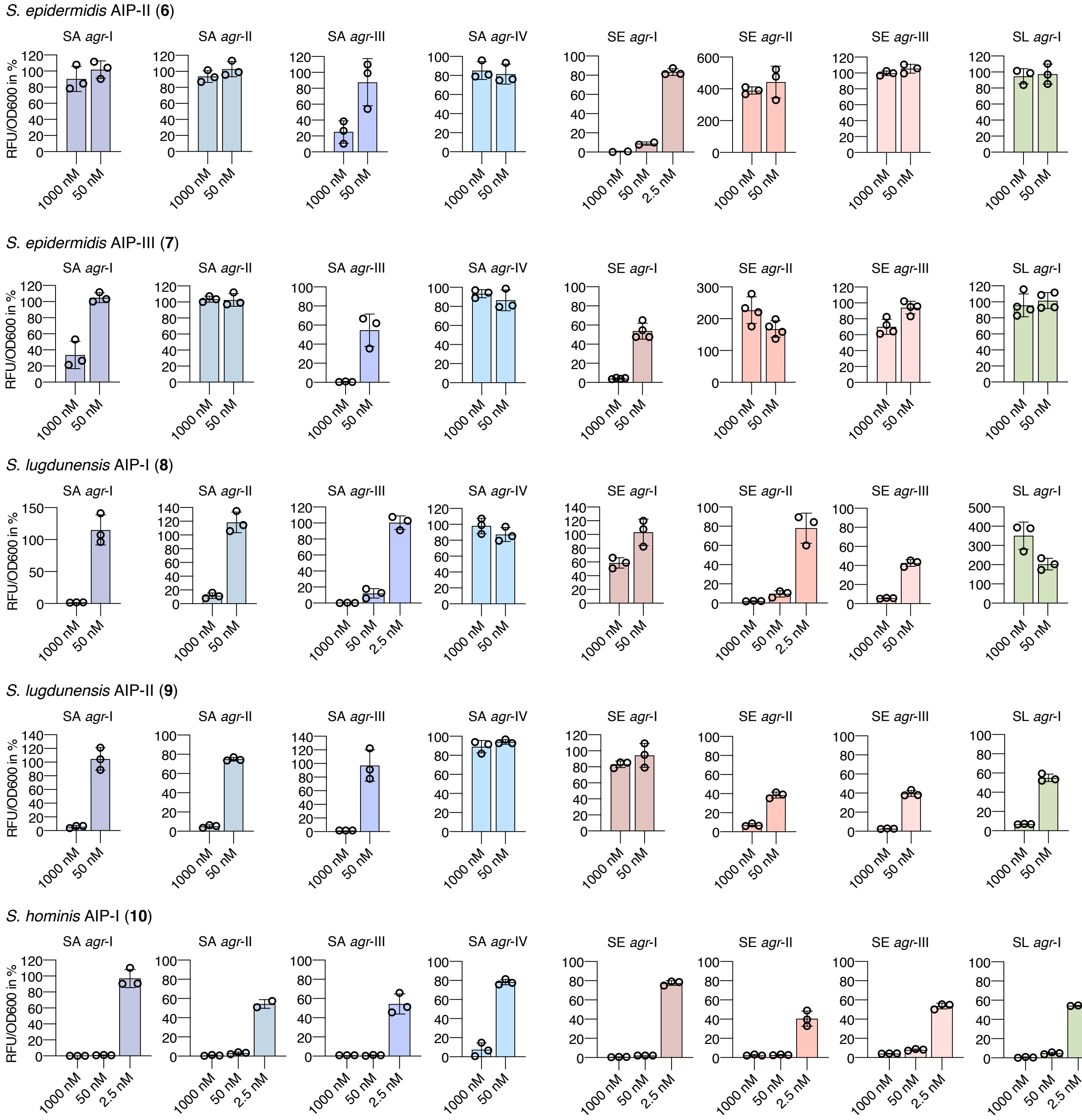

**Supplementary Figure 16. Fluorescence reporter strain assay for *agr* interference with synthetic AIPs.** Fluorescent reporter strains of *S. aureus* (SA), *S. epidermidis* (SE), S. *lugdunensis* (SL) were treated with AIPs at 1000 nM and 50 nM. AIPs were further tested at 2.5 nM and 0.125 nM in case >75% inhibition was observed at higher concentrations. Error bars are the standard error of the mean (SEM) of three individual biological assays performed in technical triplicate.

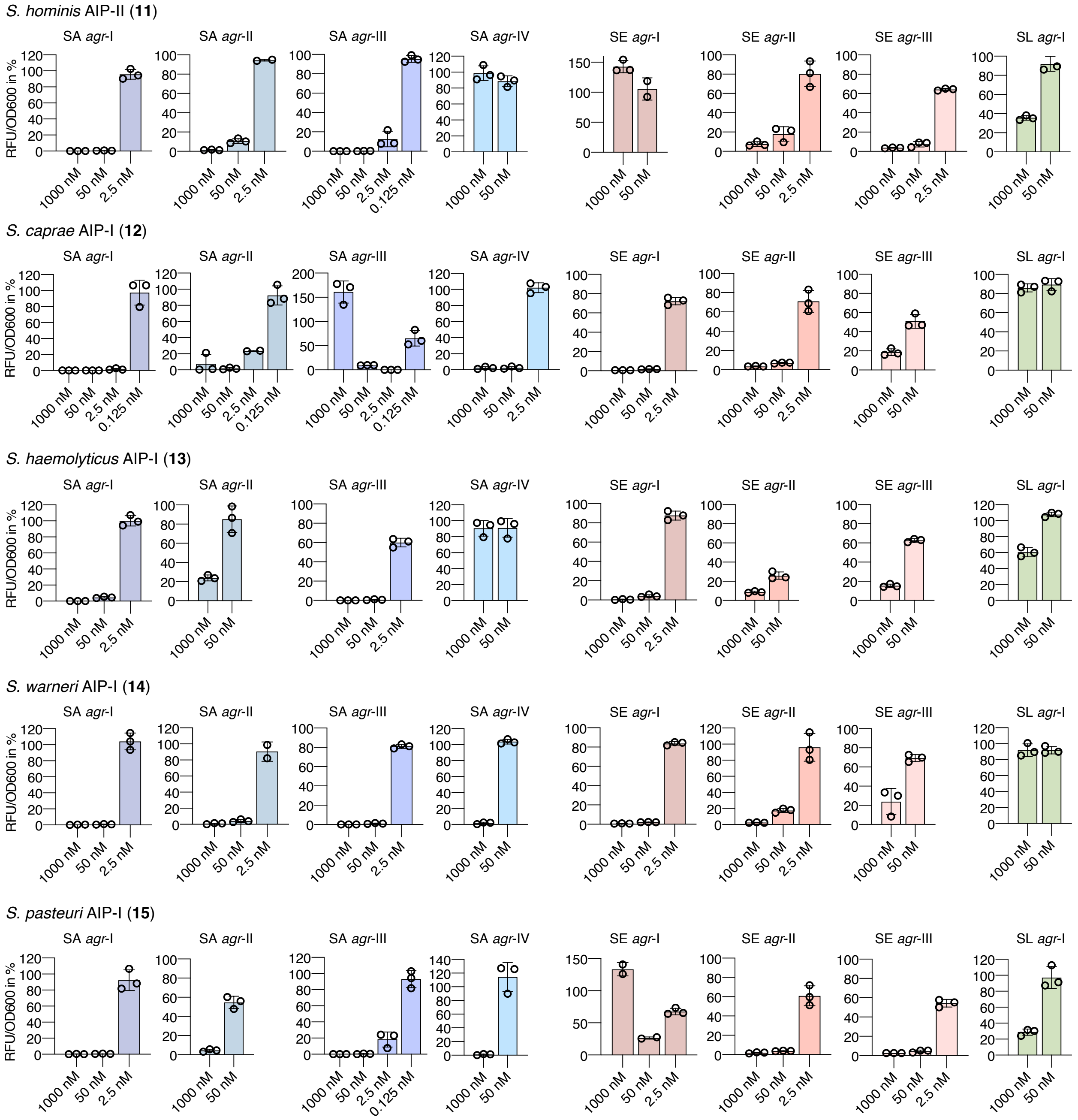

**Supplementary Figure 17. Fluorescence reporter strain assay for *agr* interference with synthetic AIPs.** Fluorescent reporter strains of *S. aureus* (SA), *S. epidermidis* (SE), S. *lugdunensis* (SL) were treated with AIPs at 1000 nM and 50 nM. AIPs were further tested at 2.5 nM and 0.125 nM in case >75% inhibition was observed at higher concentrations. Error bars are the standard error of the mean (SEM) of three individual biological assays performed in technical triplicate.

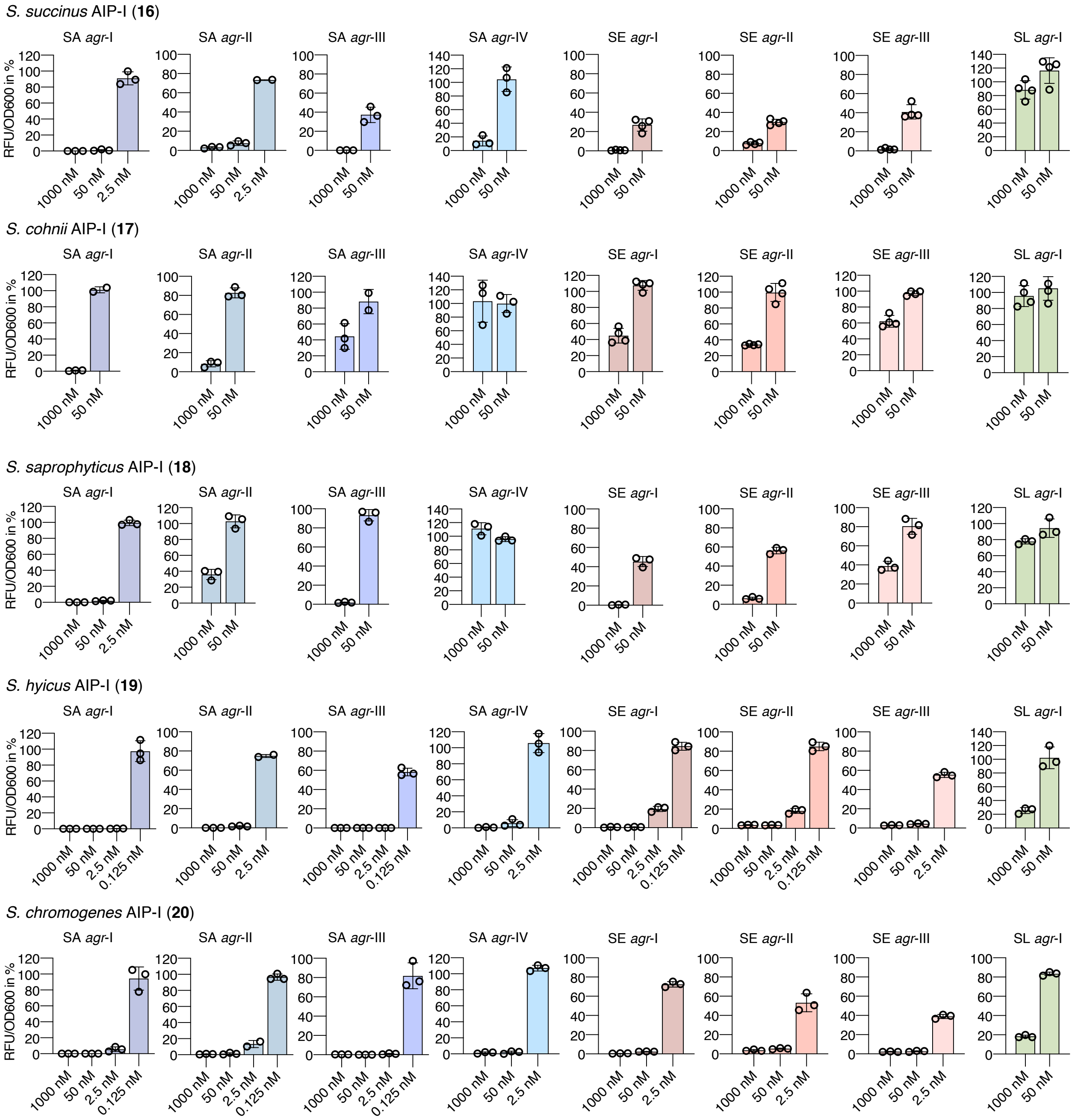

**Supplementary Figure 18. Fluorescence reporter strain assay for *agr* interference with synthetic AIPs.** Fluorescent reporter strains of *S. aureus* (SA), *S. epidermidis* (SE), S. *lugdunensis* (SL) were treated with AIPs at 1000 nM and 50 nM. AIPs were further tested at 2.5 nM and 0.125 nM in case >75% inhibition was observed at higher concentrations. Error bars are the standard error of the mean (SEM) of three individual biological assays performed in technical triplicate.

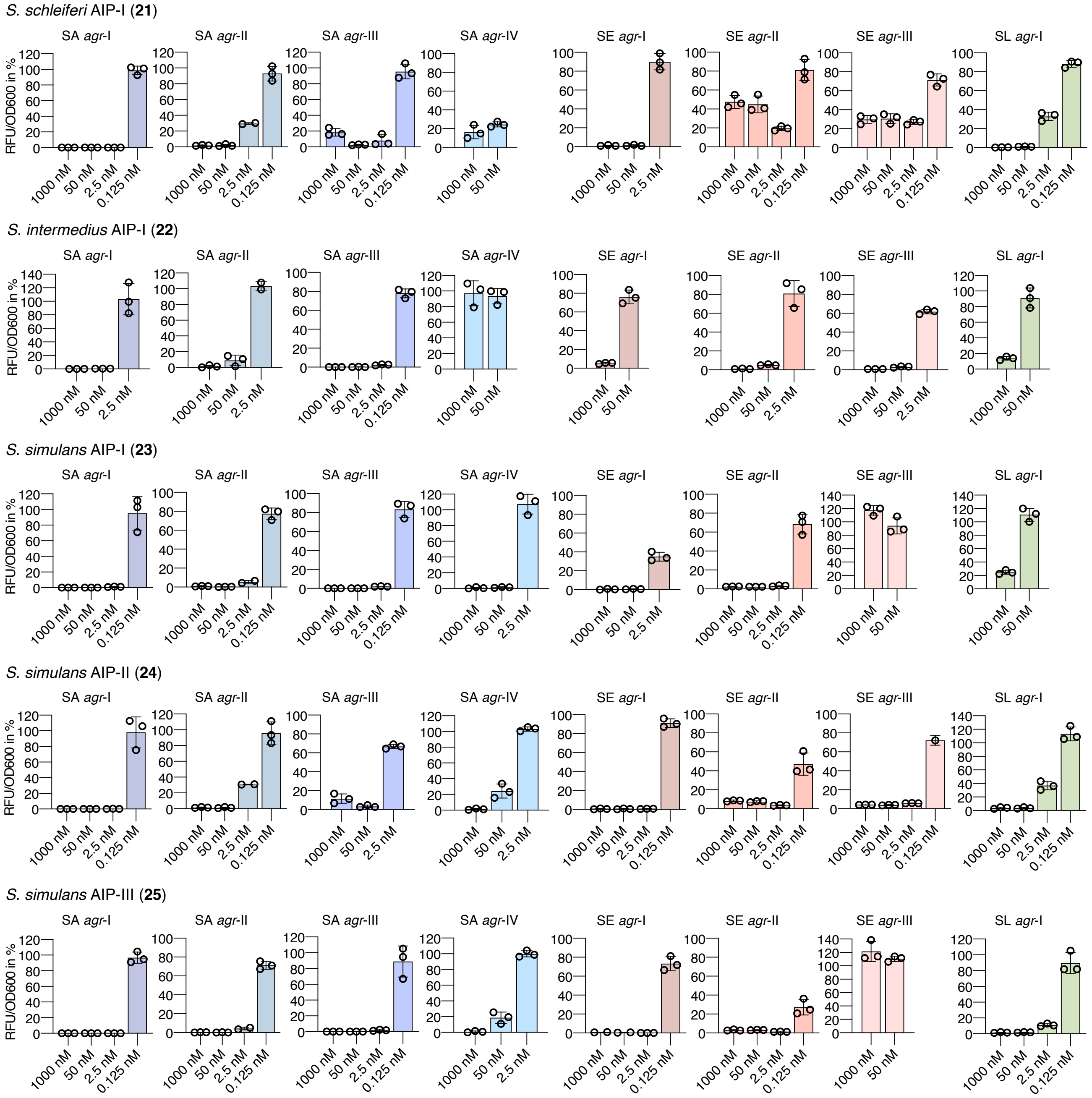

**Supplementary Figure 19. Fluorescence reporter strain assay for *agr* interference with synthetic AIPs.** Fluorescent reporter strains of *S. aureus* (SA), *S. epidermidis* (SE), S. *lugdunensis* (SL) were treated with AIPs at 1000 nM and 50 nM. AIPs were further tested at 2.5 nM and 0.125 nM in case >75% inhibition was observed at higher concentrations. Error bars are the standard error of the mean (SEM) of three individual biological assays performed in technical triplicate.

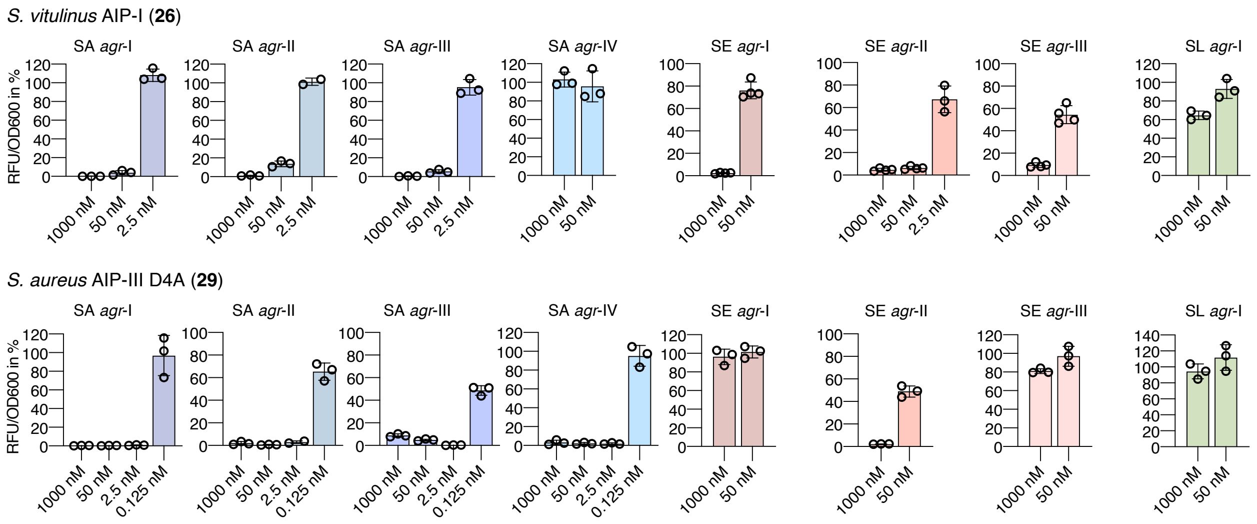

**Supplementary Figure 20. Fluorescence reporter strain assay for *agr* interference with synthetic AIPs.** Fluorescent reporter strains of *S. aureus* (SA), *S. epidermidis* (SE), S. *lugdunensis* (SL) were treated with AIPs at 1000 nM and 50 nM. AIPs were further tested at 2.5 nM and 0.125 nM in case >75% inhibition was observed at higher concentrations. Error bars are the standard error of the mean (SEM) of three individual biological assays performed in technical triplicate.

#### Overnight growth and fluorescence curves in presence of synthetic AIPs

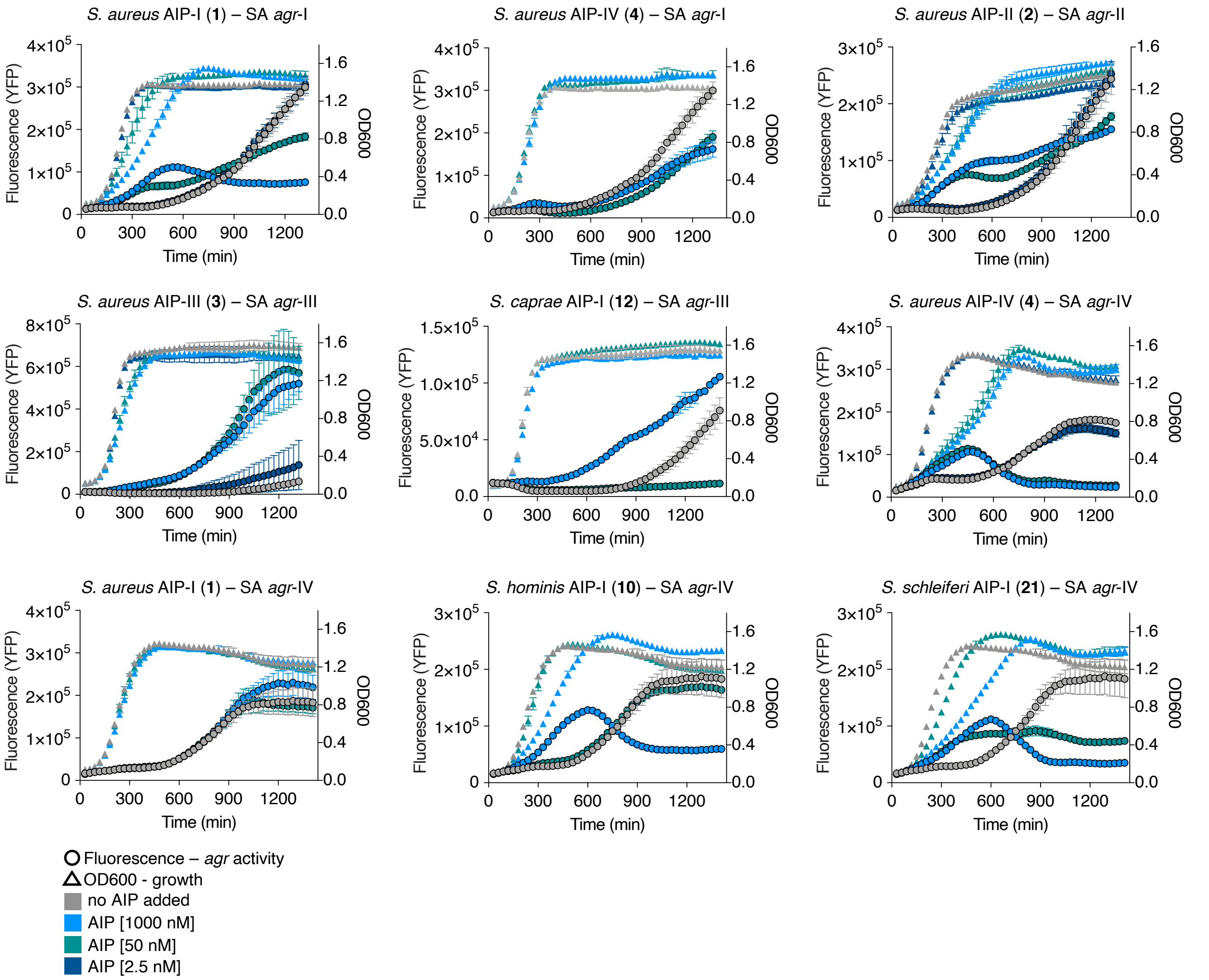

**Supplementary Figure 21. Fluorescence and $OD_{600}$ curves for selected AIPs with *S. aureus* (SA) *agr*-I–IV.** YFP-producing reporter strains of SA were grown overnight in the absence or presence of AIPs (1000 nM, 50 nM and 2.5 nM) with continuous measurements of fluorescence (YFP) and $OD_{600}$. The curves represent single experiments performed in technical triplicate. Data points are the mean and error bars are the standard deviation of the mean (SD).

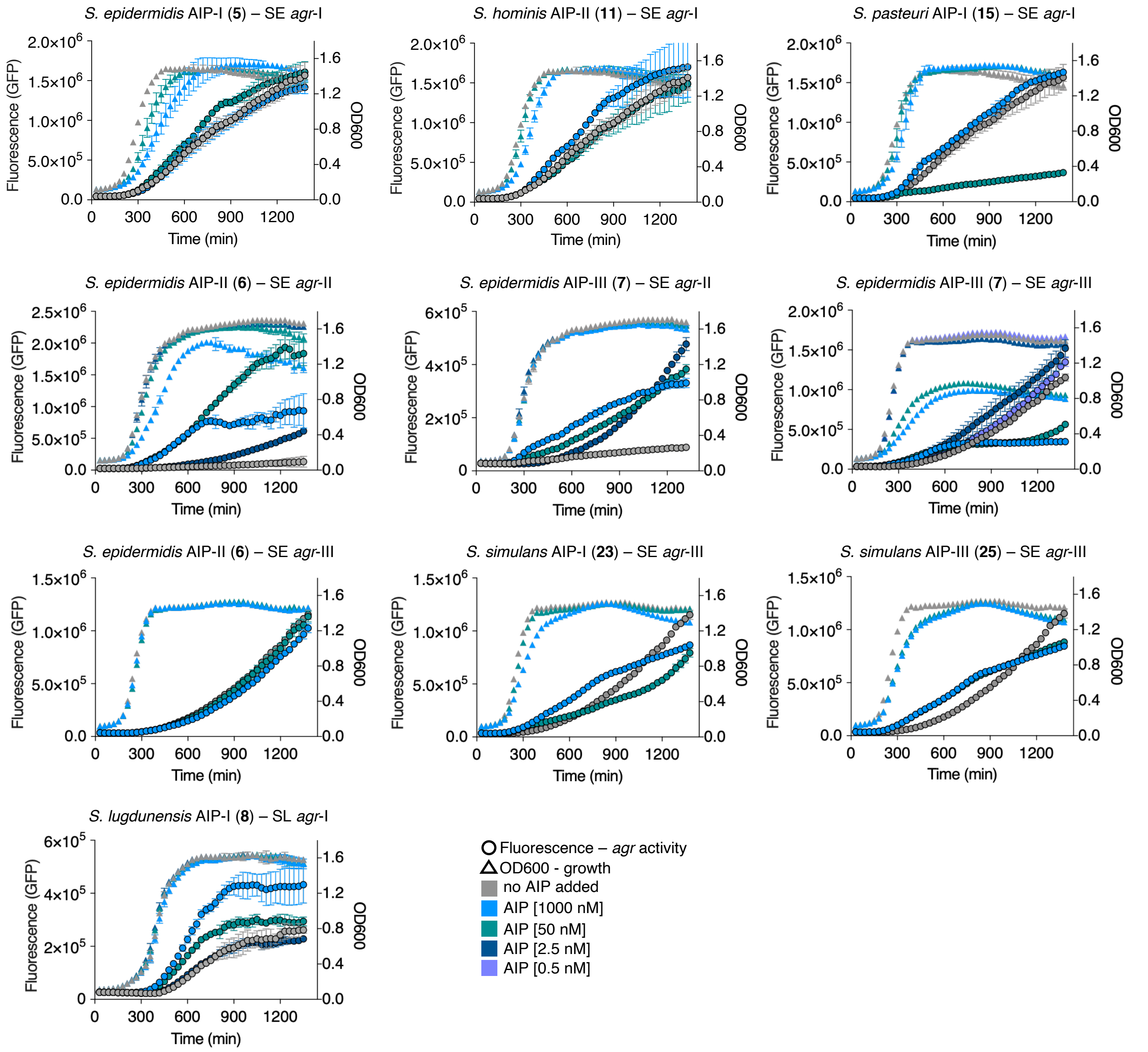

**Supplementary Figure 22. Fluorescence and $OD_{600}$ curves for selected AIPs with *S. epidermidis* (SE) *agr*-I–III and *S. lugdunensis* (SL) *agr*-I.** sGFP-producing reporter strains of SE and SL were grown overnight in the absence or presence of AIPs (1000 nM, 50 nM, 2.5 nM and 0.5 nM) with continuous measurements of fluorescence (GFP) and $OD_{600}$. The curves represent single experiments performed in technical triplicate. Data points are the mean and error bars are the standard deviation of the mean (SD).

#### Dose-response curves for *agr* interference using fluorescence reporter assays

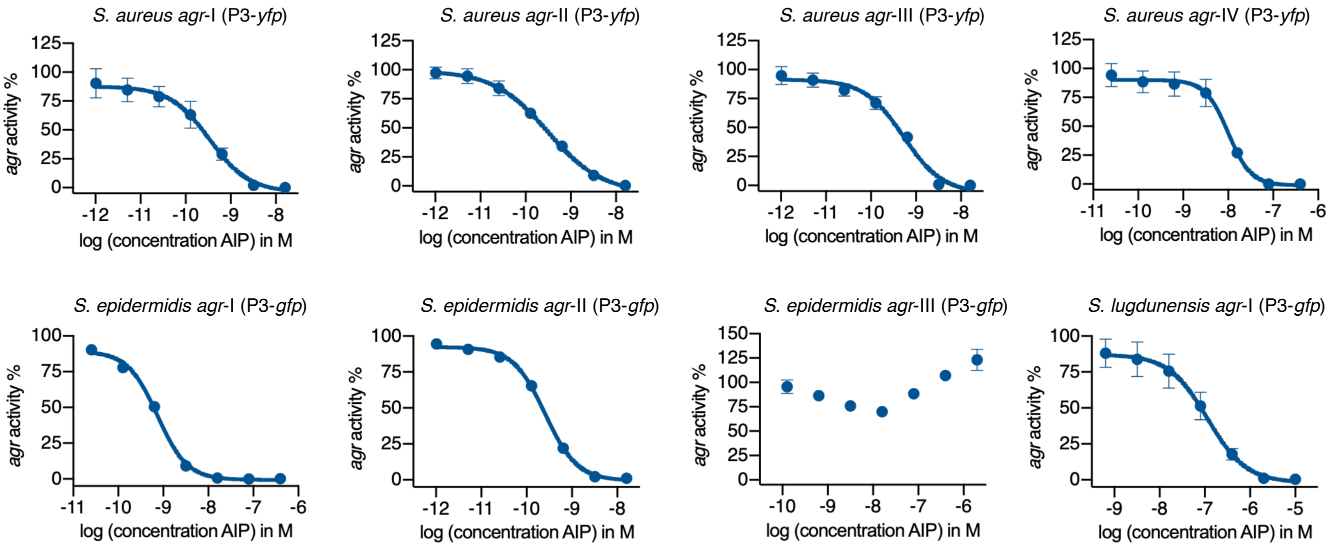

**Supplementary Figure 23. Dose-response ($IC_{50}$) curves for *agr* inhibition by *S. simulans* AIP-I (23).** Inhibition properties of *S. simulans* AIP-I (**23**) were determined through fluorescence readout as a measure of *agr* activity against reporter strains of SA, SE and SL. The curves were generated from three individual assays performed in technical duplicate and shown error bars are the standard deviation of the mean (SD).

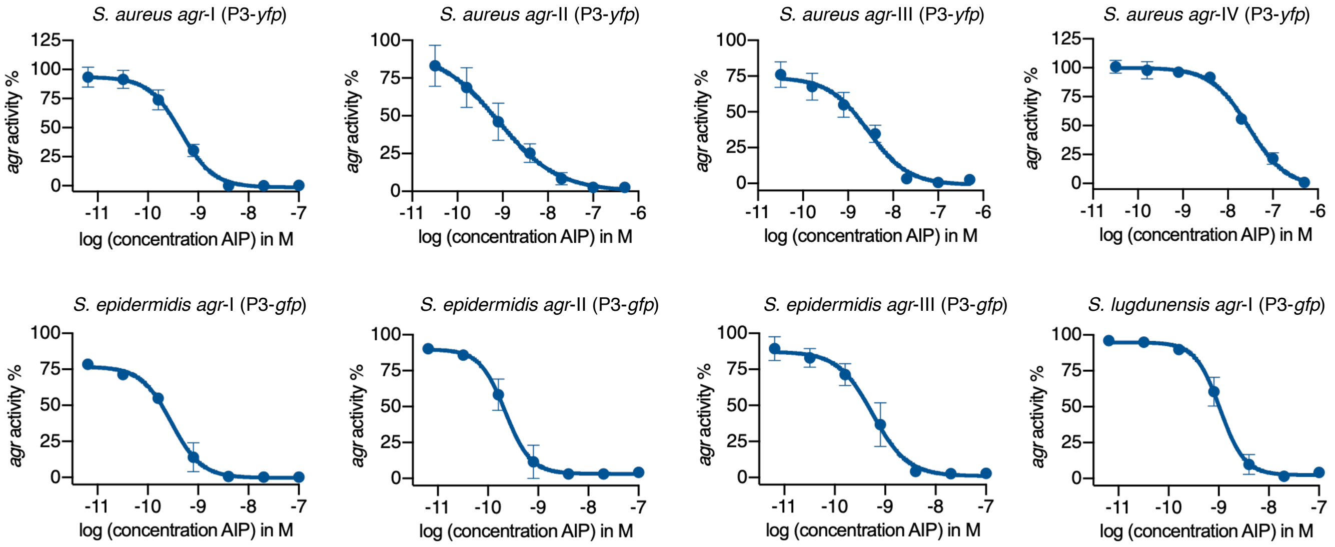

**Supplementary Figure 24. Dose-response ($IC_{50}$) curves for *agr* inhibition by *S. simulans* AIP-II (24).** Inhibition properties of *S. simulans* AIP-II (**24**) were determined through fluorescence readout as a measure of *agr* activity against reporter strains of SA, SE and SL. The curves were generated from three individual assays performed in technical duplicate and shown error bars are the standard deviation of the mean (SD).

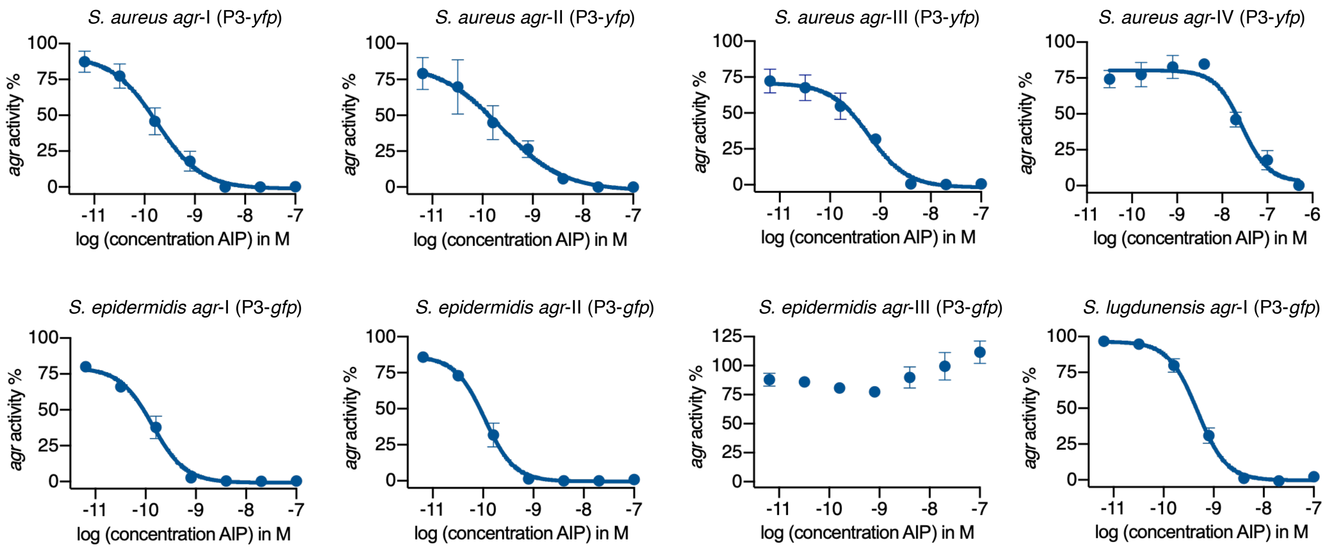

**Supplementary Figure 25. Dose-response ($IC_{50}$) curves for *agr* inhibition by *S. simulans* AIP-III (25).** Inhibition properties of *S. simulans* AIP-III (**25**) were determined through fluorescence readout as a measure of *agr* activity against reporter strains of SA, SE and SL. The curves were generated from three individual assays performed in technical duplicate and shown error bars are the standard deviation of the mean (SD).

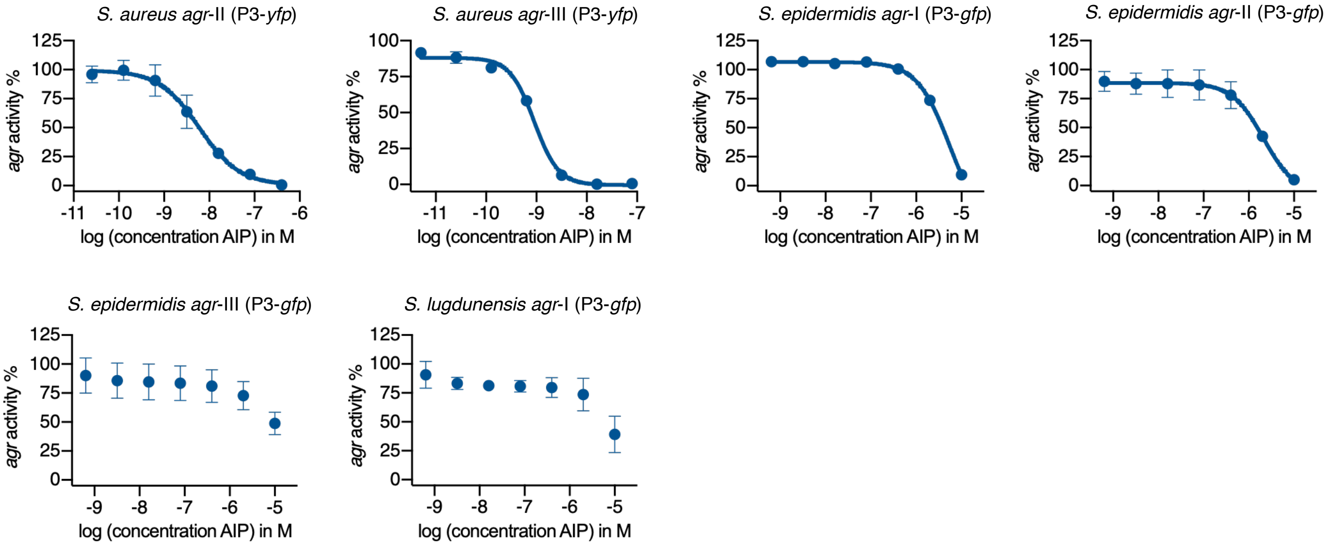

**Supplementary Figure 26. Dose-response ($IC_{50}$) curves for *agr* inhibition by *S. aureus* AIP-I (1).** Inhibition properties of *S. aureus* AIP-I (**1**) were determined through fluorescence readout as a measure of *agr* activity against reporter strains of SA, SE and SL. The curves were generated from three individual assays performed in technical duplicate and shown error bars are the standard deviation of the mean (SD).

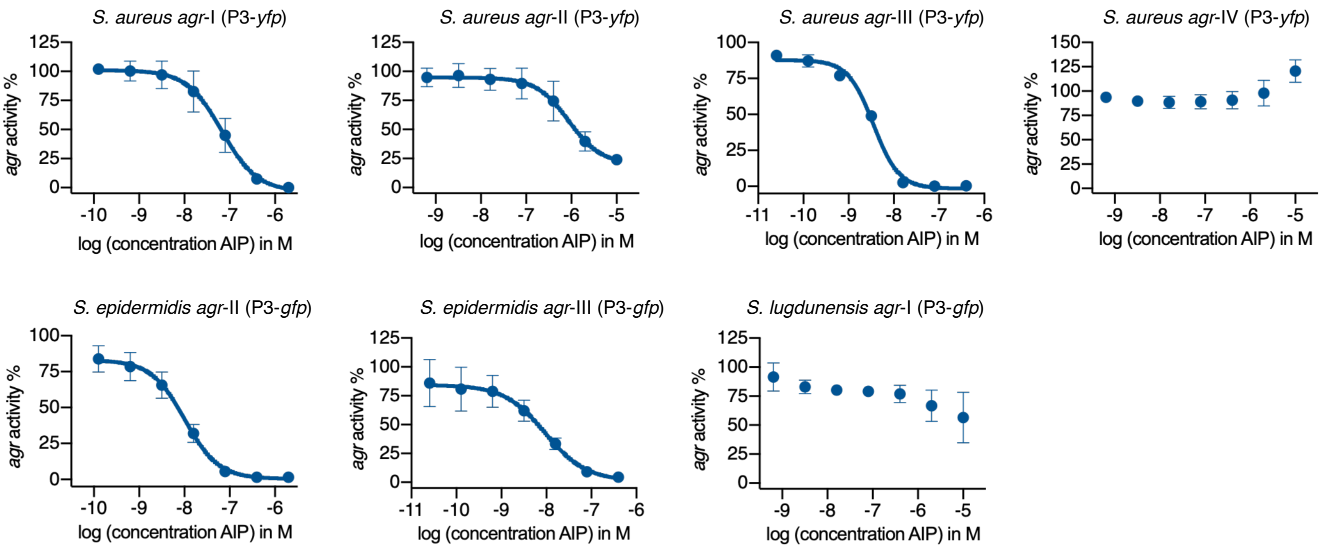

**Supplementary Figure 27. Dose-response ($IC_{50}$) curves for *agr* inhibition by *S. epidermidis* AIP-I (5).** Inhibition properties of *S. epidermidis* AIP-I (**5**) were determined through fluorescence readout as a measure of *agr* activity against reporter strains of SA, SE and SL. The curves were generated from three individual assays performed in technical duplicate and shown error bars are the standard deviation of the mean (SD).

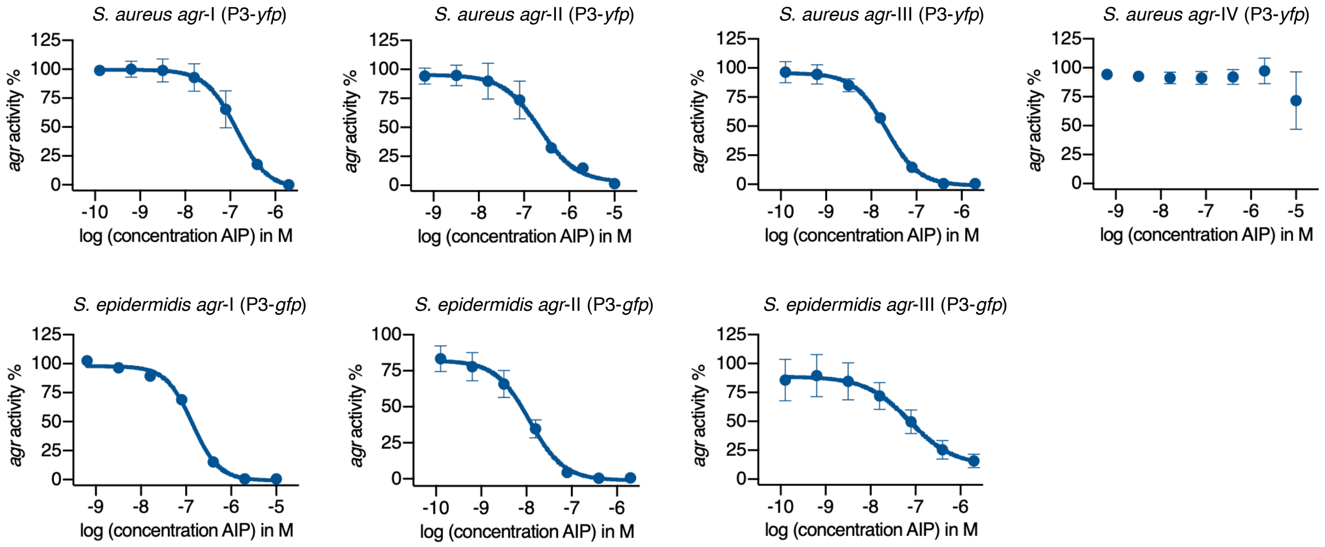

**Supplementary Figure 28. Dose-response ($IC_{50}$) curves for *agr* inhibition by *S. lugdunensis* AIP-I (8).** Inhibition properties of *S. lugdunensis* AIP-I (**8**) were determined through fluorescence readout as a measure of *agr* activity against reporter strains of SA and SE. The curves were generated from three individual assays performed in technical duplicate and shown error bars are the standard deviation of the mean (SD).

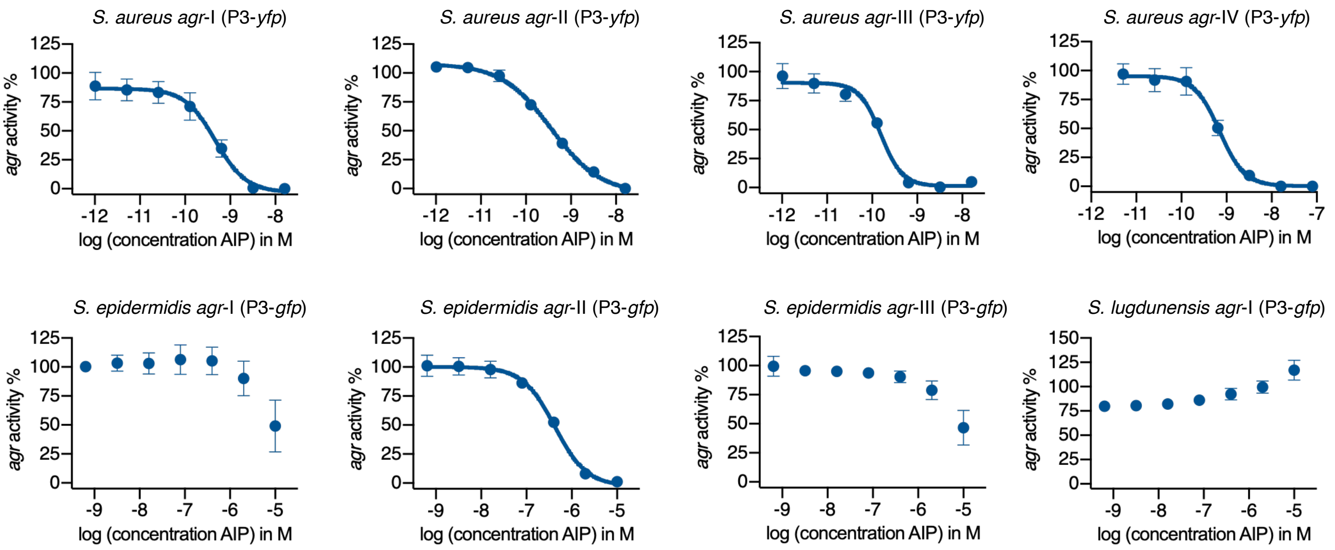

**Supplementary Figure 29. Dose-response ($IC_{50}$) curves for *agr* inhibition by *S. aureus* AIP-III D4A (27).** Inhibition properties of *S. aureus* AIP-III D4A (**27**) were determined through fluorescence readout as a measure of *agr* activity against reporter strains of SA, SE and SL. The curves were generated from three individual assays performed in technical duplicate and shown error bars are the standard deviation of the mean (SD).

#### 2. Supplementary tables

##### List of reported quorum sensing interactions of staphylococcal AIPs

**Supplementary Table 1. Reported QS interactions of staphylococcal AIPs against *S. aureus* (SA) *agr*-I–IV.**

| Species | SA *agr*-I | SA *agr*-II | SA *agr*-III | SA *agr*-IV | Assay type | Ref. |
|---|---|---|---|---|---|---|
| *S. aureus* AIP-I (**1**) | n. t. | **IC$_{50}$:** 3.4 ± 1.2 nM | **IC$_{50}$:** 3.1± 1.3 nM | n. t. | *β*-lactamase | 1 |
| | **EC$_{50}$:** 40 ± 9 nM | **IC$_{50}$:** 26 ± 7 nM | n. t. | n. t. | *β*-lactamase | 2 |
| | **EC$_{50}$:** 23 ± 7 nM | **IC$_{50}$:** 135 ± 52 nM | n. t. | n. t. | *β*-lactamase | 2 |
| | **EC$_{50}$:** 28 nM (18–46 CI) | **IC$_{50}$:** 25 nM (14–45 CI) | **IC$_{50}$:** 3 nM (2–5 CI) | **EC$_{50}$:** 26 μM (23–29 CI) | *β*-lactamase | 3 |
| | n. v. | **IC$_{50}$:** 8.00 nM (3.7–17.5 CI) | **IC$_{50}$:** 0.53 nM (0.31–0.88 CI) | n. v. | fluorescence | 4 |
| | n. t. | **IC$_{50}$:** 3.34 nM (0.67–16.5 CI) | **IC$_{50}$:** 6.12 nM (5.20–7.21 CI) | n. t. | hemolysis | 4 |
| | **EC$_{50}$:** 11 nM (9–12 CI) | n. t. | n. t. | n. t. | *β*-lactamase | 5 |
| | **EC$_{50}$:** 3.21 nM (1.21–8.57 CI) | n. t. | n. t. | n. t. | *β*-lactamase | 6 |
| *S. aureus* AIP-II (**2**) | **IC$_{50}$:** 2.9 ± 1.2 nM | n. t. | **IC$_{50}$:** 3.2 ± 1.3 nM | n. t. | *β*-lactamase | 1 |
| | **IC$_{50}$:** 90 ± 30 nM | **EC$_{50}$:** 34 ± 6 nM | n. t. | n. t. | *β*-lactamase | 2 |
| | **IC$_{50}$:** 78 ± 12 nM | **EC$_{50}$:** 28 ± 14 nM | n. t. | n. t. | *β*-lactamase | 2 |
| | **IC$_{50}$:** 40 nM (12–140 CI) | **EC$_{50}$:** 30 nM (10–90 CI) | **IC$_{50}$:** 1 nM (0.7–2.6 CI) | **IC$_{50}$:** 86 nM (65–111 CI) | *β*-lactamase | 3 |
| | **IC$_{50}$:** 1.62 nM (0.93–2.82 CI) | n. v. | **IC$_{50}$:** 0.53 nM (0.24–1.19 CI) | **IC$_{50}$:** 0.40 nM (0.21–0.76 CI) | fluorescence | 4 |
| | **IC$_{50}$:** 0.89 nM (0.37–2.15 CI) | n. t. | **IC$_{50}$:** 3.59 nM (1.22–10.6 CI) | **IC$_{50}$:** 1.19 nM (0.55–2.60 CI) | hemolysis | 4 |
| | n. t. | **EC$_{50}$:** 40.9 nM (30.3–55.3 CI) | n. t. | n. t. | *β*-lactamase | 6 |
| | **IC$_{50}$:** 12 ± 2.9 nM | n. t. | n. t. | n. t. | *β*-lactamase | 7 |
| *S. aureus* AIP-III (**3**) | **IC$_{50}$:** 70 nM (30–150 CI) | **IC$_{50}$:** 6 nM (5–6.5 CI) | **EC$_{50}$:** 26 nM (22–31 CI) | **IC$_{50}$:** 150 nM (104–207 CI) | *β*-lactamase | 3 |
| | **IC$_{50}$:** 5.05 nM (2.46–10.4 CI) | **IC$_{50}$:** 5.63 nM (1.89–16.7 CI) | n. v. | **IC$_{50}$:** 8.53 nM (4.15–17.5 CI) | fluorescence | 4 |
| | **IC$_{50}$:** 8.07 nM (4.34–15.0 CI) | **IC$_{50}$:** 0.46 nM (0.27–0.77 CI) | n. t. | **IC$_{50}$:** 23.8 nM (13.7–41.3 CI) | hemolysis | 4 |
| | n. t. | n. t. | **EC$_{50}$:** 406 nM (281–586 CI) | n. t. | *β*-lactamase | 6 |

| | | | | | | |
|---|---|---|---|---|---|---|
| | **$IC_{50}$:** 8 ± 1.1 nM | n. t. | n. t. | n. t. | *β*-lactamase | 7 |
| *S. aureus* AIP-IV (**4**) | **$EC_{50}$:** 62 nM (52–75 CI) | **$IC_{50}$:** 4 nM (3–5 CI) | **IC50:** 1 nM (0.5–3 CI) | **$EC_{50}$:** 13 nM (7–40 CI) | *β*-lactamase | 3 |
| | n. v. | **$IC_{50}$:** 0.37 nM (0.22–0.64 CI) | **IC50:** 0.46 nM (0.21–0.99 CI) | n. v. | fluorescence | 4 |
| | n. t. | **$IC_{50}$:** 0.090 nM (0.078–0.103 CI) | **IC50:** 1.49 nM (0.70–3.14 CI) | n. t. | hemolysis | 4 |
| | n. t. | n. t. | n. t. | **$EC_{50}$:** 7.90 nM (4.93–12.7 CI) | *β*-lactamase | 6 |
| *S. epidermidis* AIP-I (**5**) | **$IC_{50}$:** ~250 nM | **$IC_{50}$:** ~30-40 nM | **$IC_{50}$:** ~10 nM | no inhibition at 1000 nM | HPLC δ-toxin | 9 |
| | **$IC_{50}$:** 166 nM (68.7–402 CI) | **$IC_{50}$:** >1000 nM | **$IC_{50}$:** 13.0 nM (6.41–26.5 CI) | **$IC_{50}$:** >1000 nM | fluorescence | 8 |
| *S. epidermidis* AIP-II (**6**) | no inhibition at 100 nM | no inhibition at 100 nM | no inhibition at 100 nM | no inhibition at 100 nM | fluorescence | 10 |
| *S. lugdunensis* AIP-I (**8**) | **$IC_{50}$:** 384 nM (353–418 CI) | **$IC_{50}$:** 419 nM (353–418 CI) | **$IC_{50}$:** 36.6 nM (353–418 CI) | **$IC_{50}$:** >1000 nM | fluorescence | 8 |
| *S. hominis* AIP-II (**11**) | **$IC_{50}$:** 0.6243 nM | ~70% inhibition by supernatant | ~80% inhibition by supernatant | no inhibition by supernatant | fluorescence | 11 |
| *S. caprae* AIP-I (**12**) | **$IC_{50}$0:** 0.6 ± 0.02 nM | **$IC_{50}$:** 0.26 ± 0.08 nM | **$IC_{50}$:** 0.2 nM | **$IC_{50}$:** 8.98 ± 0.32 nM | fluorescence | 12 |
| *S. intermedius* AIP-I (**22**) | 80–100% inhibition from supernatant | 40–90% inhibition from supernatant | 60–100% inhibition from supernatant | 10–60% inhibition from supernatant | *β*-lactamase | 13 |
| *S. simulans* AIP-I (**23**) | **$IC_{50}$:** 2.2 nM | **$IC_{50}$:** 1.1 nM | **$IC_{50}$:** 3.5 nM | **$IC_{50}$:** 23 nM | fluorescence | 14 |
| *S. simulans* AIP-II (**24**) | **$IC_{50}$:** 1.6 nM | **$IC_{50}$:** 15 nM | **$IC_{50}$:** 11.5 nM | **$IC_{50}$:** 40 nM | fluorescence | 14 |
| *S. simulans* AIP-III (**25**) | **$IC_{50}$:** 1.7 nM | **$IC_{50}$:** 6.0 nM | **$IC_{50}$:** 3.2 nM | **$IC_{50}$:** 48 nM | fluorescence | 14 |
| *S. aureus* AIP-III D4A (**27**) | **$IC_{50}$:** 0.49 nM (0.29–0.81 CI) | **$IC_{50}$:** 0.43 nM (0.21–0.89 CI) | **$IC_{50}$:** 0.051 nM (0.023–0.113CI) | **$IC_{50}$:** 0.035 nM (0.012–0.099 CI) | fluorescence | 4 |
| | **$IC_{50}$:** 0.082 nM (0.047–0.142 CI) | **$IC_{50}$:** 0.060 nM (0.038–0.094 CI) | **$IC_{50}$:** 0.16 nM (0.069–0.39 CI) | **$IC_{50}$:** 0.11 nM (0.066–0.17 CI) | hemolysis | 4 |
| | **$IC_{50}$:** 0.16 ± 0.01 nM | n. t. | n. t. | n. t. | *β*-lactamase | 7 |

**Supplementary Table 2. Reported QS interactions of staphylococcal AIPs against *S. epidermidis* (SE) *agr*-I–III.**

| Species | SE *agr*-I | SE *agr*-II | SE *agr*-III | Assay type | Ref. |
|---|---|---|---|---|---|
| *S. aureus* AIP-I (**1**) | no inhibition at 1000 nM | n. t. | n. t. | HPLC δ-toxin | 9 |
| | no inhibition at 1000 nM | n. t. | n. t. | fluorescence | 10 |
| *S. aureus* AIP-II (**2**) | no inhibition at 1000 nM | n. t. | n. t. | HPLC δ-toxin | 9 |
| | **$IC_{50}$:** 62.9 nM (26.1–151 CI) | n. t. | n. t. | fluorescence | 10 |
| *S. aureus* AIP-III (**3**) | no inhibition at 1000 nM | n. t. | n. t. | HPLC δ-toxin | 9 |
| | no inhibition at 1000 nM | n. t. | n. t. | fluorescence | 10 |
| *S. aureus* AIP-IV (**4**) | 40% inhibition at 1000 nM | n. t. | n. t. | HPLC δ-toxin | 9 |
| | **$IC_{50}$:** >1000 nM | n. t. | n. t. | fluorescence | 10 |
| *S. epidermidis* AIP-I (**5**) | activation from supernatant | ~90% inhibition from supernatant | ~90% inhibition from supernatant | fluorescence | 15 |
| | **$EC_{50}$:** 196 nM (162–238 CI) | n. t. | n. t. | fluorescence | 10 |
| *S. epidermidis* AIP-II (**6**) | ~90% inhibition from supernatant | activation from supernatant | no effect from supernatant | fluorescence | 15 |
| | **$IC_{50}$:** 9.64 nM (7.99–11.6 CI) | activation at 10 µM | no effect at 10 µM | fluorescence | 10 |
| *S. epidermidis* AIP-III (**7**) | ~50% inhibition from supernatant | no effect from supernatant | activation from supernatant | fluorescence | 15 |
| | **$IC_{50}$:** 34.3 nM (31.4–37.4 CI) | n. t. | n. t. | fluorescence | 10 |
| *S. aureus* AIP-III D4A (**27**) | ~25% inhibition at 10 µM | ~70% inhibition at 100 nM | ~25% inhibition at 10 µM | fluorescence | 10 |

#### AgrD and AgrC sequences with accession numbers

##### Supplementary Table 3. List of AgrD sequences retrieved from NCBI.

| Species | *agr* | Accession | AgrD sequence |
|---|---|---|---|
| *S. agnetis* | *agr*-I | WP_107368992.1 | MAFFDSLLNLLTVLFKSLGNFARINPCTGFFNEPEVPHELSEAE |
| | *agr*-II | WP_081858219.1 | MAFFESVLKLFTIFFKTLGNFAAWDPCTGYFDEPEIPQELREAK |
| *S. argensis* | *agr*-I | WP_103371625.1 | MAILDTLFKILINVFTYFGNIAAKVGCGAYIDEPTVPKELIDKMDK |
| *S. argenteus* | *agr*-I | WP_001093930.1 | MNTLFNLLFELITGILKNIGNIAAYSTCDFIMDEVEVPKELTQLHE |
| *S. arlettae* | *agr*-I | WP_107376912.1 | MNLLNAFFSFFAKFFELIGTVAGVNPCGGWFDEPEVPEELTKLSE |
| | *agr*-II | WP_002509117.1 | MNLLNSFFSFFAKFFELIGTVAGINPCSGWFDEPEVPEELTKLYE |
| *S. aureus* | *agr*-I | WP_001093929.1 | MNTLFNLFFDFITGILKNIGNIAAYSTCDFIMDEVEVPKELTQLHE |
| | *agr*-II | WP_001094921.1 | MNTLVNMFFDFIIKLAKAIGIVGGVNACSSLFDEPKVPAELTNLYDK |
| | *agr*-III | WP_000735197.1 | MKKLLNKVIELLVDFFNSIGYRAAYINCDFLLDEAEVPKELTQLHE |
| | *agr*-IV | WP_001094303.1 | MNTLLNIFFDFITGVLKNIGNVASYSTCYFIMDEVEIPKELTQLHE |
| *S. auricularis* | *agr*-I | WP_083498188.1 | MKLVNLLLSSTTSFLQMVGNRQKAKTCTVLYDEPEVPKELTQELEK |
| | *agr*-II | WP_107392671.1 | MKLLDLLLSSTTSFLQMVGNRSKTKTCTVLYDEPEVPKELIQELEK |
| *S. caeli* | *agr*-I | WP_069996593.1 | MNILESIFKLIAKFFSVIGAMSGVRPCTAFADEPEIPKELTKLYE |
| *S. capitis* | *agr*-I | WP_030063609.1 | MEGLFNLFIKFFTMIFEFIGFVAGANPCASYFDEPEVPKELSKLYE |
| | *agr*-II | WP_107361624.1 | MNSLFNLIFKFFTMIFEFIGFVAGANPCQLYYDEPEVPEELTKLYE |
| | *agr*-II | WP_049428106.1 | MDALFNLVLKFFTIIFEFIGFVAGANPCALYYDEPEVPDELSKLYE |
| | *agr*-IV | WP_064210779.1 | MMQIFDVLFKVISFIFEKIGFIAGYSTCNTYFDEPEVPKELFETYQK |
| *S. caprae* | *agr*-I | WP_002445331.1 | MMQIINLLFKVITAVFEKIGFIAGYSTCSYYFDEPEVPKELLEIYKK |
| | *agr*-II | AAL65821.1 | MKMMQIFDLLFKVISFIFEKIGFLAGYRTCNTYFDEPEVPKELFETYQK |
| *S. carnosus* | *agr*-I | WP_015900791.1 | MDILNGIFKFFAFIFEQIGNIAKYNPCVGYFDEPEVPSELLDEQK |
| *S. casei* | *agr*-I | PNZ56808.1 | MNIFESILNLFAKFFSVLGVMAGAKPCGGFFDEPEVPSEITKLYE |
| *S. chromogenes* | *agr*-I | WP_107381457.1 | MAIFETLFNLFTTLFKTLGNLASINPCTGFFDEPEVPQELRETE |
| | *agr*-II | WP_069191685.1 | MAIFETLFNLFTALFKTLGNFAAMDPCTGFFDEPEVPQELRETE |
| | *agr*-III | WP_103159704.1 | MAIFETLFNLFINVFKALGNFASINPCTAFFDEPEVPKELREVE |
| *S. coagulans* | *agr*-I | WP_050345386.1 | MKFLDSLLKLLTSVFASISTYAKYPFCIGYFYEPEIPAELLEEE |
| *S. cohnii* | *agr*-I | WP_064264468.1 | MNFFESIITVFAKFFAFIGTISSVKPCTGFVDEPEIPKELTDLYK |
| | *agr*-II | WP_019469556.1 | MHIFESIINLFVKFFSVLGAVSGGKVCSAYFDEPEVPKEITDLYK |
| | *agr*-III | WP_073345993.1 | MNIFESILTVFAKFFAFIGTISSVKPCTGFADEPEIPKELTDLYK |
| | *agr*-IV | WP_107505417.1 | MHIFESIVNLFVKFFSVLGAVSGGKVCSAFFDEPEVPKEITDLYK |
| *S. condimenti* | *agr*-I | WP_047132797.1 | MDILNGIFKFFAFIFEQLGNIAKYNPCAGYFDEPEVPKELLEENK |
| *S. croceilyticus* | *agr*-I | WP_103328222.1 | MNLLSVLFAKVFSFIFGLVGTFSVQDVCSLYFDEPEVPEELQNLSDK |
| *S. delphini* | *agr*-I | WP_096597589.1 | MRILEVLFNLITNLFQSIGSFARIPISTGFFDEPEIPSELLEEDK |
| *S. devriesei* | *agr*-I | WP_103165878.1 | MMFITDLFFKFFAAILETLGNVAAYKPCFGYFDEAEVPEELTNLKR |
| | *agr*-II | WP_107508274.1 | MMFFADLFFKFFTAILETLGNVAAYKVCVVYFDEPEVPEELTNLQR |
| *S. edaphicus* | *agr*-I | WP_099091241.1 | MNIFETIFTFIAKFFAVIGAMSGVRPCTAYADEPEIPEELTKLYE |
| *S. epidermidis* | *agr*-I | WP_001830021.1 | MENIFNLFIKFFTTILEFIGTVAGDSVCASYFDEPEVPEELTKLYE |
| | *agr*-II | WP_002447513.1 | MNLLGGLLLKIFSNFMAVIGNASKYNPCSNYLDEPQVPEELTKLDE |
| | *agr*-III | WP_002490348.1 | MNLLGGLFLKIFSNFMAVIGNAAKYNPCASYLDEPQVPEELTKLDE |
| | *agr*-IV | WP_002487333.1 | MNLLGGLLLKIFSNFMAVIGNASKYNPCVMYLDEPQVPEELTKLDE |
| *S. equorum* | *agr*-I | WP_021339970.1 | MHIFESIFSIIAKFFTVLGAVAGARPCYGYFDETEVPKEITELYE |
| | *agr*-II | WP_002507141.1 | MHIFESIFSLIAKFFSTLGAVAGLRPCGGYFDEPEVPKEITELYE |
| | *agr*-III | WP_046466292.1 | MHIFESIFSLIAKFFSTLGAVAGLSPCGGYFDEPEVPKEITDLYE |
| | *agr*-IV | WP_069817770.1 | MHIFESIFSLIAKFFSTLGAVAGLRPCYGYFDETEVPKEITELYE |
| | *agr*-V | WP_056935777.1 | MFIYEVFVNLITKIFSALGTIAAINPCMGYFDEPEVPKEISKLYE |
| | *agr*-VI | WP_065338397.1 | MHIFESIFSIIAKFFSTLGAAAGLRPCGGYFDEPEVPKEITDLYE |
| *S. felis* | *agr*-I | WP_115856975.1 | MNFLENVLSLITKLFQVIGNFAKINTCTGFFNEPEVPSELRDSSK |
| | *agr*-II | WP_103207460.1 | MNFLENVLSLITKLFQVIGNFAKINTCTVYFDEPEVPRELLDSNK |
| | *agr*-III | WP_115871108.1 | MNIFESILSLISKLFQAIGNFAAIETCHYFFDEPEVPRELLDFEK |
| | *agr*-IV | WP_115902416.1 | MNIFESILSLISKLFQAIGNFAAIETCHGFFDEPEVPRELIDFEK |
| | *agr*-V | WP_115924802.1 | MNFLENVLSLITKLFQVIGNFAKINTCTGYFDEPEIPRELLDSNK |

| | | | |
|---|---|---|---|
| *S. fleurettii* | *agr*-I | WP_078358467.1 | MNLLTSLFSKSVSNILSAIGEKAVIRGCMAFLDETEVPAELLKEKQ |
| *S. gallinarum* | *agr*-I | WP_042739402.1 | MNILDSLLNLATKFFSALGASVGARPCGGFFDEPEVPAEITELHK |
| | *agr*-II | WP_107512825.1 | MNILDSLLNLATKLFSTLGATVGALPCGGFFDEPEVPAEITELHK |
| | *agr*-III | WP_107590068.1 | MNILDSVFNLAIKLFSTLGAAVKASPCGGYFDEPEIPAELLELHK |
| | *agr*-IV | WP_107527999.1 | MSILESLVSLATKLFSALGTAAKASPCGGFFDEPEVPAELTELHK |
| | *agr*-V | WP_107580325.1 | MNILDSLLNLATKFFSALGTAVGAKPCGGFFDEPEVPAEITELHK |
| | *agr*-VI | WP_119486062.1 | MNILDSLVSLTTKLFSALGAAAQASPCGGFFDEPEIPAELTELHK |
| *S. haemolyticus* | *agr*-I | WP_011275303.1 | MTVLVDLIIKLFTFLLQSIGTIASFTPCTTYFDEPEVPEELTNAK |
| | *agr*-II | WP_033079904.1 | MTVLFDLFIKFFSFLLESVGTLASYTPCATYFDEPEVPEELTNLKR |
| | *agr*-III | WP_070855048.1 | MNLLTMLFAKVFSFIFGLVGTFSVQDVCSFYFDEPEVPKELQDLSKK |
| | *agr*-IV | RIO86649.1 | MTVLVDLIIKLFTFLLQSIGTVASFTPCTTYFDEPEVPEELTNTKN |
| *S. hominis* | *agr*-I | WP_017175278.1 | MTFITDLFIKLFSLILETVGTLATYSTCYGYFDESEVPEELTNLER |
| | *agr*-II | WP_002488089.1 | MTFITDLFIKLFSLILETVGTLASYNVCGGYFDEPEVPKELTDLNR |
| | *agr*-III | WP_019835517.1 | MTFITQLFIKLFSLILETVGTLASYSPCATYFDEPEVPEELTNLER |
| | *agr*-IV | WP_049413902.1 | MTFITQLFIKLFSFFLETIGNIATINTCGGYFDEPEVPQELTDLKK |
| | *agr*-V | WP_002448265.1 | MTFITDLFIKLFSLILETVGTLASQTVCSGYFDEPEVPKELTNLKR |
| | *agr*-VI | WP_070663365.1 | MTFFTDLFIKLFSLVLETVGTLASKTVCSGYFDEPEVPEELTNLKR |
| *S. hyicus* | *agr*-I | WP_039644712.1 | MAFFESLLNLLTSVFKSLGNFAKINPCTVFFDEPEVPKELRESE |
| | *agr*-II | WP_107633492.1 | MAFFESLLNLLTILFKSLGNFAKINPCTAFFDEPDVPEELKANK |
| | *agr*-III | WP_167694634.1 | MAFFEAIFKLFTIFFKTLGNFAAMDPCTAYFDEPEVPQELREAK |
| *S. intermedius* | *agr*-I | WP_019167558.1 | MRILEVLFNLITNLFQSIGTFARIPTSTGFFDEPEIPAELLEEEK |
| *S. kloosii* | *agr*-I | WP_061855583.1 | MDILNSIFSFFAKFFEVLGAVSGVNPCSGWFDEPEVPEELTKLYE |
| *S. lentus* | *agr*-I | WP_016999092.1 | MNLLAGLFSKTISNVLSAIGEKAVIRGCTAFLDETEVPAELLKEKQ |
| *S. lugdunensis* | *agr*-I | WP_002477921.1 | MNLLSGLFTKGISAIFEFIGNFSAQDICNAYFDEPEVPQELIDLQRK |
| | *agr*-II | WP_002461416.1 | MNLLSGLFTKGISVIFEFIGNFSVQDMCNGYFDEPEVPQELIDLHRN |
| *S. lutrae* | *agr*-I | WP_085236664.1 | MKFFDSILNFIVHFFQSIGNFAKIPFSFGFFDEPEIPEELLEEE |
| *S. massiliensis* | *agr*-I | WP_009384253.1 | MNIFSIFAKVLADVFSKIGTFGSIRICYGFFDEAEVPQELIDEANK |
| *S. microti* | *agr*-I | WP_084207673.1 | MNILDAIVKFFANLFKAIGNFGWINTCTHFFDEPEIPRELLELDK |
| *S. muscae* | *agr*-I | WP_095118143.1 | MNILDAIISFFSNIFKAIGNFGWINTCTHFFDEPEIPRELLELDK |
| *S. nepalensis* | *agr*-I | WP_103373157.1 | MNIFGSILTIFAKFFAFIGAISTVNPCGGYVDEPEVPKELTNLYE |
| | *agr*-II | WP_096808854.1 | MNIFESFLTFFAKFFATIGAISGVKPCTAFADEPEIPKELTDLYE |
| *S. pasteuri* | *agr*-I | WP_029056063.1 | METLVNLFFKFFTSIMEFVGLVAGANPCAGYFDEPEVPDELTKLYE |
| | *agr*-II | WP_017638097.1 | METLVNLFLKFFTSIMEFVGLVAGANPCAAYFDEPEVPEELSKLFE |
| | *agr*-III | WP_046467592.1 | METLVNLFLKFFTSIMEFVGLVAGANPCSAYFDEPEVPEELSQLFE |
| | *agr*-IV | WP_117238865.1 | METLVNLFLKFFTSIMEFVGFVAGYSPCANFFDEPEVPKELTQIYE |
| | *agr*-V | WP_119623149.1 | METLVNLFFKFFTSIMEFVGLVAGANPCAAFYDEPEVPAEITDLYK |
| *S. petrasii* | *agr*-I | WP_103297944.1 | MTFITNLFIKFFAFVLETVGTLASHTACATYFDEPEVPEELTNLKR |
| *S. pettenkoferi* | *agr*-I | WP_002471753.1 | MKLINIFFKILITSFSILGKQSGVYPCGGWWDEPEVPKEITDLHK |
| | *agr*-II | WP_124225583.1 | MFILNAIINFIIKLFSNFGGLAAKKGCGGMMDEPEVPNELIRNIKK |
| *S. piscifermentans* | *agr*-I | WP_095104393.1 | MDILNGIFKLFAFIFEQIGNIAKYNPCVGFFDEPEVPSELLDEQK |
| *S. pragensis* | *agr*-I | WP_126565907.1 | MTLITSLVIKLFTIILETIGTIATYTPCASYFDEPEVPEELTKIND |
| *S. pseudintermedius* | *agr*-I | WP_014613413.1 | MRILEVLFNLITNLFQSIGTFAKIPTSTGFFDEPEIPEELLEEDK |
| | *agr*-II | WP_019166405.1 | MRILEVLFNLITNLFQSIGTFARIPISTGFFDEPEIPAELLEEDK |
| | *agr*-III | WP_020219911.1 | MRILEVLFNLITNLFQSIGTFAKYPTSTGFFDEPEIPAELLEEDK |
| *S. rostri* | *agr*-I | WP_103358062.1 | MNILDAIVKFFANLFKAIGNFGWINTCTGFFDEPEIPRELLELDK |
| *S. saccharolyticus* | *agr*-I | WP_115312978.1 | MENIFNLFLKIFTTILEFIGTVAGDSVCSSYFDEPEVPEELSKLYE |
| *S. saprophyticus* | *agr*-I | WP_002482783.1 | MNVIKSISKSISKSISNYFAKVFASIGSISTINPCFGYTDESEIPKELTDLYE |
| | *agr*-II | WP_048793296.1 | MNFLNSIFSFFAKFFEVLGAVSGVNPCSGFFDEPEVPEELTKLYE |
| | *agr*-III | WP_017723435.1 | MNIFESLFNLIAKLFTVIGTIAGAKPCGGWLDEPEVPKEITDLYE |
| *S. schleiferi* | *agr*-I | WP_016425574.1 | MKFLDSLLKLLTNVFASISTYAKYPLCIGYFYEPEIPDELLEEEE |
| | *agr*-II | WP_050330469.1 | MDLLTKLIEFITGFFKSIGSFADIPLCRVFFDEPEIPRELLDLDE |
| *S. schweitzeri* | *agr*-I | WP_047427315.1 | MNTLLNIFFDFITGILKNIGNVASYSTCYFIMDEVEIPKELTQLHE |
| | *agr*-II | WP_047560140.1 | MKKLLNKVIELLVDFFNSIGYRAAYMNCDFLLDEAEVPKELTQIHE |
| *S. sciuri* | *agr*-I | WP_078099959.1 | MNLLSSLFSKSVSNILSAIGEKAVIRGCMAFLDETEVPAELLKEKQ |
| | *agr*-II | WP_107565491.1 | MNLLSSLFSKSVSNILSAIGEKAVIRGCNVFLDETEVPQELLNEKQ |

| | | | |
|---|---|---|---|
| | *agr*-III | WP_075578463.1 | MNLLSSLFSKSVSNILSAIGEKAVIRGCTAFLDETEVPAELLKEKQ |
| *S. simiae* | *agr*-I | WP_002464789.1 | MATLINLFLNLFTFIIERVGNVAAYSLCSLWYDEPEVPEELTKLHE |
| *S. simulans* | *agr*-I | WP_023015908.1 | MDLLNGIFKLFAFIFEKIGNLAKYNPCLGFLDEPTVPKELLEEDK |
| | *agr*-II | WP_107540291.1 | MDLLNGIFKLFAFIFEKIGNLAKYYPCWGYFDETEVPKELLEEDN |
| | *agr*-III | WP_002481721.1 | MDLLNGIFKLFAFIFEKIGNLAKYNPCWGYFDETEVPRELLEEDN |
| | *agr*-IV | AAL65848.1 | MELLNGIFKLFAFIFEKIGNLAKYYPCFGYFDESEVPQELLDEDK |
| *S. stepanovicii* | *agr*-I | WP_095086927.1 | MNILKGLFSKSISNVLAAIGGKSVIRGCNVFLDETEVPSELLNEKA |
| *S. succinus* | *agr*-I | WP_073504071.1 | MNILESLLTLITKFFSVLGATAGALPCGGFFDEPEVPSEITKLHE |
| | *agr*-II | WP_046838134.1 | MNIFESILNLFAKFFSVLGVIAGAKPCGGFFDEPEVPSEITKLYE |
| | *agr*-III | WP_069824787.1 | MNIFESILNLVAKFFTLLGTVAGAKPCGGWLDEPEVPSEITKLYE |
| *S. vitulinus* | *agr*-I | WP_016912307.1 | MNLLATLFSKSASNFLSSFGEKAVIRGCTAFLDETEVPAELLKEKQ |
| *S. warneri* | *agr*-I | WP_002466272.1 | MEFLVNLFFKFFTSIMEFVGFVAGYSPCTNFFDEPEVPSELTKIYE |
| | *agr*-II | WP_058709855.1 | MEFLVNLFFKFFTSILEFVGFVAGANPCAMFYDEPEVPSELTKLYE |
| | *agr*-III | WP_023373239.1 | MEFLFNFVLNIFTKIFETIGLVAGANPCVMYYDEPEVPEELSKIYE |
| | *agr*-IV | WP_124227772.1 | MEFLVNLFFKFFTSIMEFVGFVAGANPCSAYFDEPEVPKELTQQFE |
| *S. xylosus* | *agr*-I | WP_029378926.1 | MNIFTSILTIFVKFFSLIGTISSVRPCGGFVDEPEVPKELTKLYE |
| | *agr*-II | WP_069795725.1 | MNIFESILNFIAKFFAVIGAISGMKPCSGFVDEPEVPKELTNLYE |
| | *agr*-III | WP_042362360.1 | MNIIKSISKSISNYFAKVFASIGSISTINPCFGFTDESEIPKELTDLYE |
| | *agr*-IV | WP_069793020.1 | MFLLDSIFKIIAKFFTVIGVASGGHVCIGFADEIEVPKEITDLYE |
| | *agr*-V | WP_070052069.1 | MNFIKSISKSISNYFAKVFASIGSISTVNPCFGFTDESEIPKELTDLYE |

**Supplementary Table 4. List of AgrC sequences retrieved from NCBI.**

| Species (Accession) | AgrC sequence |
|---|---|
| *S. aureus agr*-I<br>(WP_001657119.1) | MELLNSYNFVLFVLTQMILMFTIPAIISGIKYSKLDYFFIIGISTLSLFLFKMFDSASLIILTSFIIIMYFVKI<br>KWYSILLIMTSQIILYCANYMYIVIYAYITKISDSIFVIFPSFFVVYVTISILFSYIINRVLKKISTPYLILNK<br>GFLIVISTILLLTFSLFFFYSQINSDEAKVIRQYSFIFIGITIFLSILTFVISQFLLKEMKYKRNQEEIETYYEY<br>TLKIEAINNEMRKFRHDYVNILTTLSEFIREDDMPGLRDYFNKNIVPMKDNLQMNAIKLNGIENLKVR<br>EIKGLITAKILRAQEMNIPISIEIPDEVSSINLNMIDLSRSIGIILDNAIEASTEIDDPIIRVAFIESENSVTFIV<br>MNKCADDIPRIHELFQESFSTKGEGRGLGLSTLKEIADNADNVLLDTIIENGFFIQKVEIINN |
| *S. aureus agr*-II<br>(WP_000447888.1) | METINNLAMATFQLVILFTVAKFISFVKFNLRDYFIIVGIIIPTMFLYYFYGSRAVLIPTFSSIIFLFFKLKY<br>YAIVTILVTMIIMYLSNFATVGLFLTLRKYTTDPAILLPLYILSFSSVSLLATYLVRISLKKFKKSYLSLN<br>KTYMIIISFVLFATFAFFYIYSTNTSSNGDSLIPYALVFIGLIIFISVVILIMSLFTLKEMKYKRNQEEIETY<br>YEYTLKIEAINNEMRKFRHDYVNILTTLSEYIREDDMIGLRAYFNKNIVPMKDNLQMNAIKLNGIENL<br>KVREIKGLITAKILRAQEMNIPISIEIPDEVSSINLNMIDLSRSIGIILDNAIEASTEIDDPIIRVAFIESENSV<br>TFIVMNKCADDIPRIHELFQESFSTKGEGRGLGLSTLKEIADNADNVLLDTIIENGFFIQKVEIINN |
| *S. aureus agr*-III<br>(WP_000387814.1) | MEALNDYNYVLFVIVQVSLMFFISAFISGIRYKKSDYIYIIGIVLSSVYFFDKIRSISLVVITIFIIIFLYFKIR<br>LYSVFLVMVTQIILYCANFVYIIIFSYIITISHSVFIVLPIFLVVYVSISYALAYILNRILKRINGTYLSLNKK<br>FLTVITIVIVITFSLLFAYSQIDASDASTIKQYSLLFLGIIILLSILIFIYSQFTLKEMKYKRNQEEIETYEYTL<br>KIEAINNEMRKFRHDYVNILTTLSEYIREDDMTGLRDYFNKNIVPMKDNLQMNALKLNGIENLKVREI<br>KGLLTAKILRAQEMNIPISIEIPDEVTRINLNMIDLSRSIGIILDNAIEASSEIDDPIIRVAFIESENSVTFIVM<br>NKCADDIPRIHELFQESFSTKGEGRGLGLSTLKEIADNADNVLLDTIIENGFFIQKVEIINN |
| *S. aureus agr*-IV<br>(WP_001807128.1) | MESLNSYNFVLFVLTQIILMFTVPSIISGIKYSKSDYLYITGITALSLILFNFIDSVTLIILTIFIIILYFSKIKW<br>YSILLIMTSQIILYCANYMYIVIFTYIVKIVDSIFVIFPIFFVVYVTISILFSYIINRVLKKISSSYLILNKGFLI<br>VISTILLLTFSLFFFYSQINSDEAKVIRQYSFIFIGITIFLSILTFVISQFLLKEMKYKRNQEEIETYYEYTLKI<br>EAINNEMRKFRHDYVNILTTLSEYIREDDMPGLRDYFNKNIVPMKDNLQMNAIKLNGIENLKVREIKG<br>LITAKILRAQEMSIPISIEIPDEVTHINLNMIDLSRSIGIILDNAIEASTEIDDPIIRVAFIESENSVTFIVMNK<br>CADDIPRIHELFQESFSTKGEGRGLGLSTLKEIADNADNVLLDTIIENGFFIQKVEIINN |
| *S. epidermidis agr*-I<br>(WP_002440376.1) | MDDINLFPFAGLQIFLMIWVTKVIINMKFNFRDYIIVFTIVIPSAIMYYFWQSKALIVLVIIIIIFFYTKIKL<br>YSILVVLFTTMILYITNFITVYIHLTIKDYIPFKFVLQLIHFTFFVIITLIIAYLTQLLFNKLKVSYLSLNKRY<br>LFIITIVLFISFILLYMVSQTDMRGNDTLKLYAILLMGIMVFLSVVILVMSNFTLREMRYKRNVKEIEAY<br>YEYTLRIESINNEMRKFRHDYVNILTTLSDYIREDDMPGLRKYFNENIVPMKDKLKTRSIKMNGIEKLK<br>VREIKGLITTKIIQAQEKRIPISIEVPDEIDRIDMNTVELSRIIGIIVDNAIEASENLEEPLINIAFIDNEEAVT<br>FIVMNKCSDDIPKIHELFEQGFSTKGDNRGLGLSTLKELTDSNENVLLDTVIENGYFVQKVEINNKES |
| *S. epidermidis agr*-II<br>(WP_002457064.1) | MGKLDFLPFAAIQVFLLVWVTKTIANIKFVRKDYIFITGIIILSAILYNVYASQALVLVVIMIIIFFYSKVR<br>WYSIVIVLMSTLLSYLTNFITVAISLYTENIIHNIYFYNIFHFSIFIILSLILAHLFKHLLIRFRYSYLYLSKR<br>YYIIISFVLAIAFIYFYIISQTNLQESNSLNFYAIIFVSITVLLSLVILLLSAFALREMKYKRKLQEIEAYYE<br>YTLRIESINNEMRKFRHDYVNILTTLSDYIREDDMPGLRKYFNENIVPMKDKLKTRSIKMNGIEKLKVR<br>EIKGLITTKIIQAQEKRIPISIEVPDEIDRISMNTVELSRIIGIIVDNAIEASENLEEPLINIAFIDNEESVTFIV<br>MNKCSDDIPKIHELFEQGFSTKGDNRGLGLSTLKELTDSNENVLLDTVIENGYFVQKVEINNKES |
| *S. epidermidis agr*-III<br>(WP_020367417.1) | MDELNFLSFAAIQIFLLVWVIKTIANIKFVRKDYIFITGIIILSAILYNVYSSQALVLVVLMIIIFFYSKVR<br>WYSIVIVLMSTLLSYLTNFITVVISLYTEDIIHNIYLFTIFHLLIYVILSLILAHLFKHLLIKLRYIYLYISKR<br>YYFIISFVLAIAFIYFYIISQTNLQENNSLKFYAIIFVSVIVFLSLVILLLSAFALREMKYKRKLQEIEAYYE<br>YTLRIESINNEMRKFRHDYVNILTTLSDYIREDDMPGLRKYFDEHIVPMKDKLKTRSIKMNGIEKLKVR<br>EIKGLITTKIIQAQEKRIPISIEVPDEIDRIDMNTVELSRIIGIIVDNAIEASENLEEPLINIAFIDNDESVTFIV<br>MNKCSDDIPKIHELFEQGFSTKGDNRGLGLSTLKELTDSNENVLLDTVIENGYFVQKVEINNKES |
| *S. lugdunensis agr*-I<br>(WP_002477920.1) | MDLLNSIPLSIFQSLMLLFVAKIVADIKFQMRDYFAIFGIIIPSTILFGVIGRQSLIFLIIGCLIFFYLKIGLYS<br>VLAIFGSALIMYVSNYISVILSVIADYFSLSYIVQIIIILVSFTLISIICAYFIRFLINKLKKTYLYFNKIYISVI<br>SIFLILSLIMLYLYTQIFKQEYQDLKIFAIIFVGILFFLAIFIIVITFSVHREMQYKRNLKEIETYYEYTLQIE<br>SINNEMRKFRHDYVNILSTMSEYIREDDMPGLREYFNNNIVSMKDNLQMNSIKINGTDKLKVRAIKGL<br>VTTKILQAQEKNIPISIEVPELIEHIEMNTVDLSRVIGIIIDNAIEASESLEDALIRIAFIKNEDSVLFIVMNK<br>CSDTTPKIHELFQENFSTKGKNRGLGLSNLKEITDATPNTLLDTTIDNGYFIQKVEILNNIP |

#### MRSA mouse skin infection model data

**Supplementary Table 5. Skin lesion size, colony-forming unit (CFU) values and body weight for MRSA mouse skin infection model.**

| Treatment | Mouse | Skin lesion in mm$^2$ | | | $\log_{10}$ CFU | | | Body weight in g | | |
|---|---|---|---|---|---|---|---|---|---|---|
| | | Day 1 | Day 2 | Day 4 | Day 1 | Day 2 | Day 4 | Day 1 | Day 2 | Day 4 |
| vehicle | 1 | 71.0 | - | - | 7.68 | - | - | 18.6 | - | - |
| | 2 | 76.2 | - | - | 7.89 | - | - | 21.4 | - | - |
| | 3 | 62.3 | - | - | 7.98 | - | - | 20.8 | - | - |
| | 4 | 69.9 | - | - | 8.11 | - | - | 19.6 | - | - |
| | 5 | 88.1 | - | - | 8.06 | - | - | 19.1 | - | - |
| | 6 | 69.7 | - | - | 7.95 | - | - | 18.9 | - | - |
| | 7 | 74.3 | - | - | 7.74 | - | - | 18.0 | - | - |
| | 8 | 79.4 | - | - | 7.68 | - | - | 18.6 | - | - |
| | 9 | 63.4 | 98.3 | - | - | 7.38 | - | 18.6 | 20.6 | - |
| | 10 | 70.5 | 99.3 | - | - | 7.89 | - | 20.8 | 20.4 | - |
| | 11 | 61.8 | 70.3 | - | - | 7.49 | - | 18.7 | 20.4 | - |
| | 12 | 52.8 | 101.3 | - | - | 7.56 | - | 20.4 | 19.0 | - |
| | 13 | 82.2 | 103.7 | 64.4 | - | - | 7.11 | 19.3 | 19.9 | 19.6 |
| | 14 | 73.4 | 96.5 | 58.8 | - | - | 7.76 | 19.9 | 18.8 | 18.6 |
| | 15 | 60.4 | 96.3 | 74.7 | - | - | 7.24 | 20.5 | 18.6 | 18.6 |
| | 16 | 71.0 | 97.5 | 58.2 | - | - | 6.19 | 19.4 | 20.9 | 21.2 |
| 2% fusidic acid ointment | 1 | 64.3 | 76.7 | - | - | 7.27 | - | 18.8 | 18.3 | - |
| | 2 | 82.5 | 98.4 | - | - | 6.89 | - | 18.8 | 20.5 | - |
| | 3 | 69.1 | 69.7 | - | - | 7.18 | - | 18.9 | 20.7 | - |
| | 4 | 47.4 | 66.4 | - | - | 7.38 | - | 20.4 | 19.1 | - |
| | 5 | 55.2 | 80.5 | 38.2 | - | - | 4.12 | 19.2 | 18.9 | 19.5 |
| | 6 | 74.9 | 81.4 | 53.9 | - | - | 5.70 | 19.1 | 20.2 | 21.4 |
| | 7 | 69 | 88.1 | 41 | - | - | 6.12 | 19.9 | 18.8 | 19.0 |
| | 8 | 63.3 | 70.1 | 56.5 | - | - | 6.04 | 19.3 | 19.6 | 19.8 |
| *Ss*-AIP-II (**24**) | 1 | 75.5 | 84.7 | - | - | 7.88 | - | 19.1 | 20.0 | - |
| | 2 | 77.6 | 91.5 | - | - | 7.95 | - | 20.7 | 19.3 | - |
| | 3 | 111.4 | 81.3 | - | - | 8.08 | - | 19.0 | 20.2 | - |
| | 4 | 79.6 | 81.5 | - | - | 7.51 | - | 18.5 | 20.7 | - |
| | 5 | 63.3 | 94.1 | 47.7 | - | - | 5.68 | 19.5 | 21.7 | 22.0 |
| | 6 | 64.6 | 77.4 | 41.9 | - | - | 5.19 | 20.3 | 19.3 | 19.8 |
| | 7 | 51 | 61 | 50 | - | - | 5.30 | 21.3 | 19.1 | 19.4 |
| | 8 | 64.5 | 69 | 52.6 | - | - | 5.04 | 19.7 | 18.6 | 18.8 |

#### Bacterial strains and *agrD* sequencing

**Supplementary Table 6. Bacterial strains used in this study.**

| Name | Characterisitics | Reference |
|---|---|---|
| P3-*blaZ*/*pagrC*-I | *S. aureus* RN10829 reporter expressing AgrC-I | 16 |
| P3-*blaZ*/*pagrC*-II | *S. aureus* RN10829 reporter expressing AgrC-II | 16 |
| P3-*blaZ*/*pagrC*-III | *S. aureus* RN10829 reporter expressing AgrC-III | 16 |
| P3-*blaZ*/*pagrC*-IV | *S. aureus* RN10829 reporter expressing AgrC-IV | 16 |
| AH3408 | *S. epidermidis* ATCC1228(AH1740) + pCM40 (ermR, arP3_sGFP, type-I) | 15 |
| AH3623 | *S. epidermidis* 1457(AH1738) ica::DHFR + pCM40 (ermR, arP3_sGFP, type-II) | 15 |
| AH3409 | *S. epidermidis* Fey 8247+ pCM40 (ermR, arP3_sGFP, type-III) | 15 |
| AH4031 | *S. lugdunensis* N920143+ pCM40 (ermR, arP3_sGFP, type-I) | 17 |
| AH1677 | *S. aureus agr*-I P3-dependent YFP expression | 18 |
| AH430 | *S. aureus agr*-II P3-dependent YFP expression | 18 |
| AH1747 | *S. aureus agr*-III P3-dependent YFP expression | 18 |
| AH1872 | *S. aureus agr*-IV P3-dependent YFP expression | 18 |
| 5927201 | *S. pasteuri* human nose isolate | This study |
| 102 L88 | *S. succinus* horse isolate | This study |
| 2934 c-30869-41 | *S. cohnii* isolate from skim milk | This study |

**Supplementary Table 7. Primer used for *agrD* sequencing.**

| Species | Forward primer | Reverse primer |
|---|---|---|
| *S. pasteuri* | ccattgttagataaaaacttacagcc | gaaaaggaaacgaatactctccattg |
| *S. cohnii* | ccgtctgaaataatgtataccttgtc | ttatgtctacaattagactacttaaccac |
| *S. succinus* | gtcaaagtagcaagccatgtag | cattagcaatcattggctttgttg |

**Supplementary Table 8. AgrD sequences used for AIP identification.**

| Species | AgrD sequence |
|---|---|
| *S. pasteuri* | METLVNLFFKFFTSIMEFVGLVAGANPCAGYFDEPEVPDELTKLYE |
| *S. cohnii* | MNIFESILTIFAKFFTFIGTISSVKPCTGFVDEPEIPKELTDLYE |
| *S. succinus* | MTILESLLTLITKFFSVLGATAGALPCGGFFDEPEVPSEITKLHE |

#### 3. Synthetic procedures

**Synthesis of Fmoc-Cys(STmp)-OH (S3)**

The protected amino acid was synthesized according to a literature procedure.[19]

**2,4,6-Trimethoxybenzenethiol (S1).** To a solution of 2,4,6-trimethoxybenzene (3.36 g, 20.0 mmol, 1.00 equiv) in anhydrous THF (25 mL) under argon atmosphere at 0 °C was added *n*-butyllithium in hexane (1.2 M, 16.7 mL, 20.0 mmol, 1.00 equiv) and *N*,*N*,*N'*,*N'*-tetramethylethylendiamine (0.25 mL, 1.68 mmol, 0.08 equiv) subsequently. The reaction mixture was allowed to warm to room temperature and stirred for 1 h. A solution of elemental sulfur (0.58 g, 18 mmol, 0.90 equiv) in toluene (15 mL) was added dropwise and the reaction mixture was stirred at room temperature. After 6 h, water (50 mL) was added, followed by aqueous HCl (1.0 M) (40 mL) and the aqueous layer was extracted with $CH_2Cl_2$ (3 × 25 mL). The combined organic layer was washed with water (3 × 25 mL), brine (25 mL), dried over $MgSO_4$, filtered and concentrated under reduced pressure. The resulting yellow oil was cyrstallized from heptane with a small amount of $CH_2Cl_2$ to yield **S1** as yellow crystaline solid (1.3 g, 32%). $^1$H NMR (600 MHz, $CDCl_3$) $\delta$ = 3.76 (s, 1H), 3.80 (s, 3H), 3.87 (s, 6H), 6.18 (s, 2H).

**Fmoc-Cys-OH (S2).** To a solution of Fmoc-Cys(Trt)-OH (5.85 g, 10.0 mmol, 1.00 equiv) in $CH_2Cl_2$ (30 mL) was added triisopropylsilane (2.04 mL, 10.0 mmol, 1.00 equiv) followed by TFA (7.70 mL, 100 mmol, 10.0 equiv). The reaction mixture was stirred for 15 min at room temperature and subsequently concentrated under reduced pressure with toluene co-evaporation. The remaining residue was suspended in heptane, transferred to centrifugal tubes, centrifuged for 5 min and the heptane decanted. The procedure was repeated 5 times and the pellet was dried under reduced pressure to yield the **S2** as white solid (2.9 g, 85%). $^1$H NMR (600 MHz, DMSO-$d_6$) $\delta$ = 2.68–2.81 (m, 1H), 2.84–2.95 (m, 1H), 4.13 (td, *J* = 8.4, 4.4 Hz, 1H), 4.24 (t, *J* = 7.0 Hz, 1H), 4.28–4.37 (m, 2H), 7.28–7.38 (m, 2H), 7.42 (t, *J* = 7.4 Hz, 2H), 7.68 (d, *J* = 8.3 Hz, 1H), 7.74 (d, *J* = 7.5 Hz, 2H), 7.89 (d, *J* = 7.5 Hz, 2H), 12.86 (br s, 1H).

**Fmoc-Cys(STmp)-OH (S3).** A solution of **S1** (500 mg, 2.50 mmol, 1.00 equiv) and **S2** (858 mg, 2.50 mmol, 1.00 equiv) in anhydrous THF (5.0 mL) was added dropwise under argon atmosphere to a suspension of *N*-chlorosuccinimide (351 mg, 2.63 mmol, 1.05 equiv) in anhydrous $CH_2Cl_2$ (4.0 mL) at -78 °C under exlusion of light. The reaction mixture was stirred for 2 h at -78 °C, subsequently warmed to room temperature and diluted with $CH_2Cl_2$ (20 mL). The organic phase was washed with aqueous HCl (1.0 M) (3 × 20 mL), water (1 × 20 mL) and brine (3 × 20 mL), dried over $MgSO_4$, filtered and concentred under reduced pressure. The crude product was purified by flash column chromatography ($CH_2Cl_2$/MeOH = 97/3 + 0.25% AcOH) to yield the **S3** as off-white foam (960 mg, 71%). $^1$H NMR (600 MHz, DMSO-$d_6$) $\delta$ = 2.90 (dd, *J* = 13.5, 10.4 Hz, 1H), 3.26 (dd, *J* = 13.5, 4.2 Hz, 1H), 3.79 (s, 3H), 3.80 (s, 6H), 4.16–4.28 (m, 2H), 4.26–4.37 (m, 1H), 4.48 (ddd, *J* = 10.4, 8.3, 4.2 Hz, 1H), 6.25 (s, 2H), 7.33 (t, *J* = 7.4 Hz, 2H), 7.43 (t, *J* = 7.5 Hz, 2H), 7.67–7.73 (m, 3H), 7.90 (d, *J* = 7.5 Hz, 2H), 12.80 (s, 1H).

##### Preparation of NCL trapping resins

Amino PEGA resin (1.00 g, loading: 0.42 mmol/g, 0.42 mmol) was placed in a polypropylene syringe equipped with a fritted disk, swelled in DMF for 15 min and washed with DMF (5 × 1 min). Fmoc-Rink-amide linker (1.13 g, 2.10 mmol, 5.00 equiv), HATU (782 mg, 2.06 mmol, 4.90 equiv), and *i*-$Pr_2NEt$ (736 µL, 4.20 mmol, 10.0 equiv) were pre-incubated in DMF (10.0 mL) for 2 min and then added to the resin. After 2 h, the resin was washed with DMF (3 × 1 min), MeOH (3 × 1 min), and $CH_2Cl_2$ (3 × 1 min) and treated with a capping solution ($Ac_2O$–*i*-$Pr_2NEt$–$CH_2Cl_2$, 2:2:6, v/v/v, 10.0 mL). After 2 h, the resin was washed with DMF (3 × 1 min), MeOH (3 × 1 min), and $CH_2Cl_2$ (3 × 1 min). The resin was then treated with piperidine in DMF (1:4, v/v, 10.0 mL) (1 × 2 min, 1 × 20 min) and washed with DMF (3 × 1 min), MeOH (3 × 1 min), and $CH_2Cl_2$ (3 × 1 min). Fmoc-Cys(S*t*-Bu)-OH (272 mg, 0.63 mmol, 1.50 equiv) or Fmoc-Cys(STmp)-OH (**S3**) (335 mg, 0.63 mmol, 1.50 equiv), HATU (240 mg, 0.63 mmol, 1.50 equiv), and *i*-$Pr_2NEt$ (220 µL, 1.26 mmol, 3.00 equiv) were pre-incubated in DMF (10.0 mL) for 2 min and then added to the resin. After 2 h, the resin was washed with DMF (3 × 1 min), MeOH (3 × 1 min), and $CH_2Cl_2$ (3 × 1 min) and dried under high vacuum for 16 h.

##### Synthesis of thiolactone-containing AIPs using the 3-4-amino-(methylamino)benzoic acid (MeDbz) linker

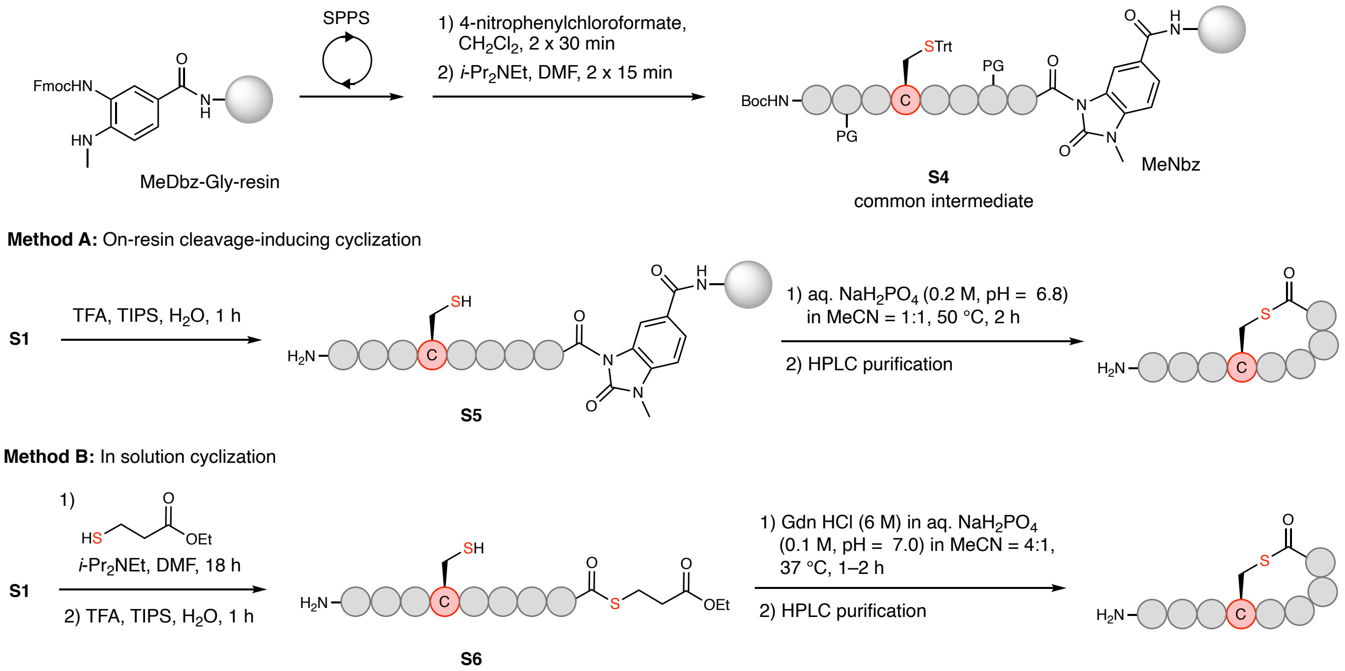

**Supplementary Figure 30. Synthesis of thiolactone-containing AIPs using the MeDbz linker.** *N*-acyl-benzimidazolinone (Nbz) intermediates **S4** were synthesized on MeDbz-Gly resin and used for method A and B. In method A, **S4** was used for an on-resin cleavage-inducing cyclization protocol and in method B **S1** was treated with a thiol to induce a cleavage-inducing thioesterifiaction followed by a chemoselective cyclization in solution.

**General protocol for automated peptide synthesis**

Automated peptide synthesis was carried out on a Biotage SyroWave™ synthesizer using standard Fmoc SPPS chemistry. The following Fmoc-protected amino acids with side chain protecting groups were used: Fmoc-Ala-OH, Fmoc-Asn(Trt)-OH, Fmoc-Cys(Trt)-OH, Fmoc-Cys(*S*t-Bu)-OH, Fmoc-Gly-OH, Fmoc-Ile-OH, Fmoc-Leu-OH, Fmoc-Phe-OH, Fmoc-Pro-OH, Fmoc-Ser(*t*-Bu)-OH, Fmoc-Ser(TBDMS)-OH, Fmoc-Thr(*t*-Bu)-OH, Fmoc-Trp(Boc)-OH, Fmoc-Tyr(*t*-Bu)-OH, and Fmoc-Val-OH. The following Boc-protected amino acids with side chain protecting groups were used as *N*-terminal amino acids: Boc-Ala-OH, Boc-Asn(Trt)-OH, Boc-Asp(O*t*-Bu)-OH, Boc-Arg(Pbf)-OH, Boc-Lys(Boc)-OH, Boc-Ser(*t*-Bu)-OH, Boc-Thr(*t*-Bu)-OH and Boc-Tyr(*t*-Bu)-OH.

SPPS was performed on 0.02 mmol (method A) or 0.04 mmol (method B) scale using MeDbz-Gly-ChemMatrix resin[7] or on 0.04 mmol scale using preloaded chlorotrityl (Cl-Trt) resin. Fmoc deprotection was performed in two stages: 1) piperidine in DMF (2:3, v/v) for 3 min and 2) piperidine in DMF (1:4, v/v) for 12 min. The deprotection was followed by washing with DMF (2 × 45 s), $CH_2Cl_2$ (1 × 45 s), and DMF (2 × 45 s). The first coupling reaction on MeDbz-Gly-ChemMatrix resin was performed as double coupling using Fmoc-AA-OH (5.00 equiv to the resin loading), HATU (4.90 equiv) and *i*-$Pr_2$NEt in NMP (10.0 equiv, 2.0 M) in DMF (final concentration = 0.2 M for Fmoc-AA-OH) for 90 min. Standard coupling reactions were performed as double couplings with Fmoc-AA-OH (5.00 equiv to the resin loading), HBTU (4.90 equiv) and *i*-$Pr_2$NEt in NMP (10.0 equiv, 2.0 M) in DMF (final concentration = 0.2 M for Fmoc-AA-OH) for 40 min for each coupling. The last amino acid was incorporated as Boc-AA-OH.

**General procedure for *N*-acyl-benzimidazolinone (Nbz) formation**

After automated peptide elongation, the peptidyl-MeDbz-Gly-ChemMatrix resin (1.00 equiv) was transferred into a polypropylene syringe equipped with a fritted disk using $CH_2Cl_2$ and the resin was then washed with $CH_2Cl_2$ (5 × 1 min). A solution of 4-nitrophenyl-chloroformate (5.00 equiv) in $CH_2Cl_2$ (concentration = 0.1 M) was added to the resin and the suspension was agitated for 30 min. The resin was then washed with $CH_2Cl_2$ (2 × 1 min) and the procedure was repeated. The resin was then washed with $CH_2Cl_2$ (3 × 1 min) and DMF (3 × 1 min) and a solution of *i*-$Pr_2$NEt (25.0 equiv) in DMF (0.5 M) was added to the resin. After 15 min, the resin was washed with DMF (3 × 1 min) and the procedure was repeated. The resin was then washed with DMF (3 × 1 min), *i*-$Pr_2$NEt in DMF (5%, v/v) (3 × 1 min), DMF (3 × 1 min), MeOH (3 × 1 min), and $CH_2Cl_2$ (3 × 1 min) and dried under high vacuum.

**Method A: General procedure for on-resin clevage-inducing cyclization**

Dried peptidyl-MeNbz-Gly-ChemMatrix resin **S4** (0.02 mmol, 1.00 equiv) was placed in a polypropylene syringe equipped with a fritted disk and treated with a deprotection/cleavage cocktail (2.0 mL, TFA–*i*-$Pr_3SiH$–water, 94:3:3, v/v/v) for 1 h and the TFA cocktail removed from the resin. The resin **S5** was washed with $CH_2Cl_2$ (3 × 1 min), DMF (3 × 1 min), and $CH_2Cl_2$ (3 × 1 min) and dried under suction for several minutes. The cyclization buffer (5.0 mL, phosphate buffer (0.2 M, pH = 6.8)–MeCN, 1:1, v/v) was added to the resin (final concentration = 4.0 mM) and the suspension was agitated at 50 °C for 2 h. The solution was removed from the resin, collected and the resin was rinsed with fresh cyclisation buffer. The combined peptide-containing soltuion was purified by preparative RP-HPLC. Fractions containing pure peptide were lyophilized to afford the desired cyclized peptide.

**Method B: General procedure for solution cyclization**

The procedure was adapted from a previously published protocol.[20] Dried peptidyl-MeNbz-Gly-ChemMatrix resin **S4** (0.04 mmol, 1.00 equiv) was placed in a polypropylene syringe equipped with a fritted disk and treated with a solution of 3-mercaptopropionic acid ethyl ester (50.6 μL, 0.40 mmol, 10.0 equiv) and *i*-$Pr_2NEt$ (69.8 μL, 0.40 mmol, 10.0 equiv) in DMF (3.0 mL). After overnight incubation, the peptide containing solution was removed from the resin and the resin washed twice with DMF (2.0 mL). The combined organic phase was concentrated to dryness under pressure and the remaining residue was treated with a deprotection cocktail (2.0 mL, TFA–*i*-$Pr_3SiH$–water, 94:3:3, v/v/v) for 2 h. The reaction mixture was concentrated under a stream of nitrogen and precipitated by addition of ice-cold diethyl ether. The crude peptide thioester **S6** was lyophilized and used in the next step without further purification.

Crude peptide thioester **S3** (10.0 μmol) was dissolved in cyclization buffer (10.0 mL, guanidinium hydrochloride (6 M) in phosphate buffer (0.1 M, pH = 7.0)–MeCN, 4:1, v/v) with a final concentration of 1.0 mM and the reaction mixture was agitated at 37 °C. After for 2 h, the reaction mixture was purified by preparative RP-HPLC. Fractions containing pure peptide were lyophilized to afford the desired cyclized peptide.

#### Alternative synthesis of *S. lugdunensis* AIP-I (8)

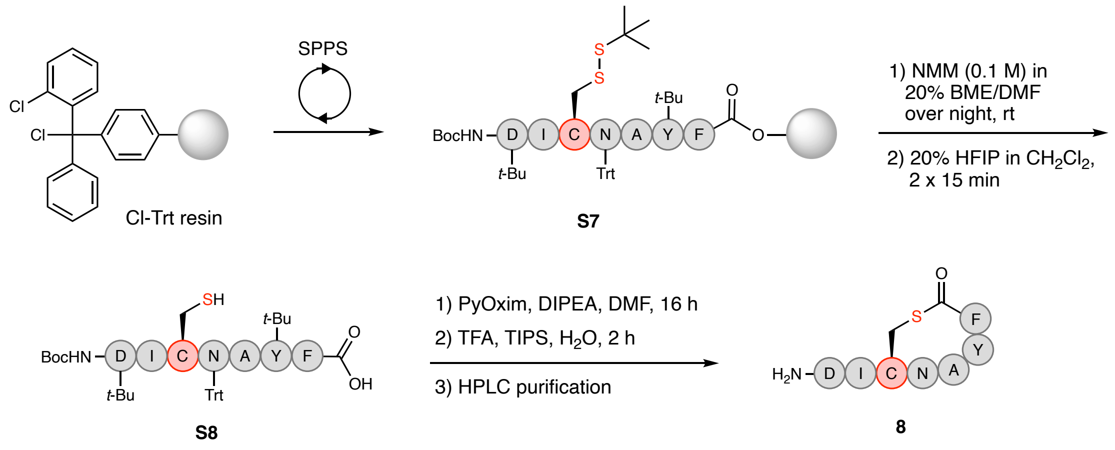

**Supplementary Figure 31. Synthesis of *S. lugdunensis* AIP-I (8) via PyOxim-mediated thiolactonization.** The linear peptide **S7** was synthesized on Cl-Trt resin and the cysteine side chain protecting group removed on-resin. The partially-protected peptide **S8** was released from the resin and the peptide cyclized via PyOxim-mediated thiolactonization.

The fully protected linear peptide **S7** was synthesized in 40.0 μmol scale on Cl-Trt polystyrene resin preloaded with Fmoc-Phe-OH (0.69 mmol/g) using the general procedures for automated SPPS. After completed peptide elongation, the resin **S7** was transferred into a polypropylene syringe equipped with a fritted disk using $CH_2Cl_2$ and washed with DMF (3 × 1 min). A solution of *N*-methylmorpholine (NMM) (22.1 μL, 0.20 mmol, final concentration = 0.1 M) in a mixture of *β*-mercaptoethanol (BME)–DMF (2.0 mL, 1:4, v/v) was added to the peptidyl resin **S7** and the resin was agitated overnight at room temperature. The next day, the thiol-containing solution was removed by suction and the resin was washed with DMF (3 × 1 min), MeOH (3 × 1 min) and $CH_2Cl_2$ (3 × 1 min) and dried under suction for 15 min. The dried resin was treated with a solution of hexafluoroisopropanol (HFIP) (0.4 mL) in $CH_2Cl_2$ (1.6 mL) for 15 min at room temperature to cleave the partially-protected peptide **S8** from the resin. The cleavage solution was removed from the resin, collected and a fresh HFIP–$CH_2Cl_2$ solution (2.0 mL, 1:4, v/v) was added to the resin. After 15 min, the cleavage solution was removed from the resin, collected and the resin rinsed with $CH_2Cl_2$ (2.0 mL). The combined organic phase was concentrated to dryness under reduced pressure to yield the partially protected peptide **S8**, which was used without further purification.

The crude partially-protected peptide **S8** (0.04 mmol based on the resin loading) was dissolved in anhydrous DMF (5.0 mL) under nitrogen atmosphere and added dropwise to a solution of PyOxim (42.2 mg, 0.08 mmol, 2.00 equiv) and *i*-$Pr_2NEt$ (48.6 μL, 0.28 mmol, 7.00 equiv) in anhydrous DMF (35.0 mL). The reaction mixture was stirred overnight at room temperature and after full consumption of **S8** was confirmed by UPLC-MS, the reaction was reduced to dryness under reduced pressure. The remaining residue was treated with a deprotection cocktail (3.0 mL, TFA–*i*-$Pr_3SiH$–water, 94:3:3, v/v/v) for 2 h and subsequently concentrated under a stream of nitrogen followed by precipitation in ice-cold diethyl ether. The crude peptide **8** was purified by preparative RP-HPLC. Fractions containing pure peptide were lyophilized to afford the desired cyclized peptide. Full characterization of *S. lugdunensis* AIP-I (**8**) in section 4.

#### Synthesis of *S. intermedius* AIP-I (22)

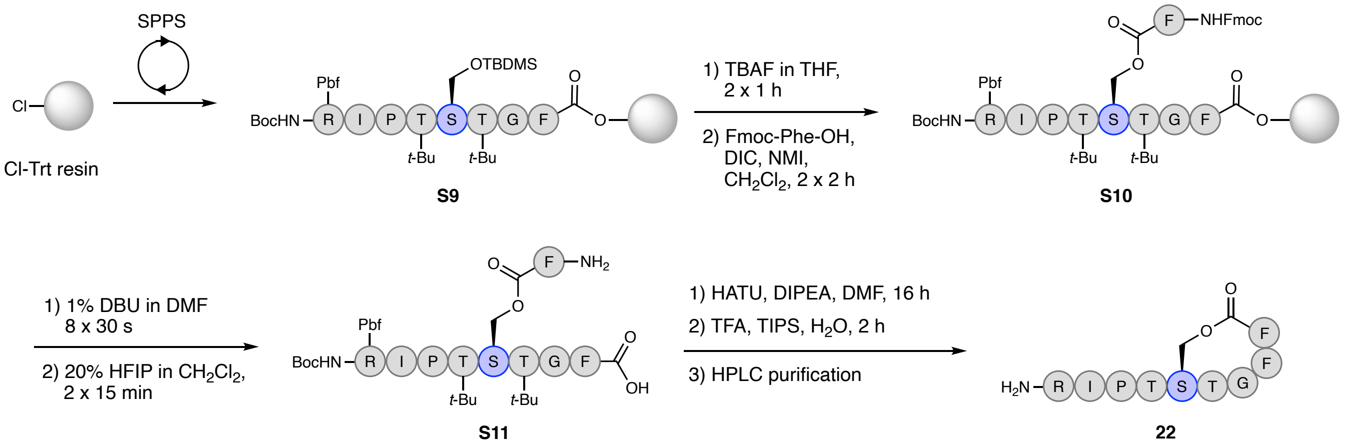

**Supplementary Figure 32. Synthesis of *S. intermedius* AIP-I (22) using an on-resin esterification strategy.** The linear peptide **S9** was synthesized on Cl-Trt resin and the serine side chain protecting group was removed on-resin followed by esterification to give **S10**. The partially-protected peptide **S11** was released from the resin and the peptide cyclized via HATU-mediated amide coupling.

The fully protected linear peptide **S6** was synthesized in 40.0 µmol scale on Cl-Trt polystyrene resin preloaded with Fmoc-Phe-OH (0.69 mmol/g) using the general procedures for automated SPPS. After completed peptide elongation, the peptidyl-resin **S6** was transferred into a polypropylene syringe equipped with a fritted disk using $CH_2Cl_2$, washed with $CH_2Cl_2$ (3 × 1 min) and dried overnight under vacuum.

The peptidyl-resin **S6** was swelled in anhydrous THF (1.5 mL) for 15 min and subsequently a solution of tetrabutylammonium fluoride (TBAF) in THF (1.0 M) (0.40 mL, 0.40 mmol, 10.0 equiv) in anhydrous THF (4.6 mL) was added to the resin and the resin was agitated at room temperature. After 1 h, the TBAF solution was removed by suction and resin treated with a fresh TBAF–THF solution. After 1 h, the TBAF solution was removed by suction and the resin washed with DMF (3 × 1 min), MeOH (3 × 1 min) and $CH_2Cl_2$ (3 × 1 min) and dried overnight under vacuum.

On-resin esterification was performed according to a previously published protocol.[21] The resin was swelled in anhydrous $CH_2Cl_2$ (1.5 mL) for 15 min and subsequently a solution of Fmoc-Phe-OH (93.0 mg, 0.24 mmol, 6.00 equiv), *N*,*N*'-diisopropylcarbodiimide (DIC) (37.6 µL, 0.24 mmol, 6.00 equiv), *N*-methylimidazole (NMI) (17.2 µL, 0.22 mmol, 5.40 equiv). in anhydrous $CH_2Cl_2$ (1.5 mL) was added. The resin was agitated at room temperature for 2 h and subsequently washed with anhydrous $CH_2Cl_2$ (3 × 1 min). A fresh Fmoc-Phe-OH–DIC–NMI solution was added to resin and after 2 h of incubation, the resin was washed with DMF (3 × 1 min), $CH_2Cl_2$ (3 × 1 min) and DMF (5 × 1 min).

Fmoc-removal of the *O*-acylated peptidyl resin **S7** was performed by treatment of the resin with a solution of 1,8-biazabicyclo[5.4.0]undec-7-ene (DBU) in DMF (1.5 mL, 1:99, v/v) (8 × 30 s). The resin was subsequently washed with DMF (3 × 1 min), MeOH (3 × 1 min) and $CH_2Cl_2$ (3 × 1 min) and dried under suction for 15 min. The dried resin was treated with a solution of hexafluoroisopropanol (HFIP) (0.4 mL) in $CH_2Cl_2$ (1.6 mL) for 15 min at room temperature to cleave the partially protected peptide **S8** from the resin. The cleavage solution was removed from the resin and collected and a fresh HFIP–$CH_2Cl_2$ solution was added to the resin. After 15 min the cleavage solution was removed from the resin and collected and the resin rinsed with $CH_2Cl_2$ (2.0 mL). The combined cleavage solutions and the rinsing solution were evaporated to dryness under reduced pressure to yield the partially protected peptide **S8,** which was used without further purification.

The crude partially-protected peptide **S8** (0.04 mmol based on the resin loading) was dissolved in anhydrous DMF (5.0 mL) under nitrogen atmosphere and added dropwise to a solution of HATU (15.2 mg, 0.04 mmol, 1.00 equiv) and *i*-$Pr_2NEt$ (20.8 µL, 0.12 mmol, 3.00 equiv) in anhydrous DMF (35.0 mL). The reaction mixture was stirred overnight at room temperature and after full consumption of **S8** was confirmed by UPLC-MS, the reaction was reduced to dryness under reduced pressure. The remaining residue was treated with a deprotection cocktail (3.0 mL, TFA–*i*-$Pr_3SiH$–water, 94:3:3, v/v/v) for 2 h and subsequently concentrated under a stream of nitrogen followed by precipitation in ice-cold diethyl ether. The crude peptide **22** was purified by preparative RP-HPLC. Fractions containing pure peptide were lyophilized to afford the desired cyclized peptide. Full characterization of *S. intermedius* AIP-I (**22**) in section 4.

### 4. Characterization of synthetic AIPs

##### *S. epidermidis* AIP-I (5)

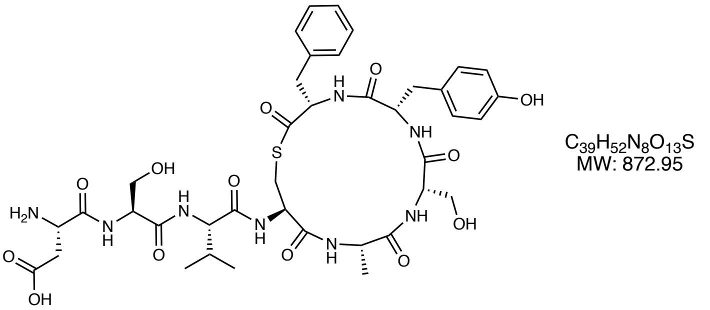

The peptide was synthesized according to general method B for solution cyclization (section 4). Purification by preparative RP-HPLC (5–40% B over 40 min, C18) afforded *S. epidermidis* AIP-I (**5**) as a fluffy white solid after lyophilization (5.0 mg, 50% based on 10.0 µmol linear thioester peptide). Purity 99% determined by UPLC ($\lambda$ = 215 nm). UPLC-MS (ESI) *m/z* calcd for $[M+H]^+$ $C_{39}H_{53}N_8O_{13}S^+$: 873.35, found 873.32. $^1$H NMR (600 MHz, DMSO-$d_6$) $\delta$ = 0.83 (t, $J$ = 7.1 Hz, 6H), 1.21 (d, $J$ = 7.3 Hz, 3H), 1.95–2.04 (m, 1H), 2.58–2.67 (m, 2H), 2.67–2.72 (m, 1H), 2.73–2.82 (m, 2H), 2.90–2.95 (m, 1H), 3.16 (dd, $J$ = 12.8, 3.8 Hz, 1H), 3.28 (dd, $J$ = 14.0, 3.9 Hz, 1H), 3.54–3.66 (m, 4H), 4.05–4.07 (m, 2H), 4.15–4.23 (m, 3H), 4.24–4.30 (m, 2H), 4.41–4.46 (m, 1H), 5.04–5.11 (m, 1H), 5.13–5.20 (m, 1H), 6.62 (d, $J$ = 8.5 Hz, 2H), 6.85 (d, $J$ = 8.5 Hz, 2H), 7.01–7.04 (m, 2H), 7.16–7.20 (m, 1H), 7.21–7.24 (m, 2H), 7.83 (d, $J$ = 8.7 Hz, 1H), 7.91 (d, $J$ = 8.8 Hz, 1H), 8.04 (d, $J$ = 7.5 Hz, 1H), 8.21 (d, $J$ = 8.3 Hz, 1H), 8.45 (d, $J$ = 7.8 Hz, 1H), 8.58 (d, $J$ = 7.6 Hz, 1H), 8.88 (d, $J$ = 8.2 Hz, 1H), 9.22 (s, 1H).

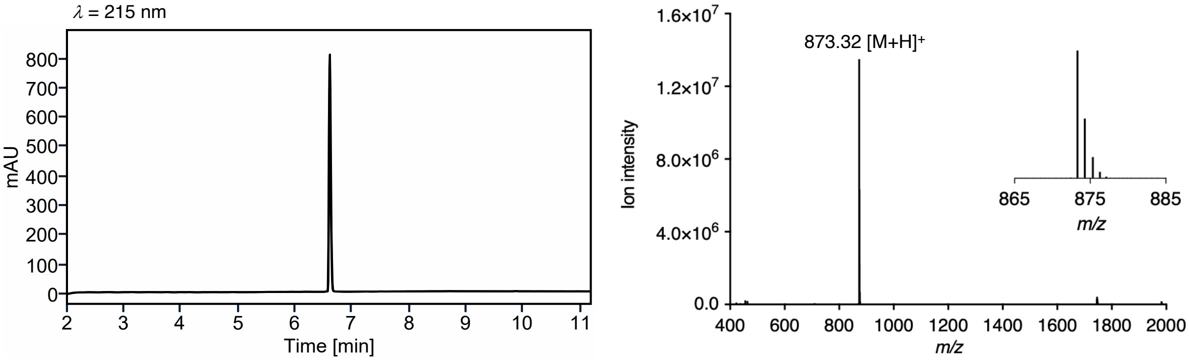

**Supplementary Figure 33. UPLC trace and MS spectrum of *S. epidermidis* AIP-I (5).**

##### *S. epidermidis* AIP-II (6)

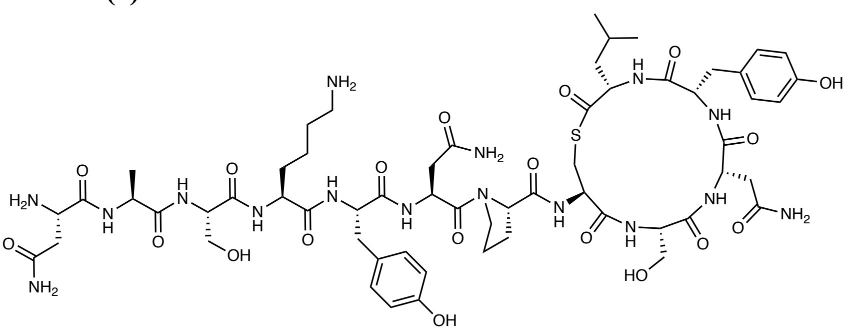

The peptide was synthesized according to general method A for on-resin cleavage-inducing cyclization (section 3). Purification by preparative RP-HPLC (5–40% B over 40 min, C18) afforded *S. epidermidis* AIP-II (**6**) as a fluffy white solid after lyophilization (8.4 mg, 26% based on resin loading). Purity 99% determined by UPLC ($\lambda$ = 215 nm). UPLC-MS (ESI) *m/z* calcd for $[M+2H]^{2+}$ $C_{59}H_{88}N_{16}O_{19}S^{2+}$: 678.31, found 678.61; $[M+H]^{+}$ $C_{59}H_{87}N_{16}O_{19}S^{+}$: 1355.61, found 1355.73. $^{1}H$ NMR (600 MHz, DMSO-$d_6$) $\delta$ = 0.75 (d, *J* = 6.5 Hz, 3H), 0.83 (d, *J* = 6.6 Hz, 3H), 1.18–1.27 (m, 5H), 1.29–1.37 (m, 1H), 1.41–1.53 (m, 3H), 1.54–1.64 (m, 2H), 1.64–1.71 (m, 1H), 1.80–1.89 (m, 3H), 1.93–2.01 (m, 1H), 2.37 (dd, *J* = 15.6, 6.3 Hz, 1H), 2.51–2.65 (m, 5H), 2.66–2.76 (m, 3H), 2.78–2.90 (m, 3H), 2.94 (dd, *J* = 13.7, 7.6 Hz, 1H), 3.13 (dd, *J* = 12.9, 4.1 Hz, 1H), 3.49 (dd, *J* = 11.2, 4.4 Hz, 1H), 3.52–3.66 (m, 5H), 4.05–4.15 (m, 2H), 4.14–4.23 (m, 3H), 4.27–4.35 (m, 3H), 4.36–4.46 (m, 3H), 4.77 (q, *J* = 7.2 Hz, 1H), 4.92 (s, 1H), 5.16 (s, 1H), 6.62 (d, *J* = 8.3 Hz, 2H), 6.65 (d, *J* = 8.3 Hz, 2H), 6.92 (s, 1H), 6.93–6.97 (m, 4H), 7.01 (s, 1H), 7.26 (s, 1H), 7.47 (s, 1H), 7.51 (s, 1H), 7.62–7.75 (m, 4H), 7.84 (d, *J* = 8.0 Hz, 1H), 7.90–7.96 (m, 2H), 8.04 (d, *J* = 8.9 Hz, 1H), 8.06–8.12 (m, 4H), 8.15 (d, *J* = 7.7 Hz, 1H), 8.19 (d, *J* = 8.0 Hz, 1H), 8.27 (d, *J* = 7.7 Hz, 1H), 8.56 (d, *J* = 8.4 Hz, 1H), 8.60 (d, *J* = 7.4 Hz, 1H), 9.20 (s, 2H).

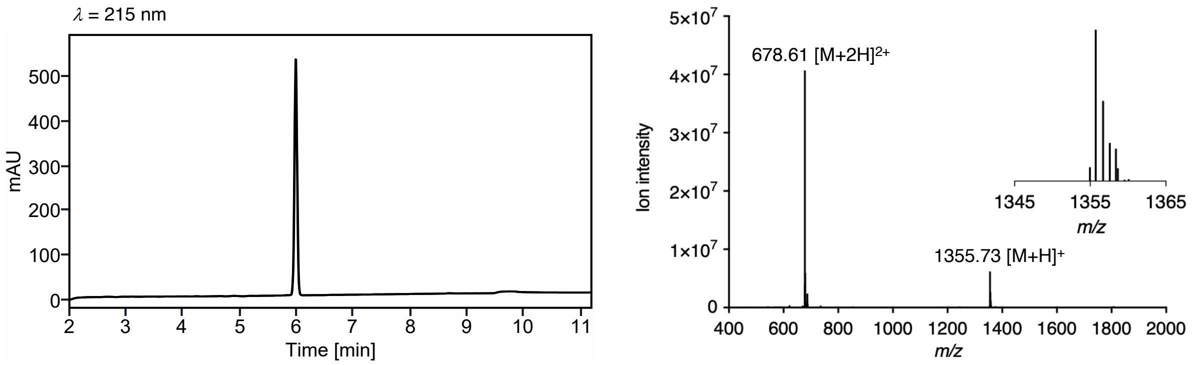

**Supplementary Figure 34. UPLC trace and MS spectrum of *S. epidermidis* AIP-II (6).**

***S. epidermidis* AIP-III (7)**

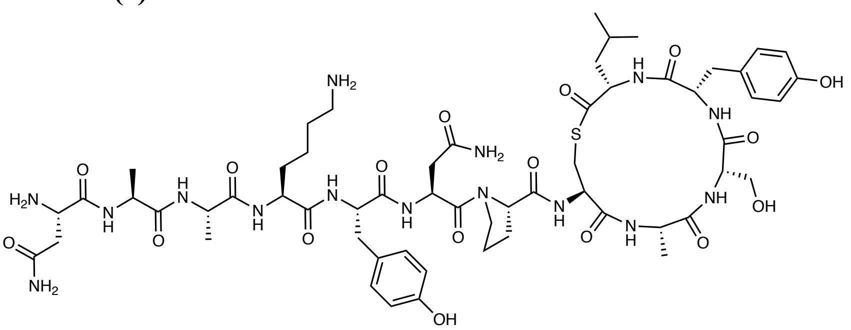

$C_{58}H_{85}N_{15}O_{17}S$
MW: 1296.47

The peptide was synthesized according to general method A for on-resin cleavage-inducing cyclization (section 3). Purification by preparative RP-HPLC (5–40% B over 40 min, C18) afforded *S. epidermidis* AIP-III (**7**) as a fluffy white solid after lyophilization (2.7 mg, 8.9% based on resin loading). Purity 95% determined by UPLC ($\lambda$ = 215 nm). UPLC-MS (ESI) *m/z* calcd for $[M+2H]^{2+}$ $C_{58}H_{87}N_{15}O_{17}S^{2+}$: 648.81, found 649.15; $[M+H]^{+}$ $C_{58}H_{86}N_{15}O_{17}S^{+}$: 1296.60, found 1296.72. $^{1}H$ NMR (600 MHz, DMSO-$d_6$) $\delta$ = 0.75 (d, *J* = 6.5 Hz, 3H), 0.82 (d, *J* = 6.6 Hz, 3H), 1.16–1.26 (m, 11H), 1.26–1.33 (m, 1H), 1.42–1.53 (m, 3H), 1.53–1.59 (m, 3H), 1.80–1.88 (m, 3H), 1.96–2.04 (m, 1H), 2.37 (dd, *J* = 15.5, 6.2 Hz, 1H), 2.54–2.71 (m, 4H), 2.71–2.78 (m, 2H), 2.80–2.89 (m, 3H), 2.90–2.95 (m, 1H), 3.11 (dd, *J* = 12.9, 3.8 Hz, 1H), 3.53–3.64 (m, 4H), 4.04–4.10 (m, 2H), 4.13–4.21 (m, 4H), 4.24 (t, *J* = 7.2 Hz, 1H), 4.25–4.30 (m, 1H), 4.30–4.36 (m, 2H), 4.44 (td, *J* = 7.9, 5.3 Hz, 1H), 4.77 (q, *J* = 7.3 Hz, 1H), 5.17 (s, 1H), 6.60 (d, *J* = 8.5 Hz, 2H), 6.64 (d, *J* = 8.5 Hz, 2H), 6.91–6.95 (m, 3H), 6.97 (d, *J* = 8.5 Hz, 2H), 7.26 (s, 1H), 7.49 (s, 1H), 7.66–7.75 (m, 5H), 7.77 (d, *J* = 8.7 Hz, 1H), 7.82 (d, *J* = 8.0 Hz, 1H), 7.96 (d, *J* = 7.6 Hz, 1H), 8.04 (d, *J* = 7.9 Hz, 1H), 8.08–8.11 (m, 3H), 8.13 (d, *J* = 7.3 Hz, 1H), 8.18 (d, *J* = 7.9 Hz, 1H), 8.38 (d, *J* = 7.7 Hz, 1H), 8.59 (d, *J* = 7.4 Hz, 1H), 8.68 (d, *J* = 8.3 Hz, 1H), 9.14–9.27 (m, 2H).

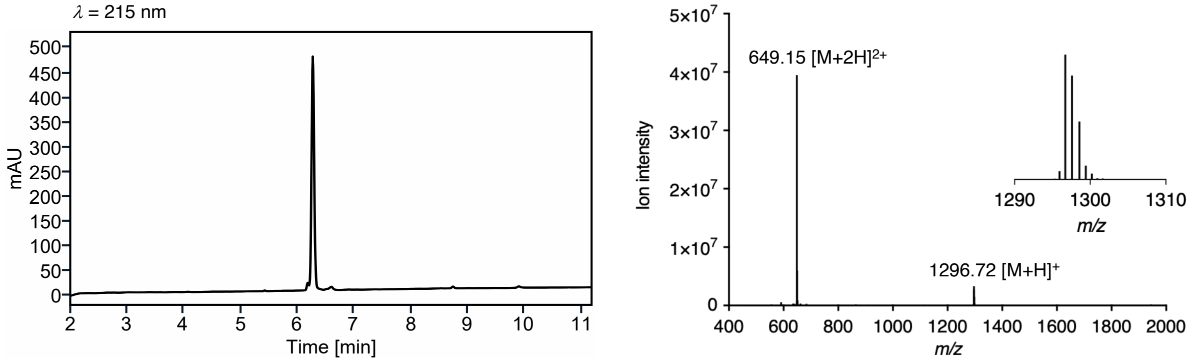

**Supplementary Figure 35. UPLC trace and MS spectrum of *S. epidermidis* AIP-III (7).**

***S. lugdunensis* AIP-I (8)**

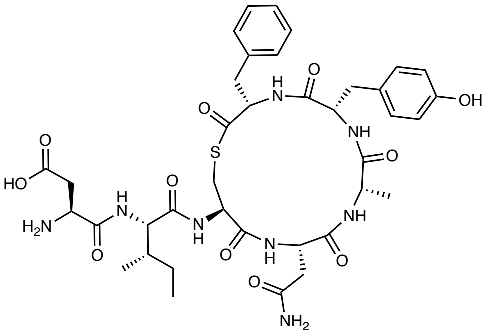

$C_{38}H_{50}N_8O_{11}S$
MW: 826.92

The peptide was synthesized as described in section 3. Purification by preparative RP-HPLC (5–40% B over 40 min, C18) afforded *S. lugdunensis* AIP-I (**8**) as a fluffy white solid after lyophilization (3.1 mg, 8% based on resin loading). Purity 97% determined by UPLC ($\lambda$ = 215 nm). UPLC-MS (ESI) *m/z* calcd for $[M+H]^+$ $C_{38}H_{51}N_8O_{11}S^+$: 827.34, found 827.40. $^1$H NMR (600 MHz, DMSO-$d_6$) $\delta$ = 0.79–0.83 (m, 6H), 1.03–1.11 (m, 1H), 1.17 (d, *J* = 6.9 Hz, 3H), 1.37–1.47 (m, 1H), 1.65–1.73 (m, 1H), 2.30 (dd, *J* = 15.7, 5.2 Hz, 1H), 2.50–2.59 (m, 2H), 2.64 (dd, *J* = 13.6, 7.0 Hz, 1H), 2.70 (dd, *J* = 17.3, 3.8 Hz, 1H), 2.76–2.86 (m, 2H), 2.92 (t, *J* = 12.0 Hz, 1H), 3.15 (dd, *J* = 12.7, 4.0 Hz, 1H), 3.27 (1H) signal partially overlapping with HDO peak, 3.98–4.04 (m, 1H), 4.06 (q, *J* = 7.6 Hz, 1H), 4.12–4.20 (m, 1H), 4.21 (t, *J* = 7.2 Hz, 1H), 4.31 (td, *J* = 9.2, 4.1 Hz, 1H), 4.43 (dt, *J* = 8.4, 5.8 Hz, 1H), 4.48 (ddd, *J* = 11.8, 8.1, 4.0 Hz, 1H), 6.64 (d, *J* = 8.2 Hz, 2H), 6.87 (d, *J* = 8.2 Hz, 2H), 6.91 (s, 1H), 7.03 (d, *J* = 7.4 Hz, 2H), 7.15–7.19 (m, 1H), 7.19–7.25 (m, 2H), 7.40 (s, 1H), 7.86 (d, *J* = 8.7 Hz, 1H), 8.13 (d, *J* = 8.5 Hz, 1H), 8.25 (d, *J* = 7.7 Hz, 1H), 8.45 (d, *J* = 8.5 Hz, 1H), 8.59 (d, *J* = 8.1 Hz, 1H), 8.80 (d, *J* = 8.4 Hz, 1H), 9.23 (s, 1H).

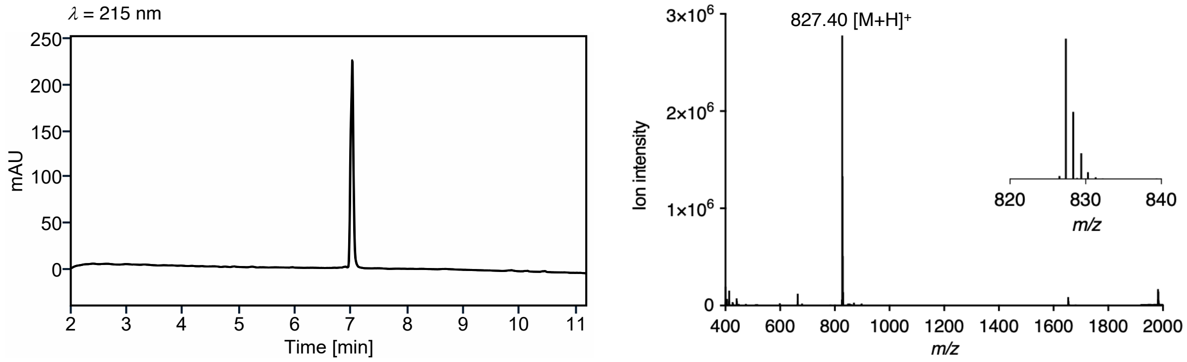

**Supplementary Figure 36. UPLC trace and MS spectrum of *S. lugdunensis* AIP-I (8).**

##### *S. hominis* AIP-II (11)

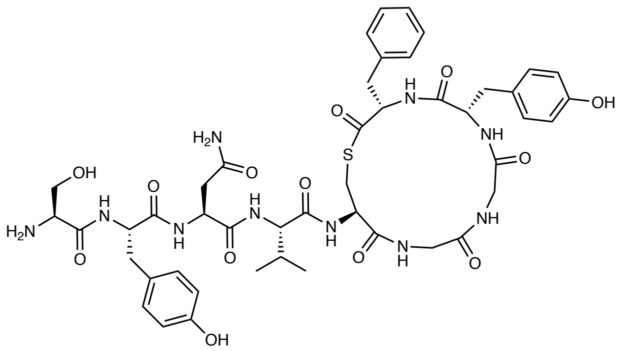

$C_{46}H_{58}N_{10}O_{13}S$
MW: 991.09

The peptide was synthesized according to general method A for on-resin cleavage-inducing cyclization (section 3). Purification by preparative RP-HPLC (5–40% B over 40 min, C18) afforded *S. hominis* AIP-II (**11**) as a fluffy white solid after lyophilization (2.8 mg, 13% based on resin loading). Purity 97% determined by UPLC ($\lambda$ = 215 nm). UPLC-MS (ESI) *m/z* calcd for $[M+H]^+$ $C_{46}H_{59}N_{10}O_{13}S^+$: 991.40, found 991.57. $^1$H NMR (600 MHz, DMSO-$d_6$) $\delta$ = 0.81 (d, $J$ = 6.8 Hz, 3H), 0.83 (d, $J$ = 6.8 Hz, 3H), 1.95–2.07 (m, 1H), 2.32 (dd, $J$ = 14.3, 10.6 Hz, 1H), 2.42 (dd, $J$ = 15.5, 7.4 Hz, 1H), 2.57 (dd, $J$ = 15.6, 6.2 Hz, 1H), 2.62–2.70 (m, 2H), 2.76 (dd, $J$ = 13.0, 10.5 Hz, 1H), 2.88–2.98 (m, 2H), 3.22 (dd, $J$ = 13.0, 4.8 Hz, 1H), 3.34 (2H) signal partially overlapping with HDO peak, 3.43–3.49 (m, 1H), 3.60–3.67 (m, 1H), 3.73–3.82 (m, 3H), 4.12–4.19 (m, 2H), 4.22 (dd, $J$ = 8.8, 5.9 Hz, 1H), 4.41 (ddd, $J$ = 10.4, 7.7, 5.0 Hz, 1H), 4.53 (td, $J$ = 8.9, 3.9 Hz, 1H), 4.58–4.65 (m, 2H), 5.43–5.49 (m, 1H), 6.60–6.65 (m, 4H), 6.89 (d, $J$ = 8.5 Hz, 2H), 6.96 (s, 1H), 7.05 (d, $J$ = 8.5 Hz, 2H), 7.20–7.26 (m, 3H), 7.27–7.32 (m, 2H), 7.42 (s, 1H), 7.76 (d, $J$ = 8.9 Hz, 1H), 7.91–8.05 (m, 3H), 8.26–8.29 (m, 2H), 8.30–8.33 (m, 1H), 8.35 (dd, $J$ = 8.2, 4.4 Hz, 1H), 8.39–8.44 (m, 2H), 8.54 (d, $J$ = 8.3 Hz, 1H), 9.18 (s, 1H), 9.19 (s, 1H).

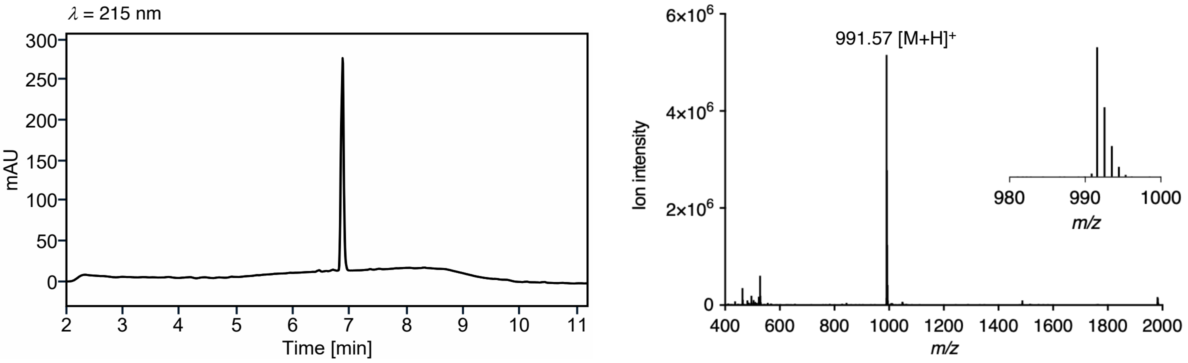

**Supplementary Figure 37. UPLC trace and MS spectrum of *S. hominis* AIP-II (11).**

***S. caprae* AIP-I (12)**

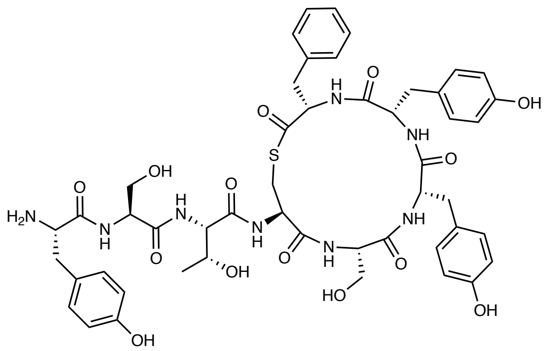

The peptide was synthesized according to general method A for on-resin cleavage-inducing cyclization (section 3). Purification by preparative RP-HPLC (5–40% B over 40 min, C18) afforded *S. caprae* AIP-I (**12**) as a fluffy white solid after lyophilization (4.5 mg, 20% based on resin loading). Purity 98% determined by UPLC ($\lambda$ = 215 nm). UPLC-MS (ESI) *m/z* calcd for $[M+H]^+$ $C_{49}H_{59}N_8O_{14}S^+$: 1015.39, found 1015.53. $^1$H NMR (600 MHz, DMSO-$d_6$) $\delta$ = 1.05 (d, $J$ = 6.3 Hz, 3H), 2.62–2.72 (m, 2H), 2.73–2.86 (m, 4H), 2.90 (dd, $J$ = 12.7, 11.0 Hz, 1H), 3.03 (dd, $J$ = 14.3, 4.7 Hz, 1H), 3.20–3.28 (m, 2H), 3.35–3.39 (m, 1H), 3.44–3.50 (m, 1H), 3.54–3.61 (m, 1H), 3.65–3.72 (m, 1H), 4.01 (dd, $J$ = 8.3, 4.8 Hz, 1H), 4.06–4.14 (m, 4H), 4.26 (dd, $J$ = 8.5, 3.4 Hz, 1H), 4.36 (dt, $J$ = 9.2, 7.0 Hz, 1H), 4.45 (ddd, $J$ = 11.7, 8.1, 4.0 Hz, 1H), 4.55 (dt, $J$ = 7.7, 6.2 Hz, 1H), 4.92 (d, $J$ = 5.1 Hz, 1H), 5.03 (t, $J$ = 5.3 Hz, 1H), 5.25 (t, $J$ = 5.2 Hz, 1H), 6.60–6.66 (m, 4H), 6.69 (d, $J$ = 8.5 Hz, 1H), 6.78 (d, $J$ = 8.5 Hz, 2H), 6.89–6.96 (m, 4H), 7.07 (d, $J$ = 8.5 Hz, 2H), 7.14–7.22 (m, 3H), 7.85 (d, $J$ = 9.1 Hz, 1H), 7.86–7.99 (m, 3H), 7.97 (d, $J$ = 8.5 Hz, 1H), 8.02 (d, $J$ = 8.1 Hz, 1H), 8.18–8.23 (m, 2H), 8.72 (d, $J$ = 7.7 Hz, 1H), 8.94 (d, $J$ = 7.9 Hz, 1H), 9.22 (s, 1H), 9.24 (s, 1H), 9.34 (s, 1H).

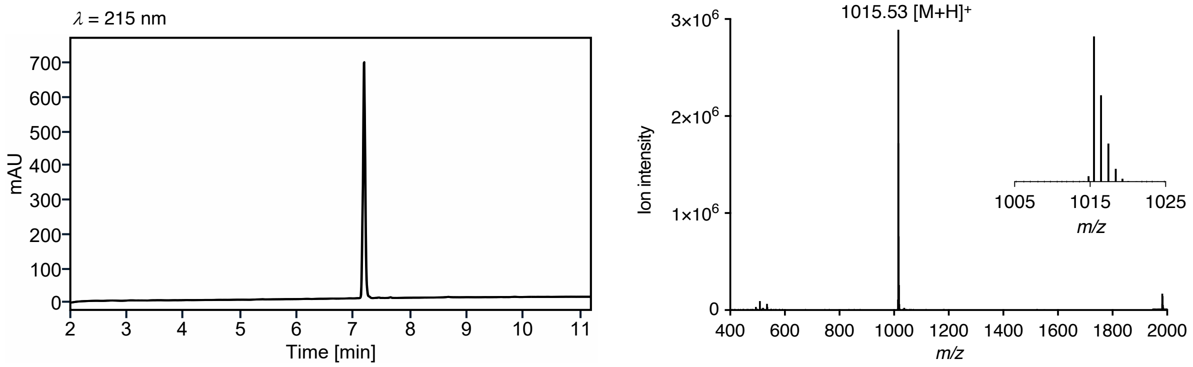

**Supplementary Figure 38. UPLC trace and MS spectrum of *S. caprae* AIP-I (12).**

***S. pasteuri* AIP-I (15)**

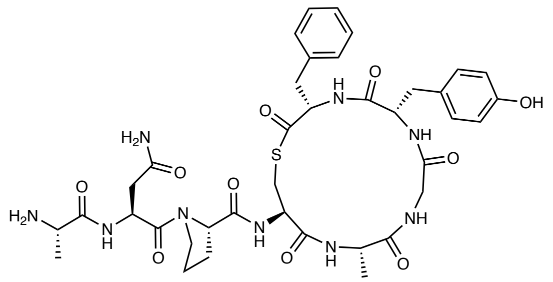

The peptide was synthesized according to general method A for on-resin cleavage-inducing cyclization (section 3). Purification by preparative RP-HPLC (5–40% B over 40 min, C18) afforded *S. pasteuri* AIP-I (**15**) as a fluffy white solid after lyophilisation (8.9 mg, 47% based on resin loading). Purity 99% determined by UPLC ($\lambda$ = 215 nm). UPLC-MS (ESI) *m/z* calcd for $[M+H]^+$ $C_{38}H_{50}N_9O_{10}S^+$: 824.34, found 824.26. Major conformer (conformers were observed in a ration of 1:0.15): $^1$H NMR (600 MHz, DMSO-$d_6$) $\delta$ = 1.19 (d, $J$ = 7.1 Hz, 3H), 1.31 (d, $J$ = 7.0 Hz, 3H), 1.79–1.94 (m, 3H), 1.95–2.03 (m, 1H), 2.31 (dd, $J$ = 14.3, 10.8 Hz, 1H), 2.42 (dd, $J$ = 15.7, 7.9 Hz, 1H), 2.59 (dd, $J$ = 15.7, 5.5 Hz, 1H), 2.63–2.72 (m, 2H), 2.97 (dd, $J$ = 14.2, 11.2 Hz, 1H), 3.18 (dd, $J$ = 13.0, 5.0 Hz, 1H), 3.30–3.39 (m, 2H), 3.62–3.67 (m, 2H), 3.78–3.88 (m, 2H), 4.14 (ddd, $J$ = 11.5, 7.9, 4.0 Hz, 1H), 4.34–4.42 (m, 2H), 4.43–4.49 (m, 1H), 4.60 (ddd, $J$ = 11.2, 9.3, 3.8 Hz, 1H), 4.81–4.88 (m, 1H), 6.62 (d, $J$ = 8.4 Hz, 2H), 6.90 (d, $J$ = 8.4 Hz, 2H), 6.99 (s, 1H), 7.20–7.24 (m, 1H), 7.25–7.33 (m, 4H), 7.48 (s, 1H), 8.03–8.10 (m, 4H), 8.21 (d, $J$ = 9.4 Hz, 1H), 8.25 (d, $J$ = 8.7 Hz, 1H), 8.48 (d, $J$ = 8.0 Hz, 1H), 8.53 (t, $J$ = 5.5 Hz, 1H), 8.70 (d, $J$ = 7.3 Hz, 1H), 9.20 (br s, 1H).

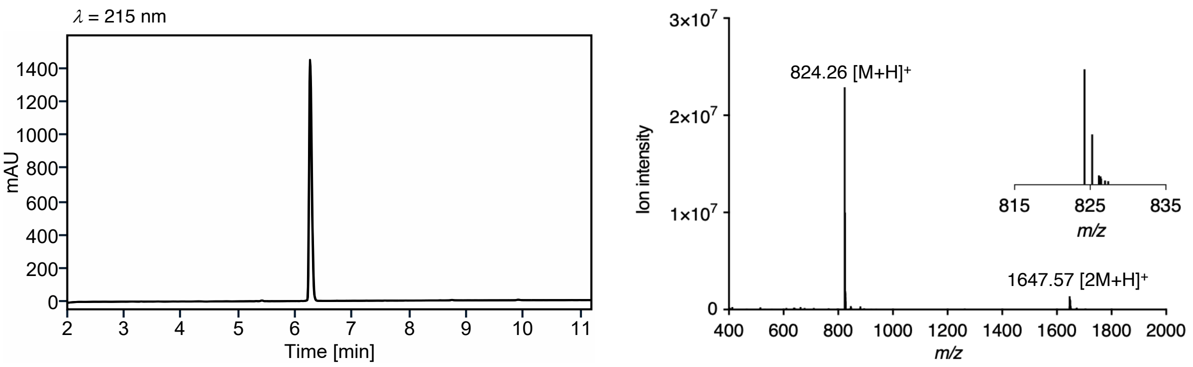

**Supplementary Figure 39. UPLC trace and MS spectrum of *S. pasteuri* AIP-I (15).**

***S. succinus* AIP-I (16)**

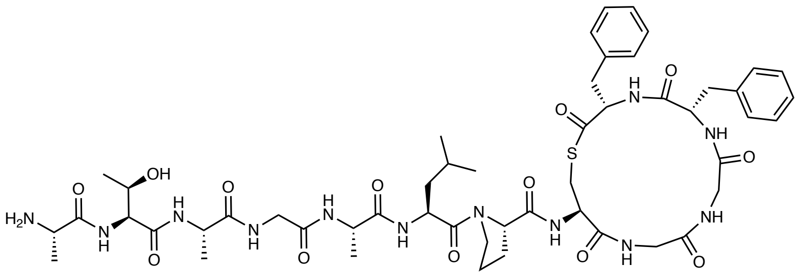

$C_{51}H_{72}N_{12}O_{13}S$
MW: 1093.27

The peptide was synthesized according to general method A for on-resin cleavage-inducing cyclization (section 3). Purification by preparative RP-HPLC (10–50% B over 40 min, C18) afforded *S. succinus* AIP-I (**16**) as a fluffy white solid after lyophilization (10.3 mg, 47% based on resin loading). Purity 98% determined by UPLC ($\lambda$ = 215 nm). UPLC-MS (ESI) *m/z* calcd for $[M+2H]^{2+}$ $C_{51}H_{74}N_{12}O_{13}S^{2+}$: 547.26, found 547.39; $[M+H]^{+}$ $C_{51}H_{73}N_{12}O_{13}S^{+}$: 1093.51, found 1093.48. Major conformer (conformers were observed in a ration of 1:0.07): $^{1}$H NMR (600 MHz, DMSO-$d_6$) $\delta$ = 0.87–0.92 (m, 6H), 1.08 (d, $J$ = 6.3 Hz, 3H), 1.18 (d, $J$ = 7.0 Hz, 3H), 1.24 (d, $J$ = 7.0 Hz, 3H), 1.35 (d, $J$ = 6.9 Hz, 3H), 1.40–1.52 (m, 2H), 1.60–1.69 (m, 1H), 1.71–1.77 (m, 1H), 1.82–1.88 (m, 1H), 1.88–1.95 (m, 1H) 1.98–2.07 (m, 1H), 2.44 (dd, $J$ = 14.3, 10.7 Hz, 1H), 2.69 (dd, $J$ = 13.0, 10.2 Hz, 1H), 2.78 (dd, $J$ = 14.3, 4.1 Hz, 1H), 2.93 (dd, $J$ = 14.1, 11.0 Hz, 1H), 3.23 (dd, $J$ = 13.0, 4.8 Hz, 1H), 3.30–3.37 (m, 2H), 3.42–3.51 (m, 2H), 3.62–3.75 (m, 3H), 3.79 (dd, $J$ = 14.2, 4.4 Hz, 1H), 3.96–4.06 (m, 2H), 4.17 (dd, $J$ = 15.6, 8.4 Hz, 1H), 4.22–4.33 (m, 4H), 4.34–4.40 (m, 2H), 4.50 (ddd, $J$ = 9.6, 8.0, 4.8 Hz, 1H), 4.64 (ddd, $J$ = 11.0, 9.2, 4.0 Hz, 1H), 7.08–7.12 (m, 2H), 7.16–7.20 (m, 1H), 7.20–7.28 (m, 5H), 7.29–7.34 (m, 2H), 7.85 (d, $J$ = 7.5 Hz, 1H), 8.01 (d, $J$ = 7.0 Hz, 1H), 8.03–8.07 (m, 4H), 8.17 (t, $J$ = 5.8 Hz, 1H), 8.20 (d, $J$ = 7.7 Hz, 1H), 8.31 (d, $J$ = 9.3 Hz, 1H), 8.33–8.39 (m, 2H), 8.41 (d, $J$ = 8.4 Hz, 1H), 8.51 (d, $J$ = 8.0 Hz, 1H).

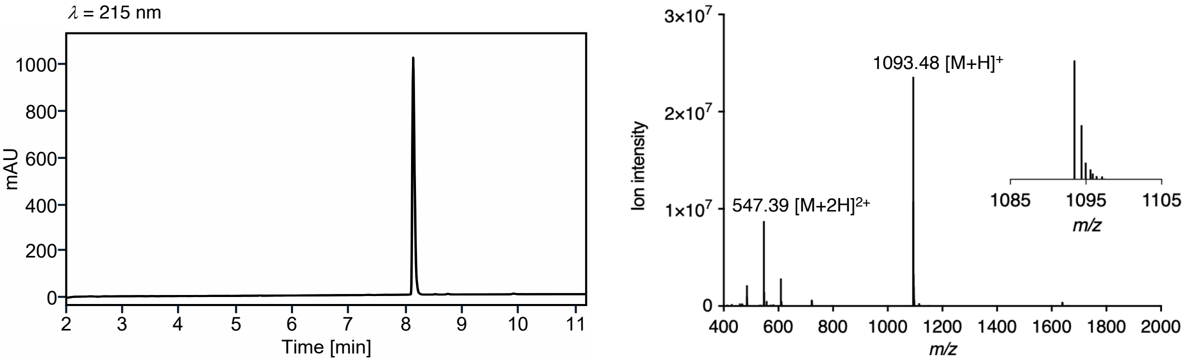

**Supplementary Figure S40. UPLC trace and MS spectrum of *S. succinus* AIP-I (16).**

***S. cohnii* AIP-I (17)**

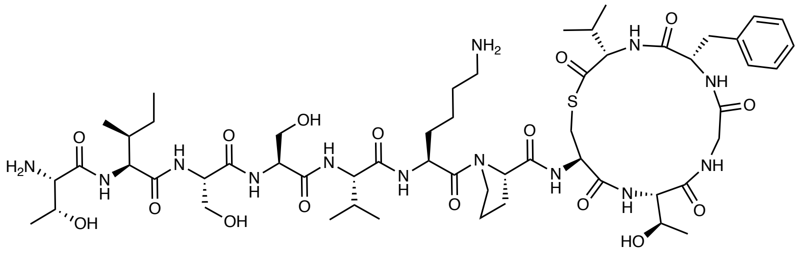

$C_{55}H_{89}N_{13}O_{16}S$
MW: 1220.45

The peptide was synthesized according to general method A for on-resin cleavage-inducing cyclization (section 3). Purification by preparative RP-HPLC (5–40% B over 40 min, C18) afforded *S. cohnii* AIP-I (**17**) as a fluffy white solid after lyophilization (5.3 mg, 18% based on resin loading). Purity 95% determined by UPLC ($\lambda$ = 215 nm). UPLC-MS (ESI) *m/z* calcd for $[M+2H]^{2+}$ $C_{55}H_{91}N_{13}O_{16}S^{2+}$: 610.82, found 610.93; $[M+H]^{+}$ $C_{55}H_{90}N_{13}O_{16}S^{+}$: 1220.63, found 1220.69. $^{1}$H NMR (600 MHz, DMSO-$d_6$) $\delta$ = 0.80 (d, $J$ = 6.7 Hz, 3H), 0.81–0.85 (m, 6H), 0.86 (d, $J$ = 6.8 Hz, 3H), 0.90 (d, $J$ = 6.8 Hz, 3H), 0.95 (d, $J$ = 6.9 Hz, 3H), 0.99 (d, $J$ = 6.2 Hz, 3H), 1.05–1.12 (m, 1H), 1.14 (d, $J$ = 6.3 Hz, 3H), 1.34 –1.40 (m 2H), 1.44–1.59 (m, 4H), 1.62–1.69 (m, 1H), 1.69–1.77 (m, 2H), 1.80–1.92 (m, 2H), 1.93–2.04 (m, 2H), 2.37–2.45 (m, 1H), 2.63 (dd, $J$ = 13.1, 10.8 Hz, 1H), 2.75–2.81 (m, 2H), 2.85 (dd, $J$ = 14.2, 11.4 Hz, 1H), 3.21 (dd, $J$ = 13.1, 4.8 Hz, 1H), 3.25 (dd, $J$ = 14.2, 3.5 Hz, 1H), 3.41–3.46 (m, 1H), 3.50–3.61 (m, 4H), 3.61–3.71 (m, 4H), 3.76–3.83 (m, 1H), 3.86–3.91 (m, 1H), 4.19 (dd, $J$ = 8.8, 6.2 Hz, 1H), 4.24 (dd, $J$ = 9.1, 4.4 Hz, 1H), 4.32 (dd, $J$ = 8.6, 7.1 Hz, 1H), 4.34–4.46 (m, 6H), 4.56 (dd, $J$ = 9.9, 4.6 Hz, 1H), 4.76-4.80 (m, 1H), 4.95–4.99 (m, 1H), 5.05–5.09 (m, 1H), 5.52–5.58 (m, 1H), 7.19–7.24 (m, 1H), 7.24–7.34 (m, 5H), 7.65–7.75 (m, 4H), 7.89–7.96 (m, 2H), 8.03–8.10 (m, 3H), 8.11–8.16 (m, 2H), 8.19 (d, $J$ = 7.7 Hz, 1H), 8.44 (d, $J$ = 8.6 Hz, 1H), 8.53 (dd, $J$ = 6.8, 4.6 Hz, 1H), 8.77 (d, $J$ = 8.4 Hz, 1H).

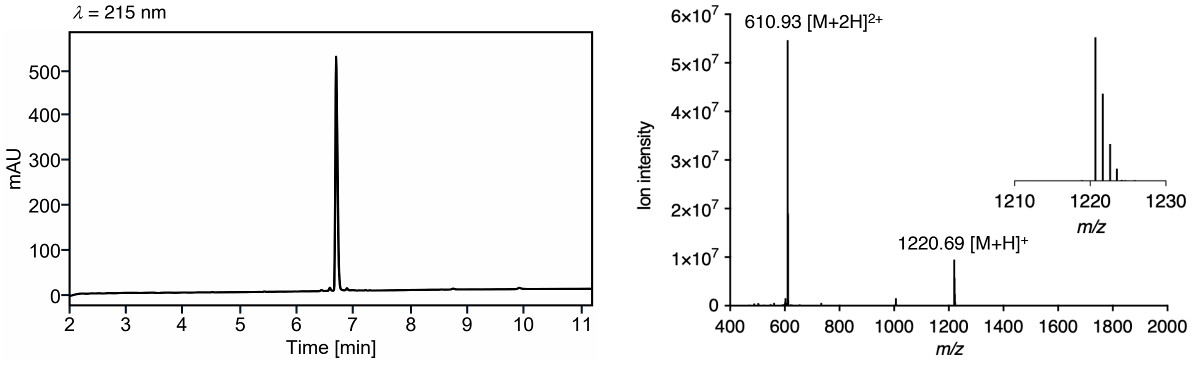

**Supplementary Figure 41. UPLC trace and MS spectrum of *S. cohnii* AIP-I (17).**

##### *S. intermedius* AIP-I (22)

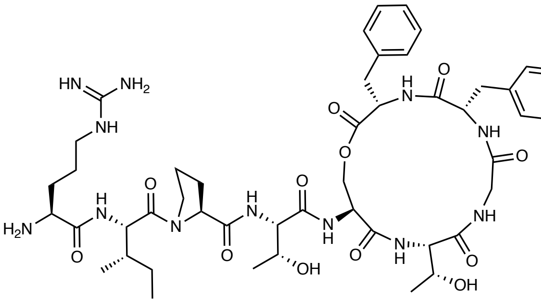

The peptide was synthesized as described in section 3. Purification by preparative RP-HPLC (5–40% B over 40 min, C18) afforded *S. intermedius* AIP-I (**22**) as a fluffy white solid after lyophilization (6.7 mg, 14% based on resin loading). Purity 96% determined by UPLC ($\lambda$ = 215 nm). UPLC-MS (ESI) *m/z* calcd for $[M+2H]^{2+}$ $C_{48}H_{72}N_{12}O_{12}^{2+}$: 504.27, found 504.53; $[M+H]^{+}$ $C_{48}H_{71}N_{12}O_{12}^{+}$: 1007.53, found 1007.68. Major conformer (conformers were observed in a ration of 1:0.12): $^{1}$H NMR (600 MHz, DMSO-$d_6$) $\delta$ = 0.81–0.86 (t, $J$ = 7.4 Hz, 3H), 0.92 (d, $J$ = 6.7 Hz, 3H), 1.03 (d, $J$ = 6.3 Hz, 3H), 1.07 (d, $J$ = 6.3 Hz, 3H), 1.08–1.13 (m, 1H), 1.41–1.51 (m, 2H), 1.51–1.59 (m, 1H), 1.60–1.71 (m, 1H), 1.72–1.88 (m, 3H), 1.88–1.96 (m, 1H), 1.96–2.05 (m, 1H), 2.82 (dd, $J$ = 14.1, 11.5 Hz, 1H), 3.03 (dd, $J$ = 13.8, 9.3 Hz, 1H), 3.06–3.16 (m, 3H), 3.26–3.32 (m, 1H), 3.60 (dt, $J$ = 9.7, 6.6 Hz, 1H), 3.70–3.79 (m, 2H), 3.82–3.92 (m, 3H), 4.01–4.06 (m, 1H), 4.12–4.20 (m, 2H), 4.19–4.27 (m, 2H), 4.38 (t, $J$ = 8.3 Hz, 1H), 4.43 (dd, $J$ = 8.2, 4.6 Hz, 1H), 4.62 (td, $J$ = 9.2, 6.2 Hz, 1H), 4.75–4.83 (m, 1H), 4.90–5.07 (m, 2H), 6.85–7.48 (4H) broad signal of guanidium group, 7.16–7.36 (m, 10H), 7.64 (d, $J$ = 7.9 Hz, 1H), 7.70 (t, $J$ = 6.0 Hz, 1H), 7.86 (d, $J$ = 8.5 Hz, 1H), 7.93 (d, $J$ = 9.1 Hz, 1H), 8.12–8.21 (m, 3H), 8.31 (d, $J$ = 8.0 Hz, 1H), 8.39 (d, $J$ = 7.2 Hz, 1H), 8.59 (d, $J$ = 8.0 Hz, 1H), 9.02 (t, $J$ = 5.6 Hz, 1H).

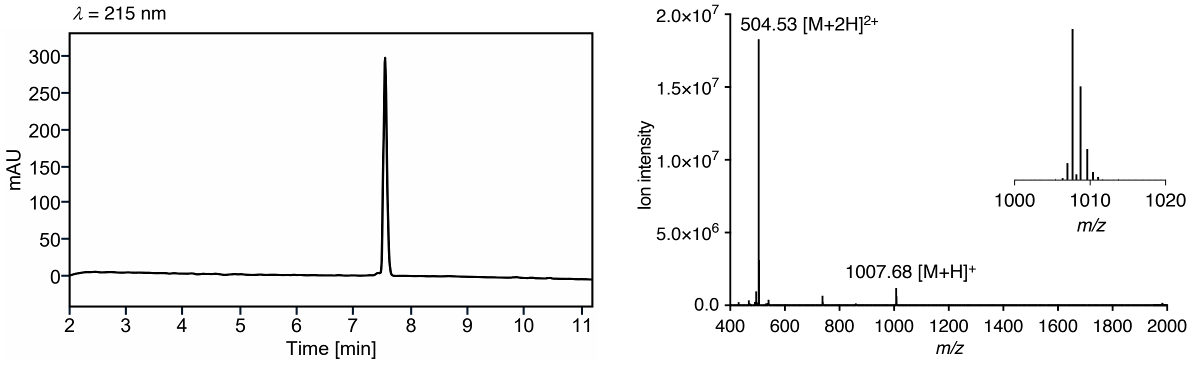

**Supplementary Figure 42. UPLC trace and MS spectrum of *S. intermedius* AIP-I (22).**

***S. simulans* AIP-II (24)**

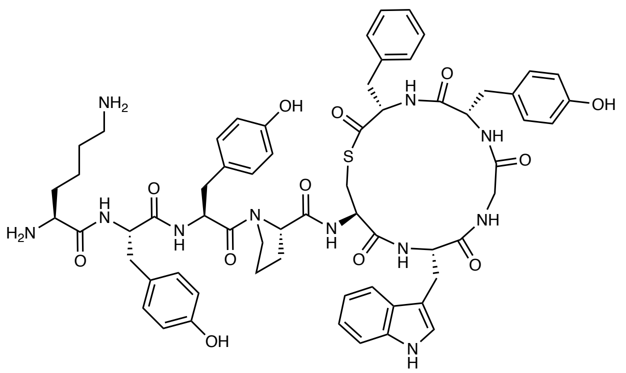

The peptide was synthesized according to general method A for on-resin cleavage-inducing cyclization (section 3). Purification by preparative RP-HPLC (10–50% B over 40 min, C18) afforded *S. simulans* AIP-II (**24**) as a fluffy white solid after lyophilization (8.2 mg, 29% based on resin loading). Purity 98% determined by UPLC ($\lambda$ = 215 nm). UPLC-MS (ESI) *m/z* calcd for $[M+2H]^{2+}$ $C_{63}H_{75}N_{11}O_{12}S^{2+}$: 604.77, found 605.07; $[M+H]^{+}$ $C_{63}H_{74}N_{11}O_{12}S^{+}$: 1208.52, found 1208.70. Major conformer (conformers were observed in a ration of 1:0.21): $^{1}$H NMR (600 MHz, DMSO-$d_6$) $\delta$ = 1.24–1.34 (m, 2H), 1.47–1.55 (m, 2H), 1.63–1.74 (m, 3H), 1.75–1.82 (m, 1H), 1.83–1.95 (m, 2H), 2.32 (dd, *J* = 14.2, 10.9 Hz, 1H), 2.57–2.78 (m, 6H), 2.86–2.99 (m, 3H), 3.00–3.09 (m, 2H), 3.27 (dd, *J* = 12.9, 5.0 Hz, 1H), 3.31–3.41 (3H) signals overlapping with HDO peak, 3.48–3.54 (m, 1H), 3.69–3.76 (m, 1H), 3.84 (dd, *J* = 14.0, 4.7 Hz, 1H), 4.11–4.19 (m, 1H), 4.33–4.38 (m, 1H), 4.44–4.53 (m, 2H), 4.52–4.60 (m, 1H), 4.59–4.66 (m, 1H), 4.65–4.72 (m, 1H), 6.59–6.69 (m, 6H), 6.89–6.93 (m, 2H), 6.94–6.99 (m, 1H), 7.01 (d, *J* = 8.5 Hz, 2H), 7.03–7.07 (m, 1H), 7.08–7.13 (m, 2H), 7.15 (d, *J* = 2.4 Hz, 1H), 7.20–7.26 (m, 1H), 7.26–7.37 (m, 5H), 7.59 (d, *J* = 7.9 Hz, 1H), 7.65–7.76 (m, 3H), 7.99–8.09 (m, 5H), 8.19 (d, *J* = 9.3 Hz, 1H), 8.34–8.40 (m, 2H), 8.46 (d, *J* = 8.2 Hz, 1H), 8.51 (d, *J* = 8.0 Hz, 1H), 8.73 (t, *J* = 5.6 Hz, 1H), 9.16–9.21 (m, 3H), 10.80 (d, *J* = 2.4 Hz, 1H).

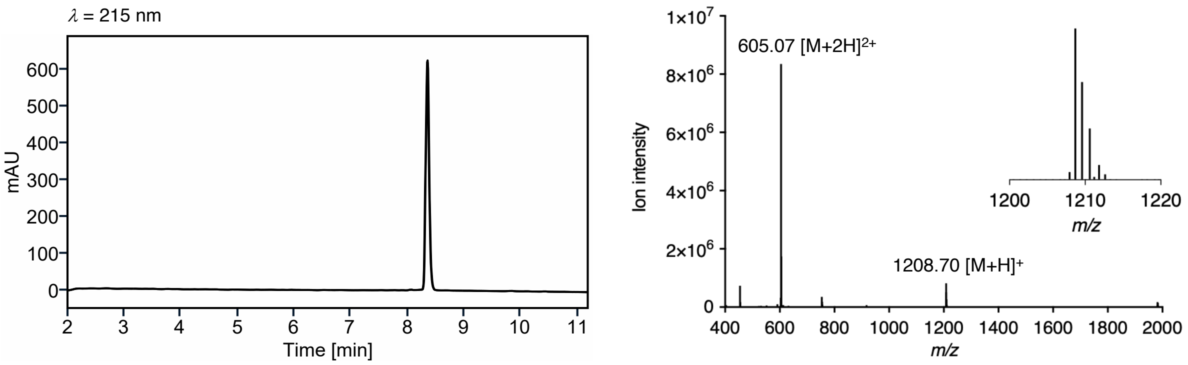

**Supplementary Figure 43. UPLC trace and MS spectrum of *S. simulans* AIP-II (24).**

***S. simulans* AIP-III (25)**

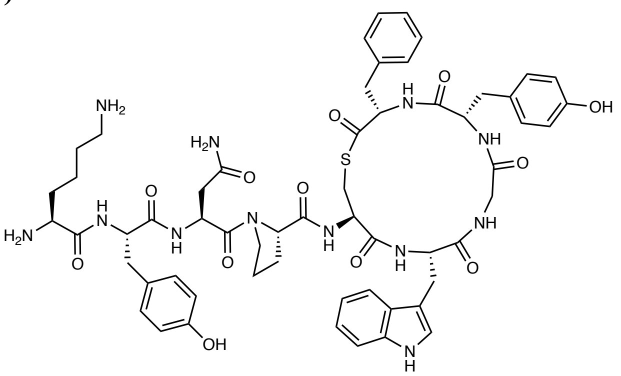

The peptide was synthesized according to general method A for on-resin cleavage-inducing cyclization (section 3). Purification by preparative RP-HPLC (10–50% B over 40 min, C18) afforded *S. simulans* AIP-III (**25**) as a fluffy white solid after lyophilization (7.3 mg, 29% based on resin loading). Purity 98% determined by UPLC ($\lambda$ = 215 nm). UPLC-MS (ESI) *m/z* calcd for $[M+2H]^{2+}$ $C_{58}H_{72}N_{12}O_{12}S^{2+}$: 580.25, found 580.46; $[M+H]^{+}$ $C_{58}H_{71}N_{12}O_{12}S^{+}$: 1159.50, found 1159.68. Major conformer (conformers were observed in a ration of 1:0.10): $^{1}$H NMR (600 MHz, DMSO-$d_6$) $\delta$ = 1.27–1.36 (m, 2H), 1.48–1.57 (m, 2H), 1.66–1.74 (m, 2H), 1.73–1.80 (m, 1H), 1.78–1.88 (m, 2H), 1.86–1.95 (m, 1H), 2.31 (dd, *J* = 14.2, 10.7 Hz, 1H), 2.38 (dd, *J* = 15.6, 7.1 Hz, 1H), 2.58–2.64 (m, 1H), 2.64–2.71 (m, 3H), 2.72–2.81 (m, 2H), 2.89 (dd, *J* = 14.1, 4.7 Hz, 1H), 2.94–3.08 (m, 3H), 3.16 (dd, *J* = 13.0, 4.9 Hz, 1H), 3.31–3.41 (2H) signals overlapping with HDO peak, 3.50–3.62 (m, 2H), 3.72–3.81 (m, 2H), 4.11–4.17 (m, 1H), 4.32–4.37 (m, 1H), 4.36–4.43 (m, 1H), 4.48–4.55 (m, 1H), 4.58–4.68 (m, 2H), 4.82 (q, *J* = 7.1 Hz, 1H), 6.60–6.67 (m, 4H), 6.90 (d, *J* = 8.4 Hz, 2H), 6.93–6.99 (m, 2H), 7.02 (d, *J* = 8.5 Hz, 2H), 7.03–7.07 (m, 1H), 7.16 (d, *J* = 2.3 Hz, 1H), 7.20–7.26 (m, 1H), 7.26–7.34 (m, 5H), 7.47 (s, 1H), 7.56 (d, *J* = 7.9 Hz, 1H), 7.68–7.76 (m, 3H), 8.03 (d, *J* = 8.0 Hz, 1H), 8.06–8.09 (m, 3H), 8.18–8.26 (m, 2H), 8.48–8.54 (m, 3H), 8.64 (t, *J* = 5.6 Hz, 1H), 9.13–9.35 (m, 2H), 10.81 (d, *J* = 2.5 Hz, 1H).

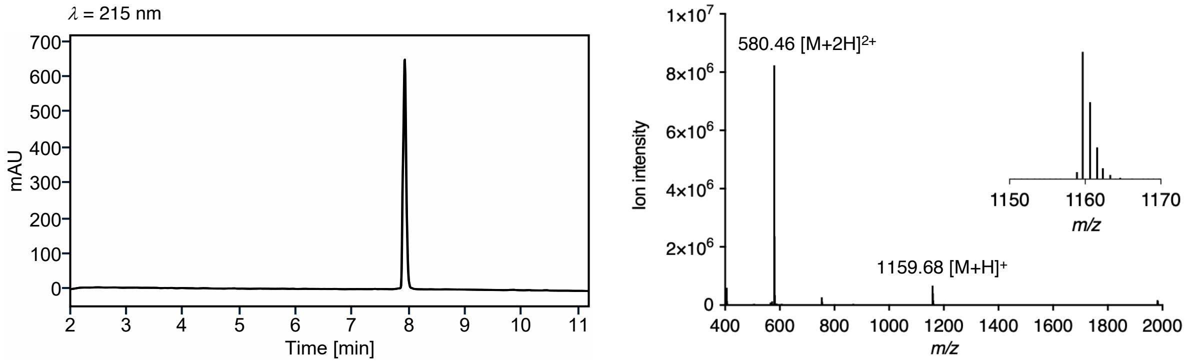

**Supplementary Figure 44. UPLC trace and MS spectrum of *S. simulans* AIP-III (25).**

#### Copies of NMR spectra

##### *S. epidermidis* AIP-I (5)

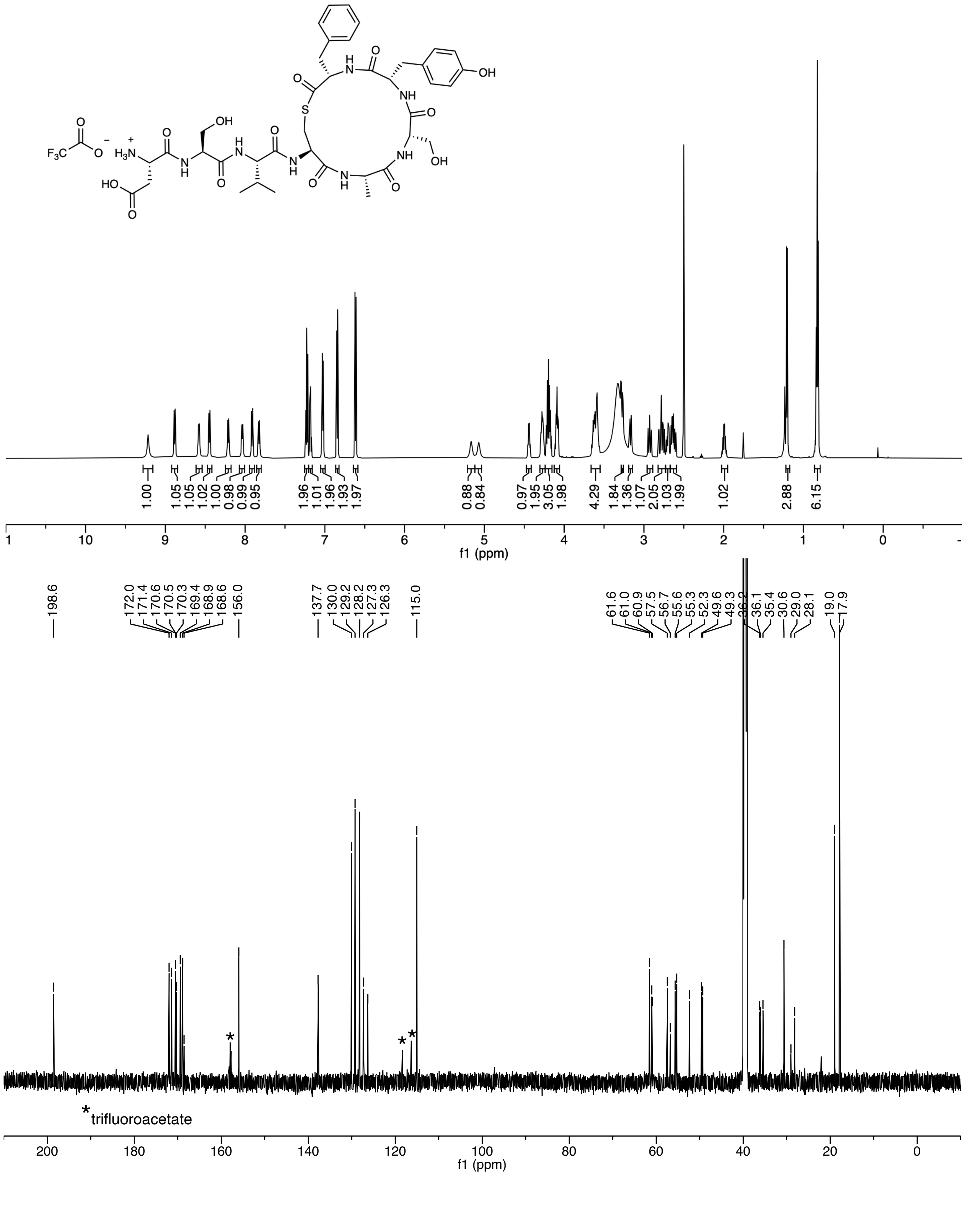

***S. epidermidis* AIP-II (6)**

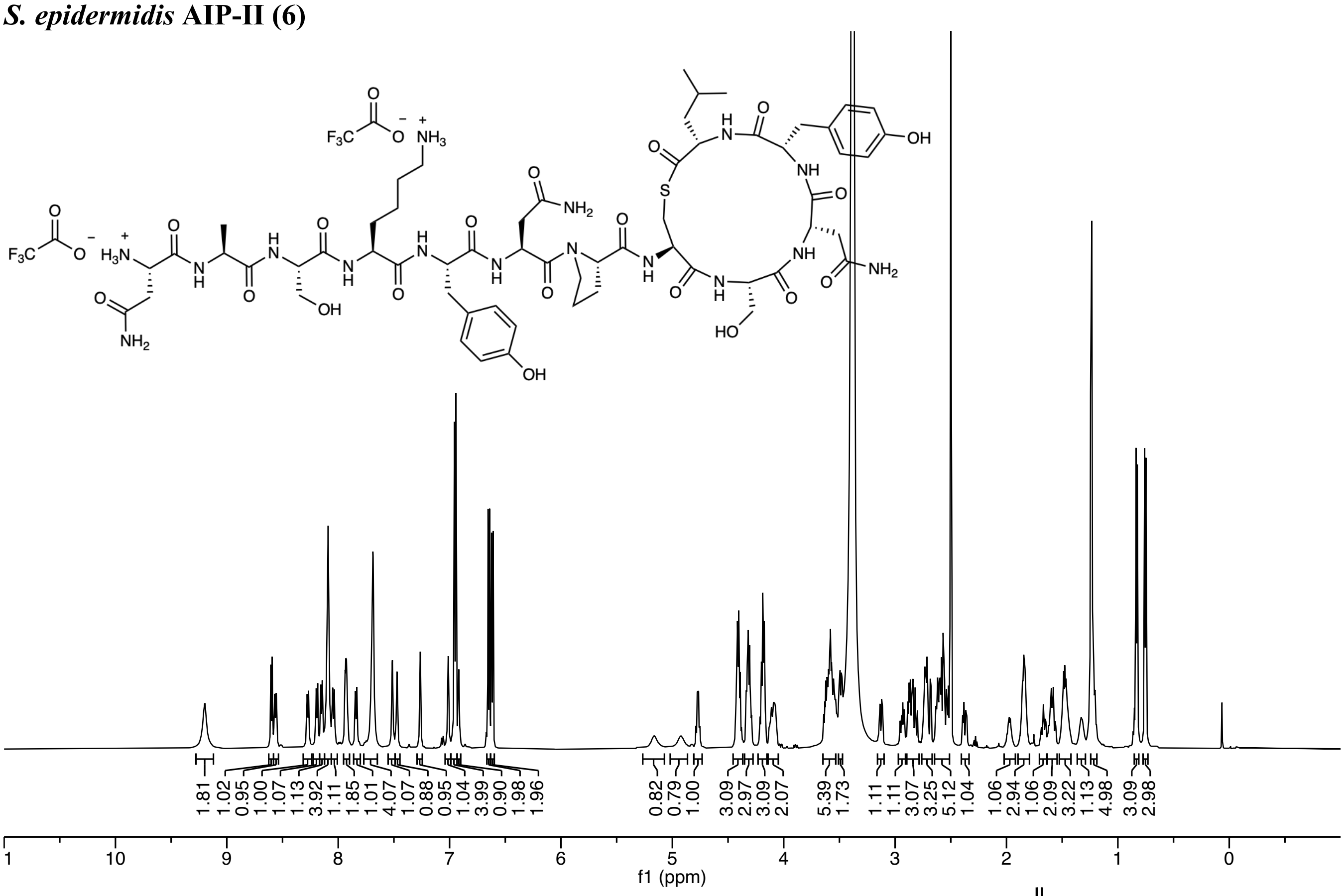

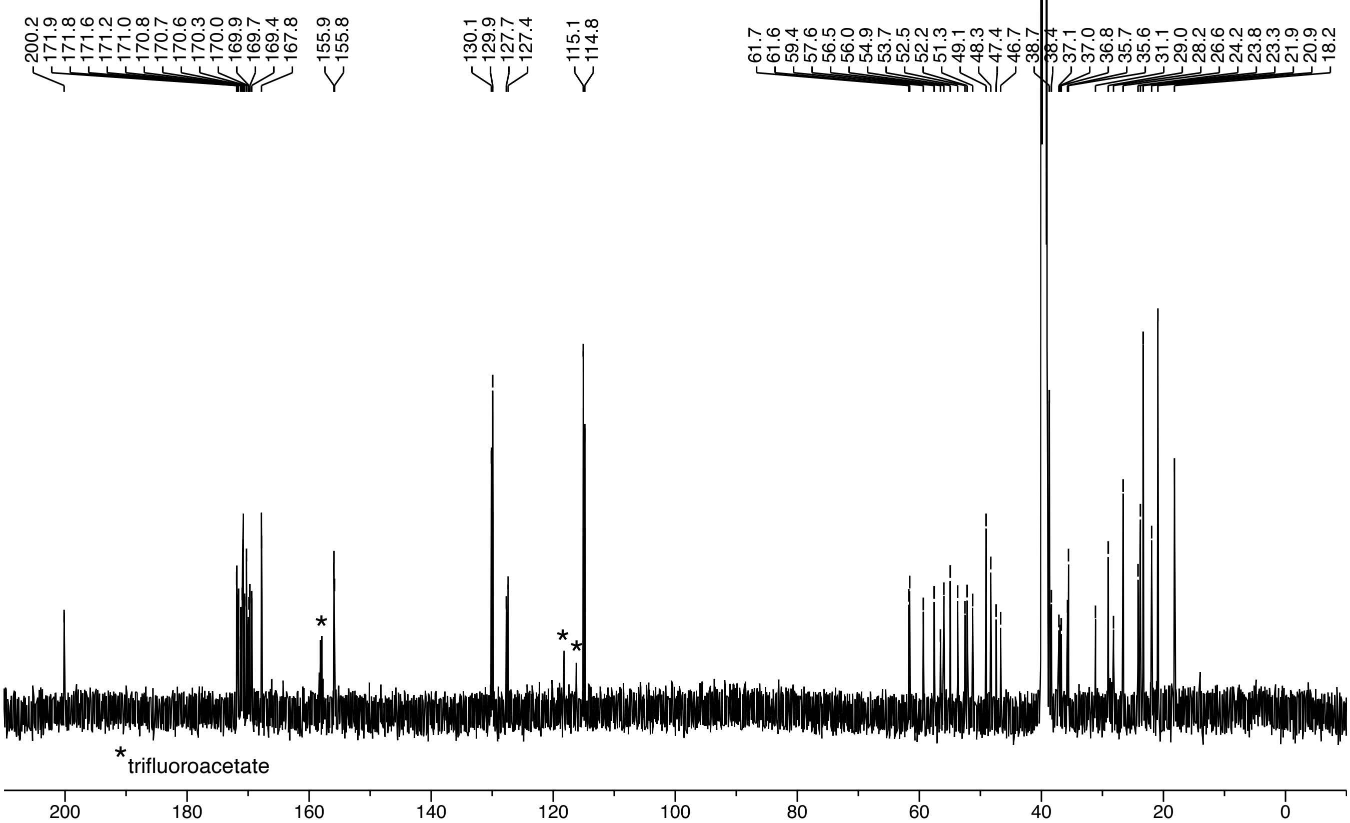

***S. epidermidis* AIP-III (7)**

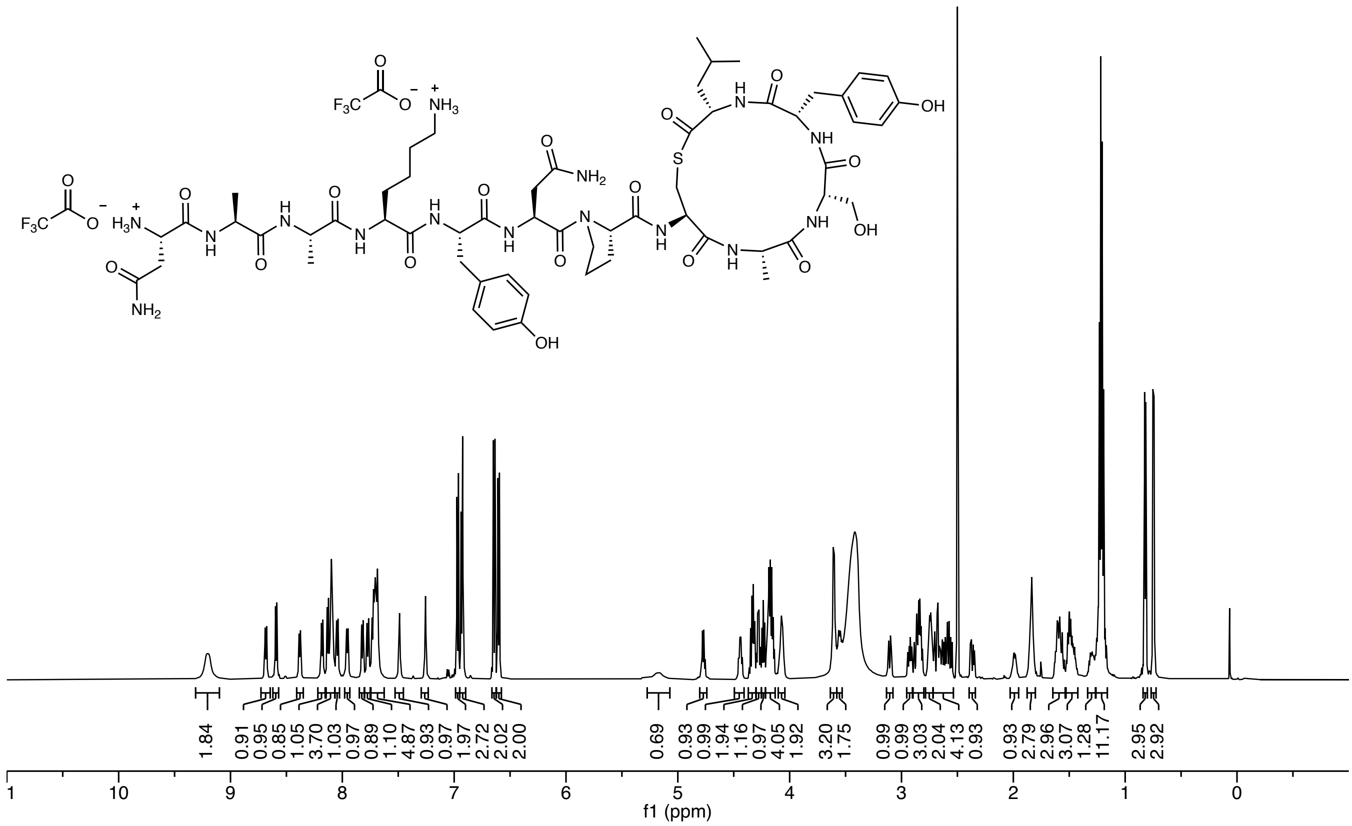

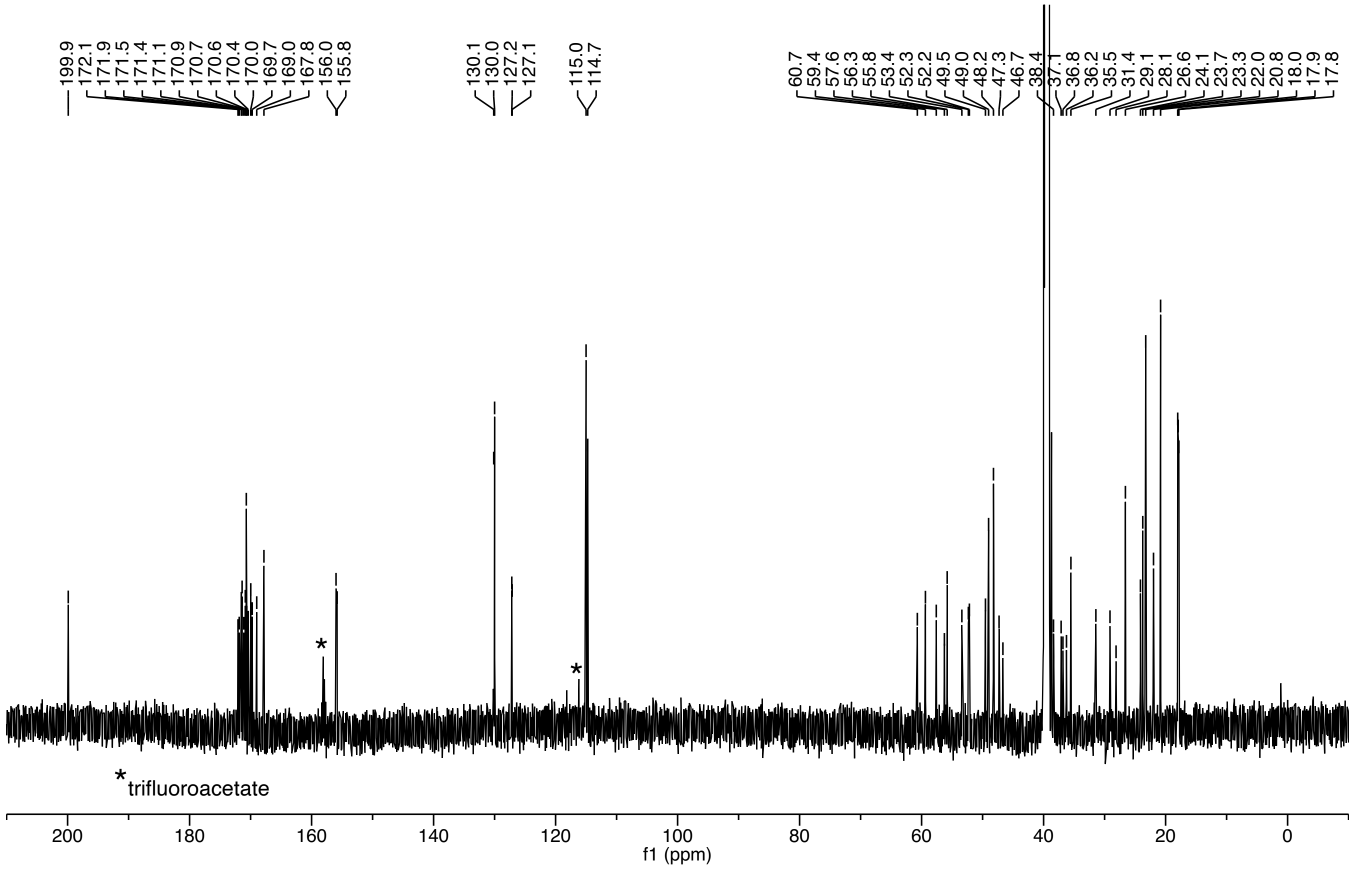

***S. lugdunensis* AIP-I (8)**

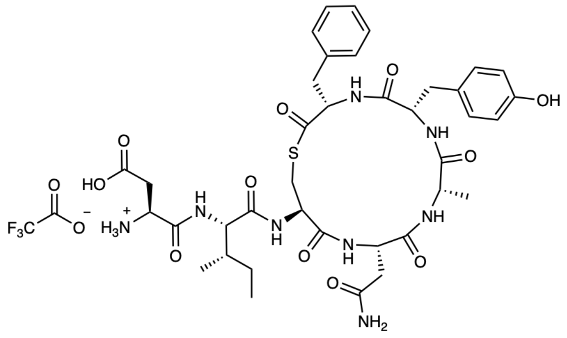

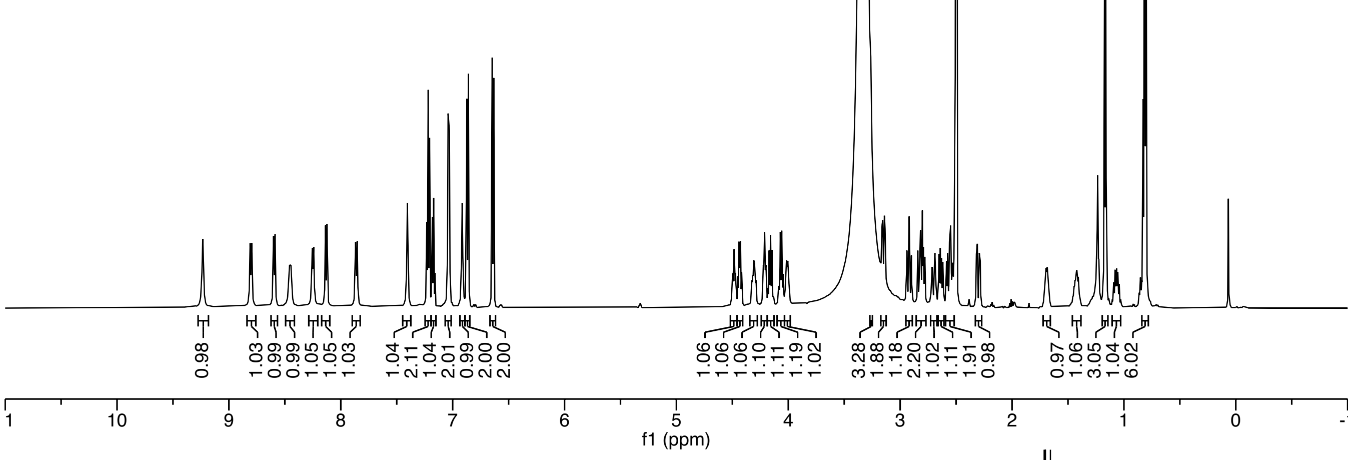

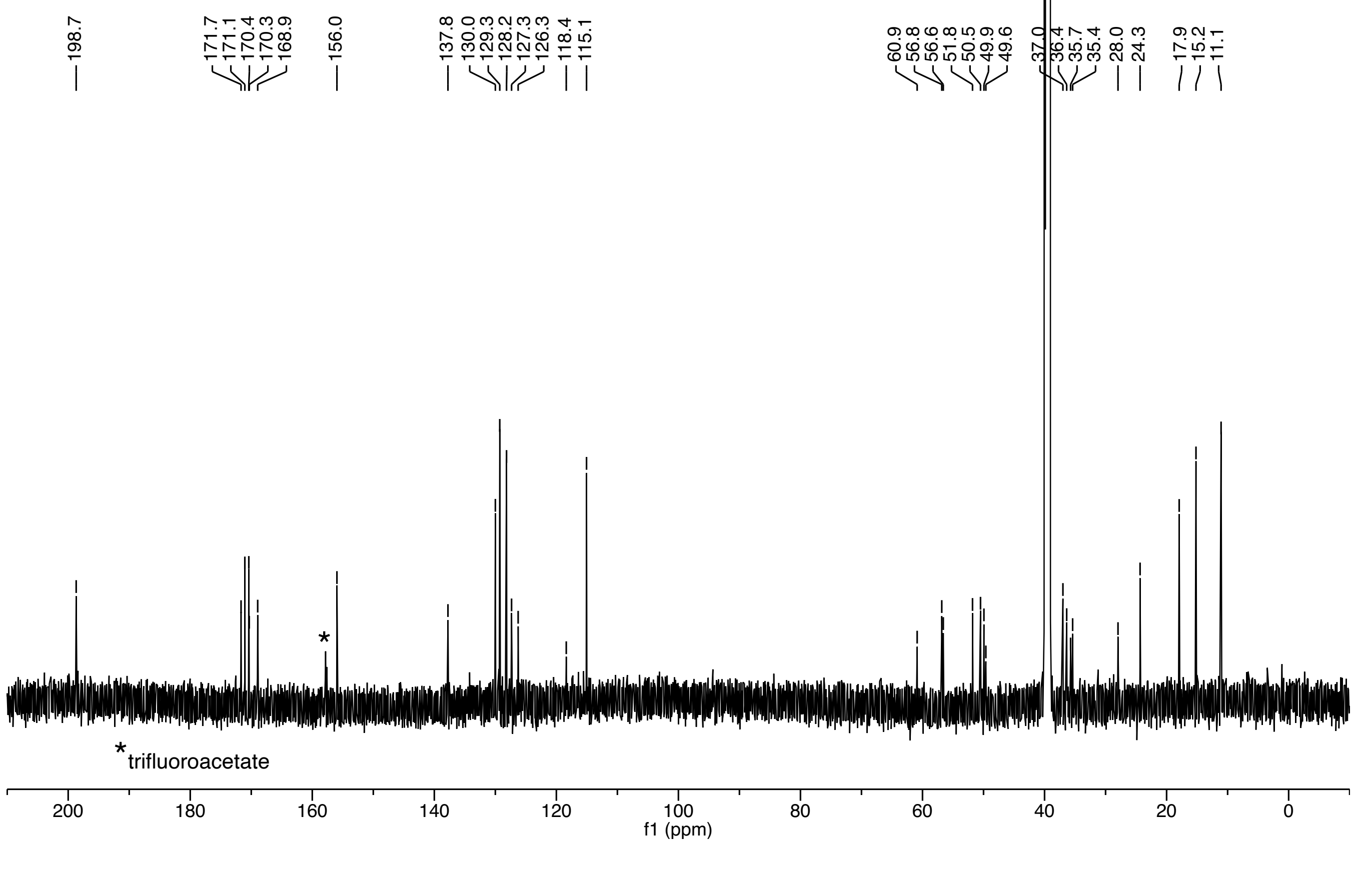

***S. hominis* AIP-II (11)**

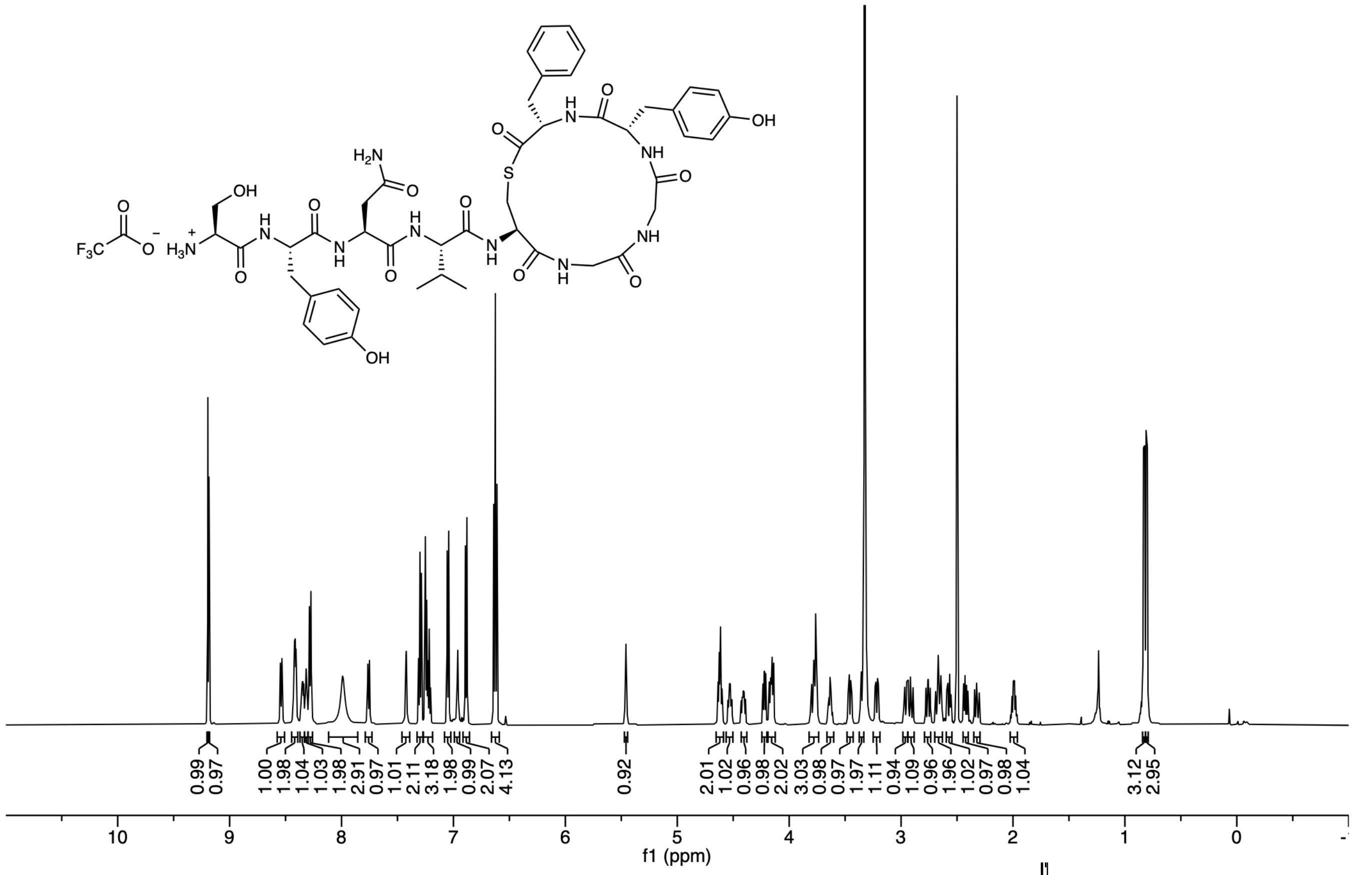

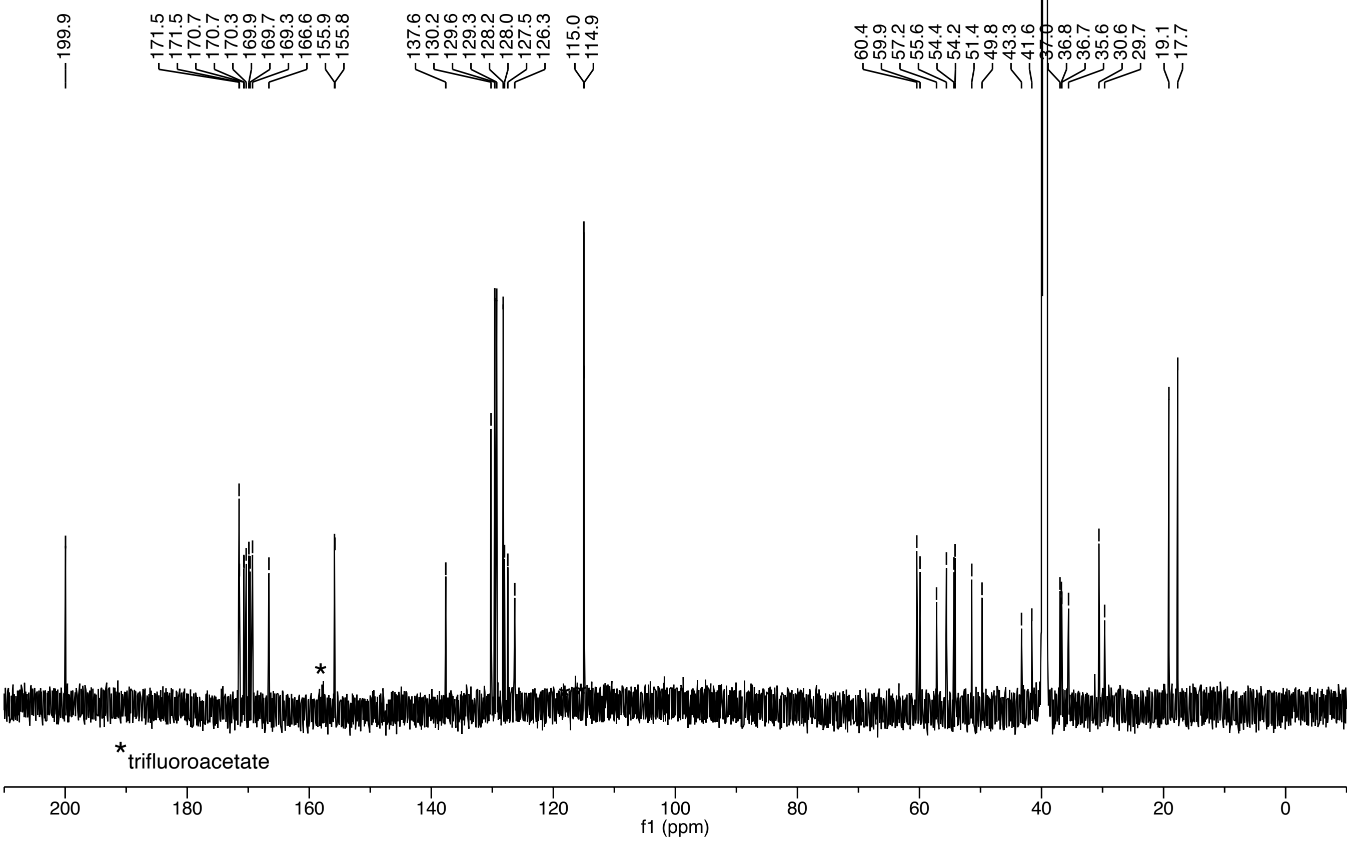

***S. caprae* AIP-I (12)**

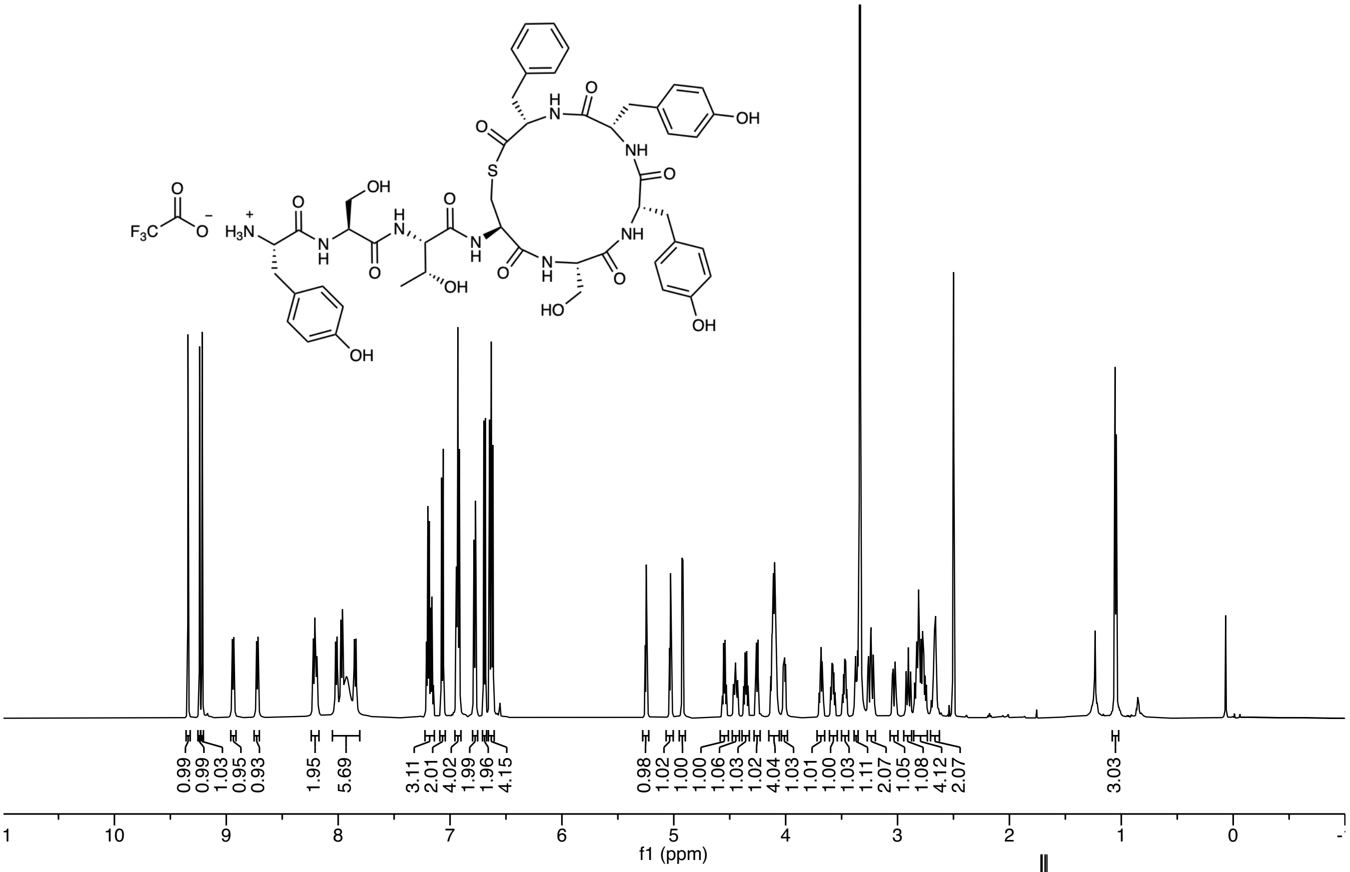

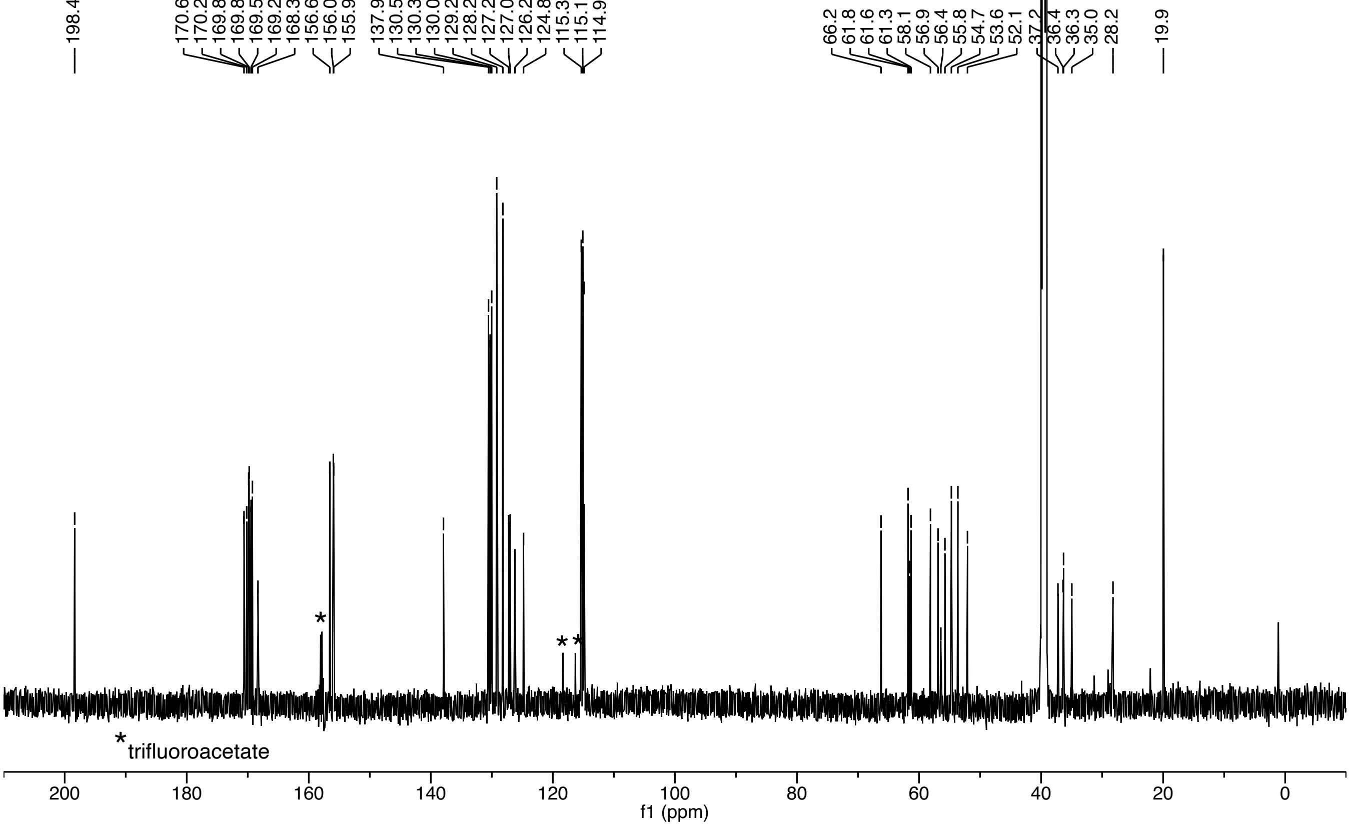

***S. pasteuri*** **AIP-I (15)**

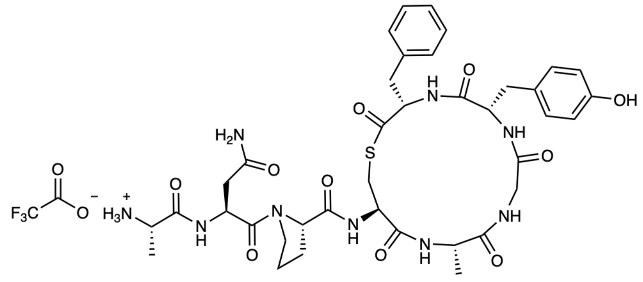

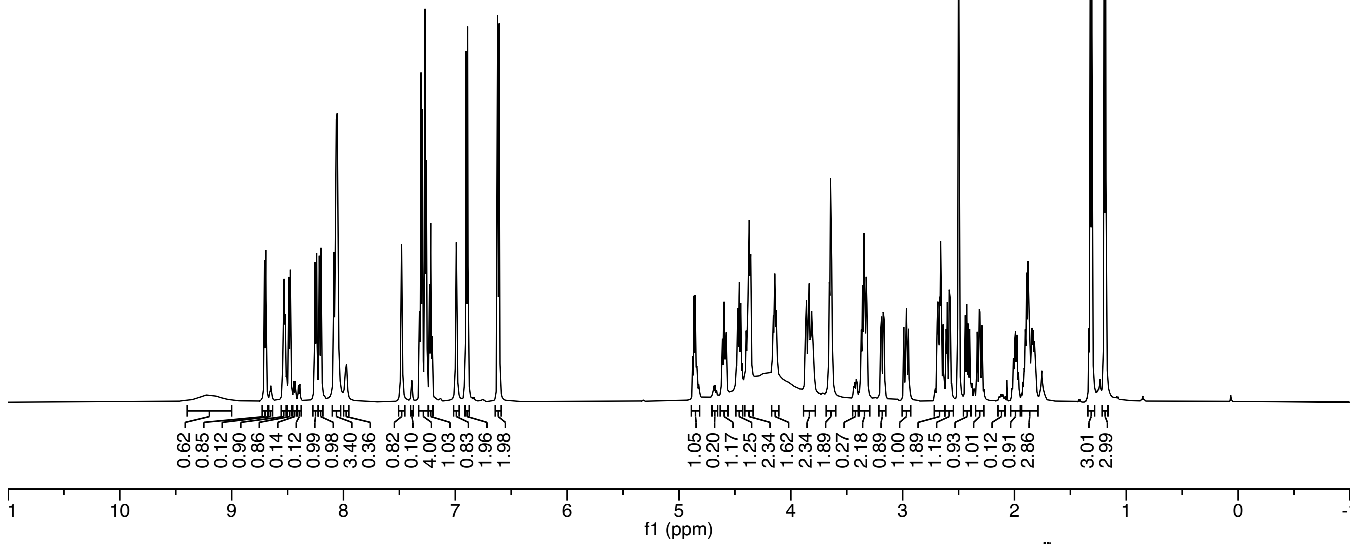

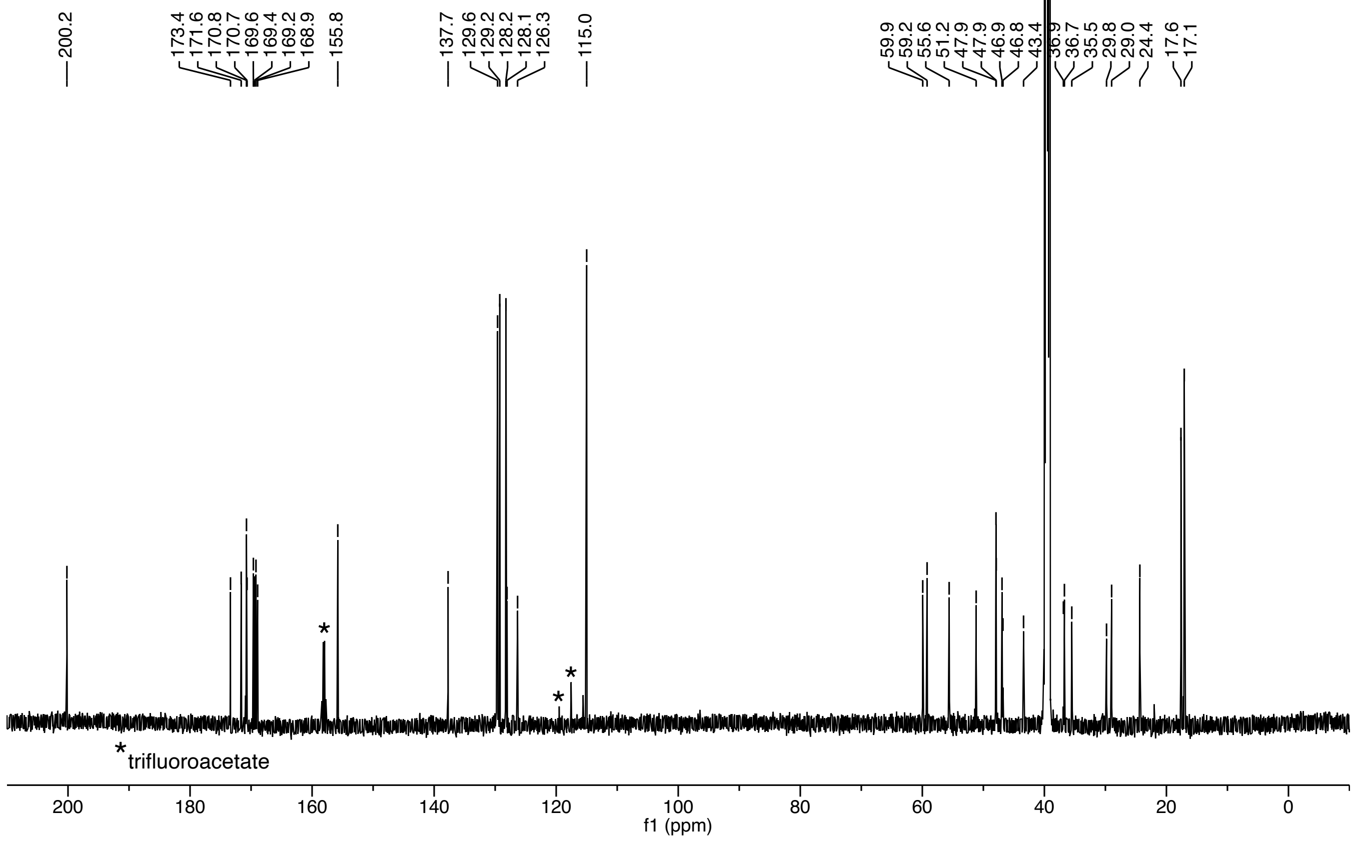

***S. succinus*** **AIP-I (16)**

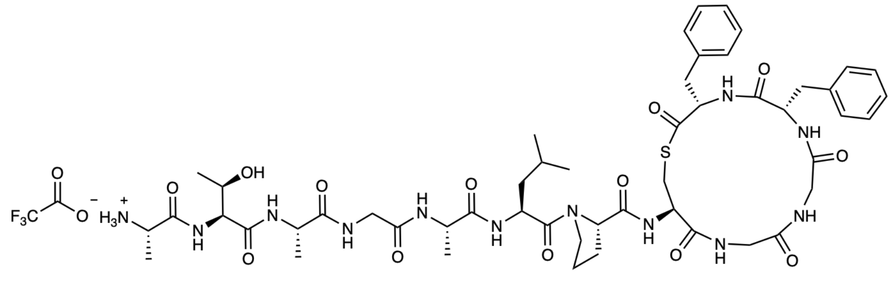

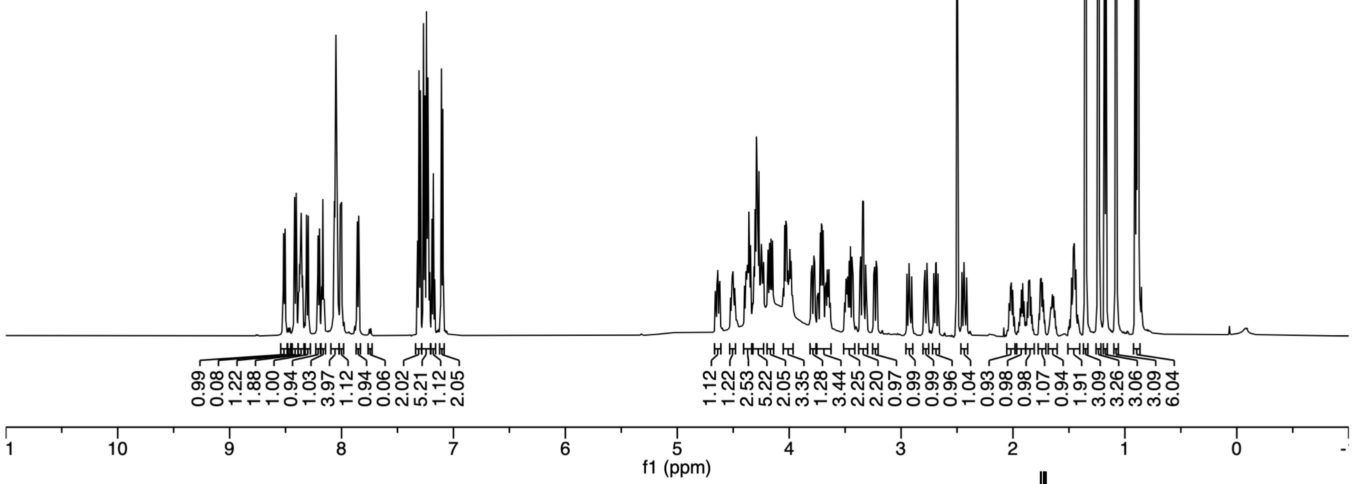

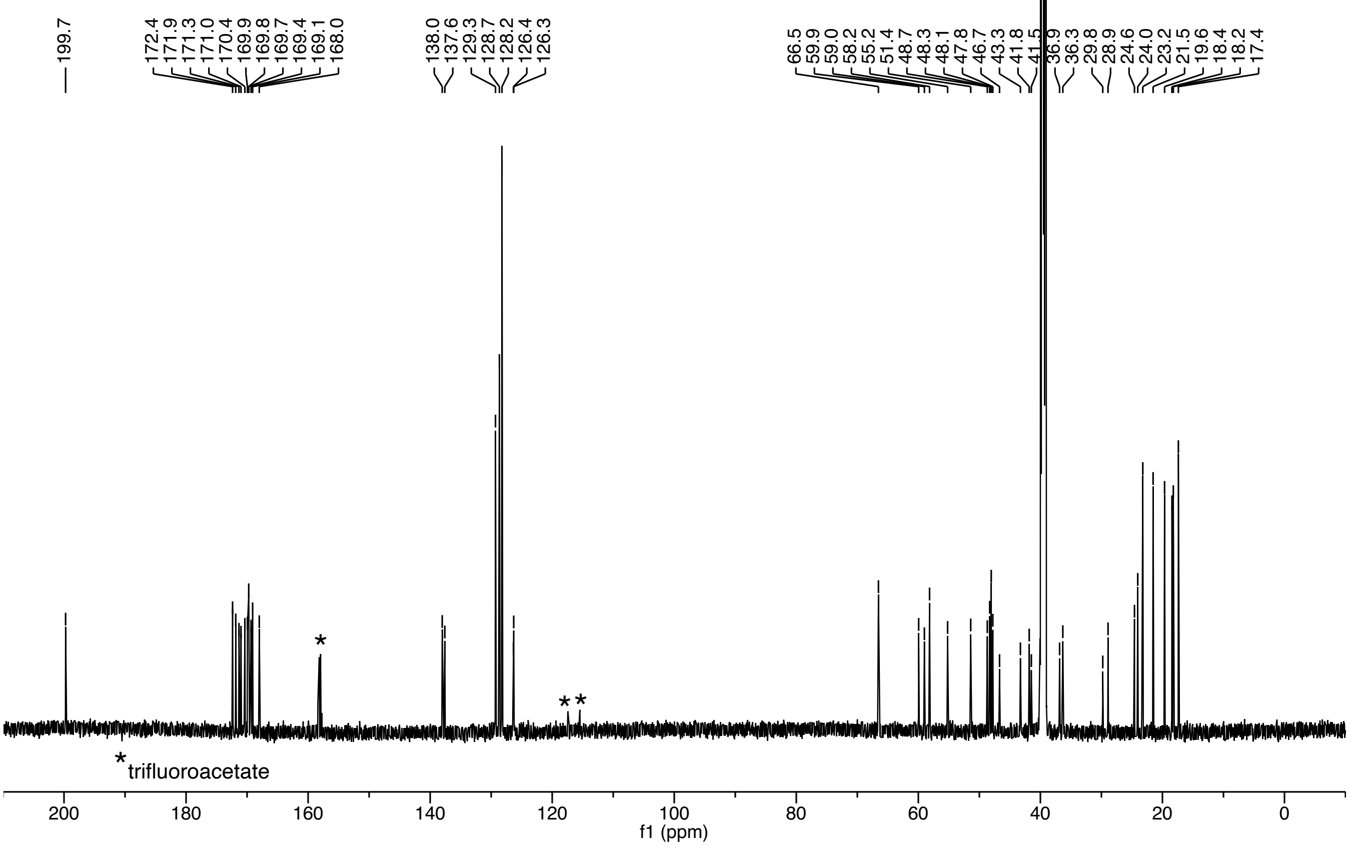

***S. cohnii* AIP-I (17)**

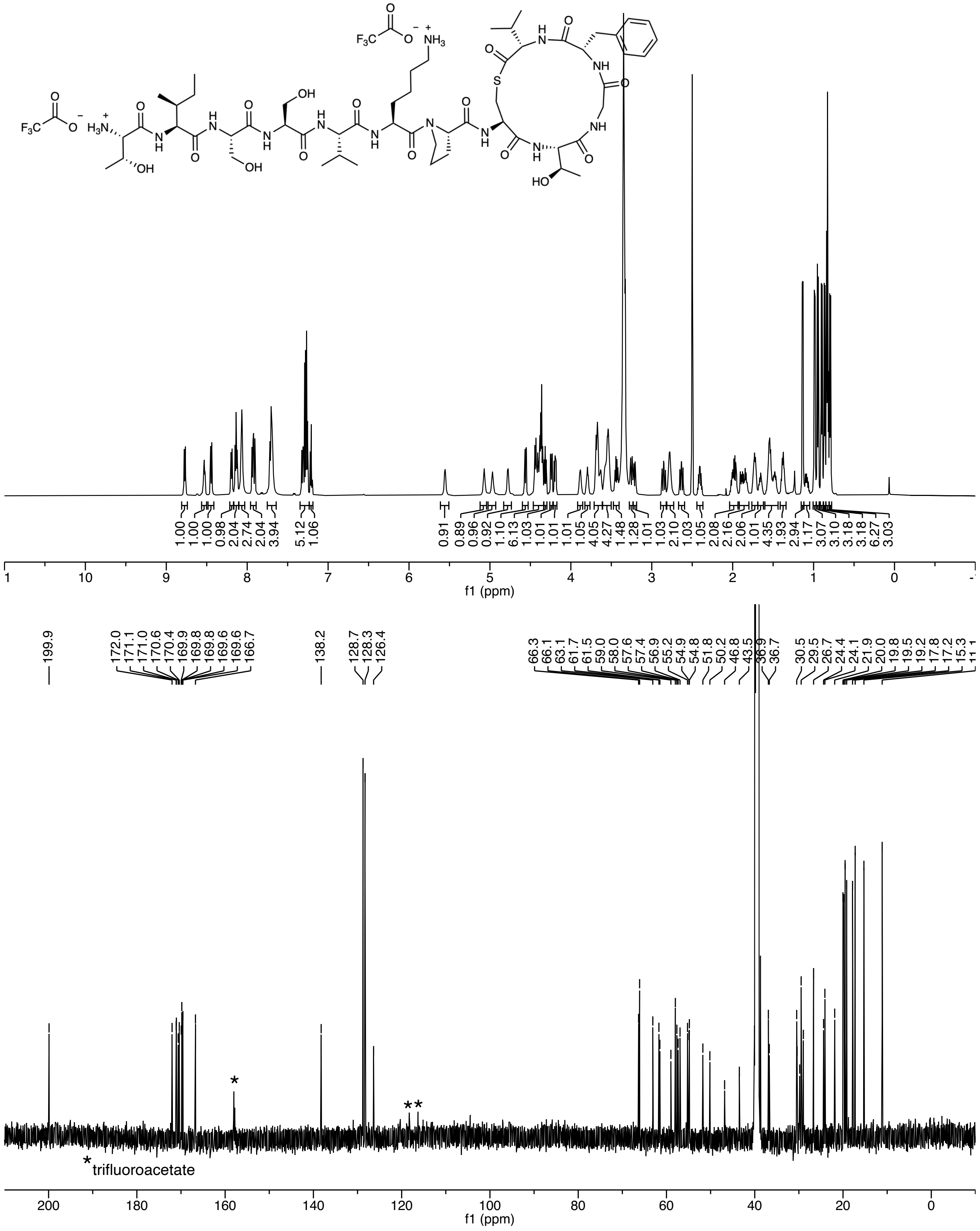

***S. intermedius* AIP-I (22)**

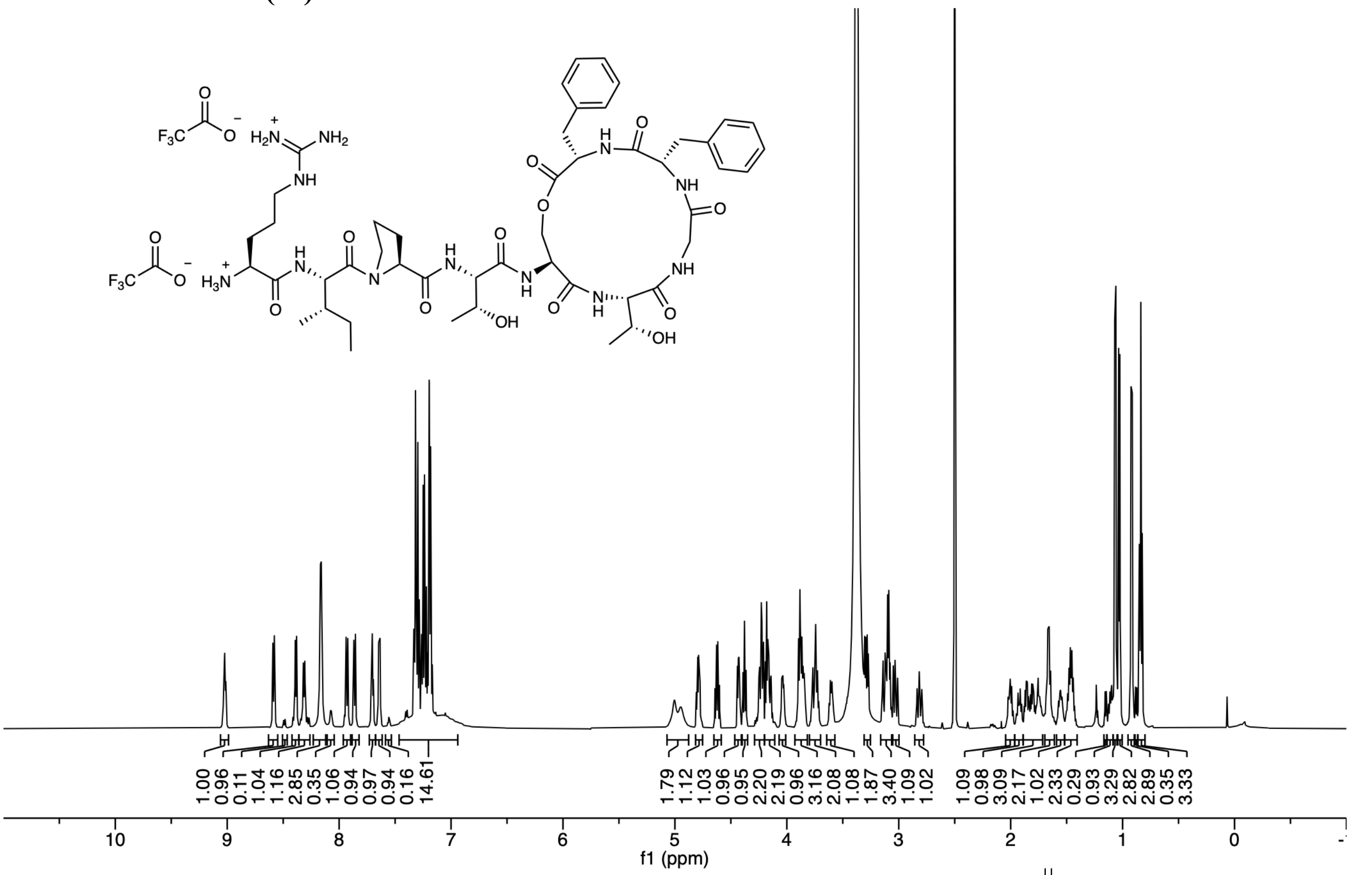

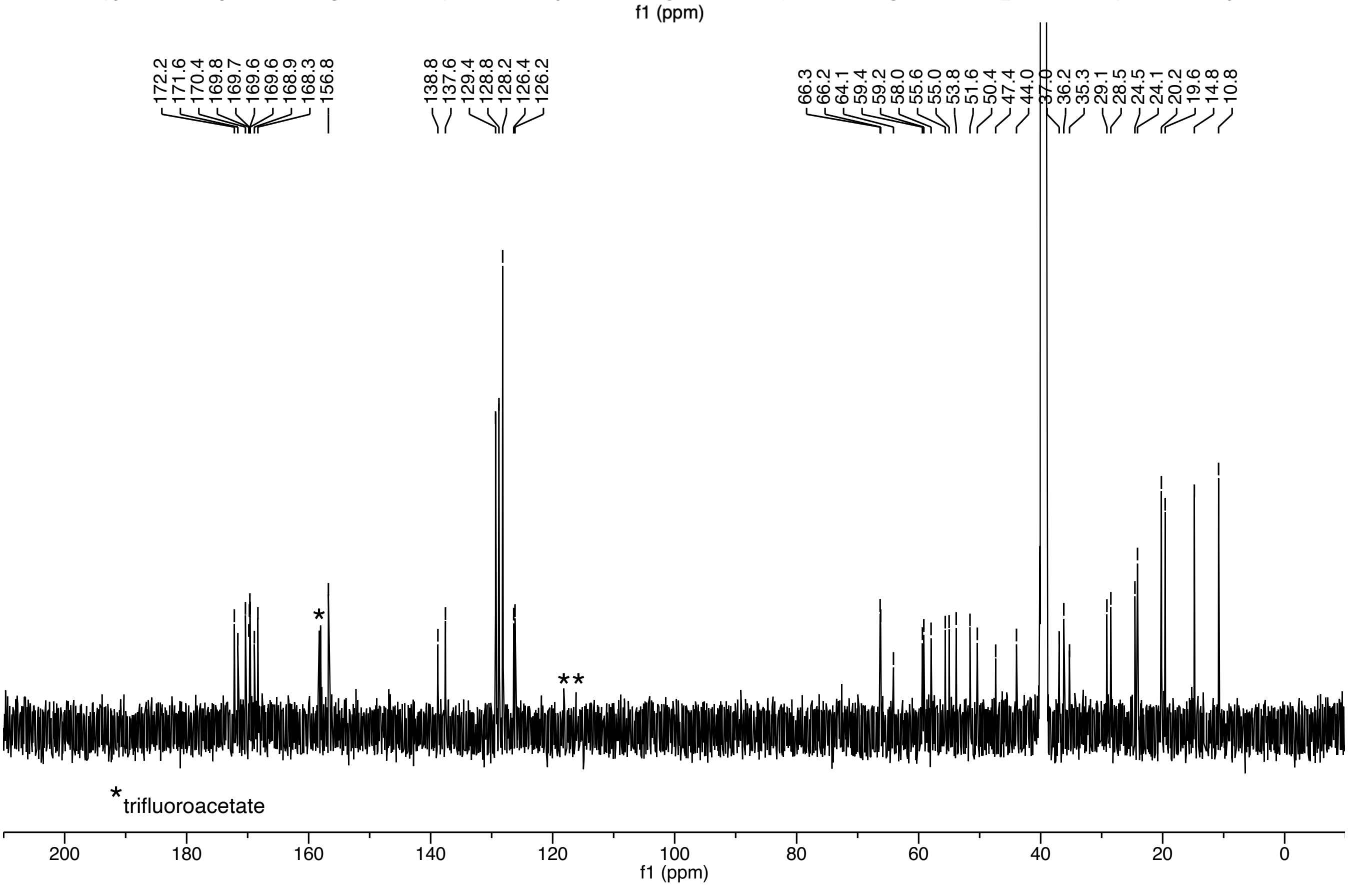

***S. simulans* AIP-II (24)**

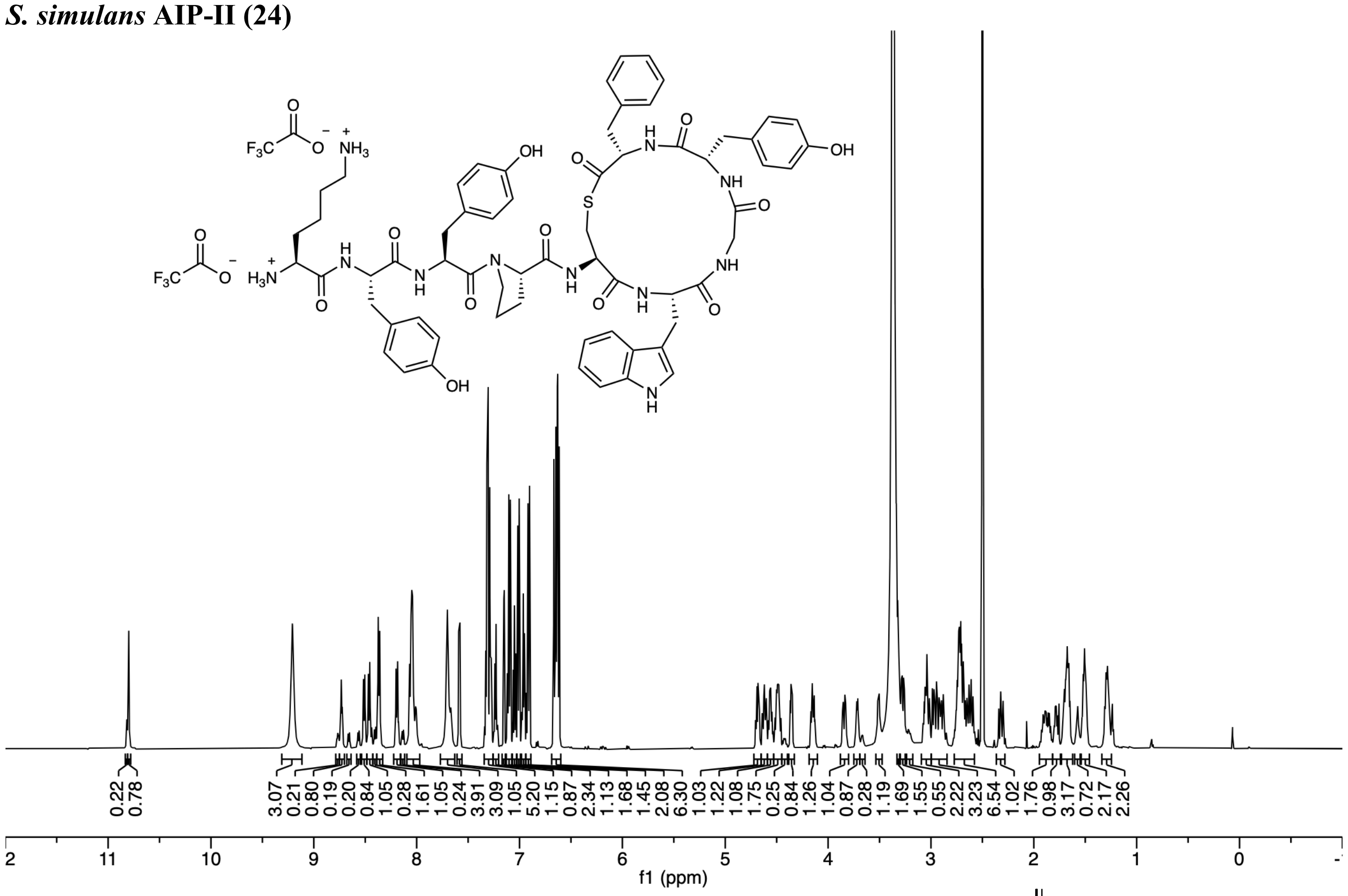

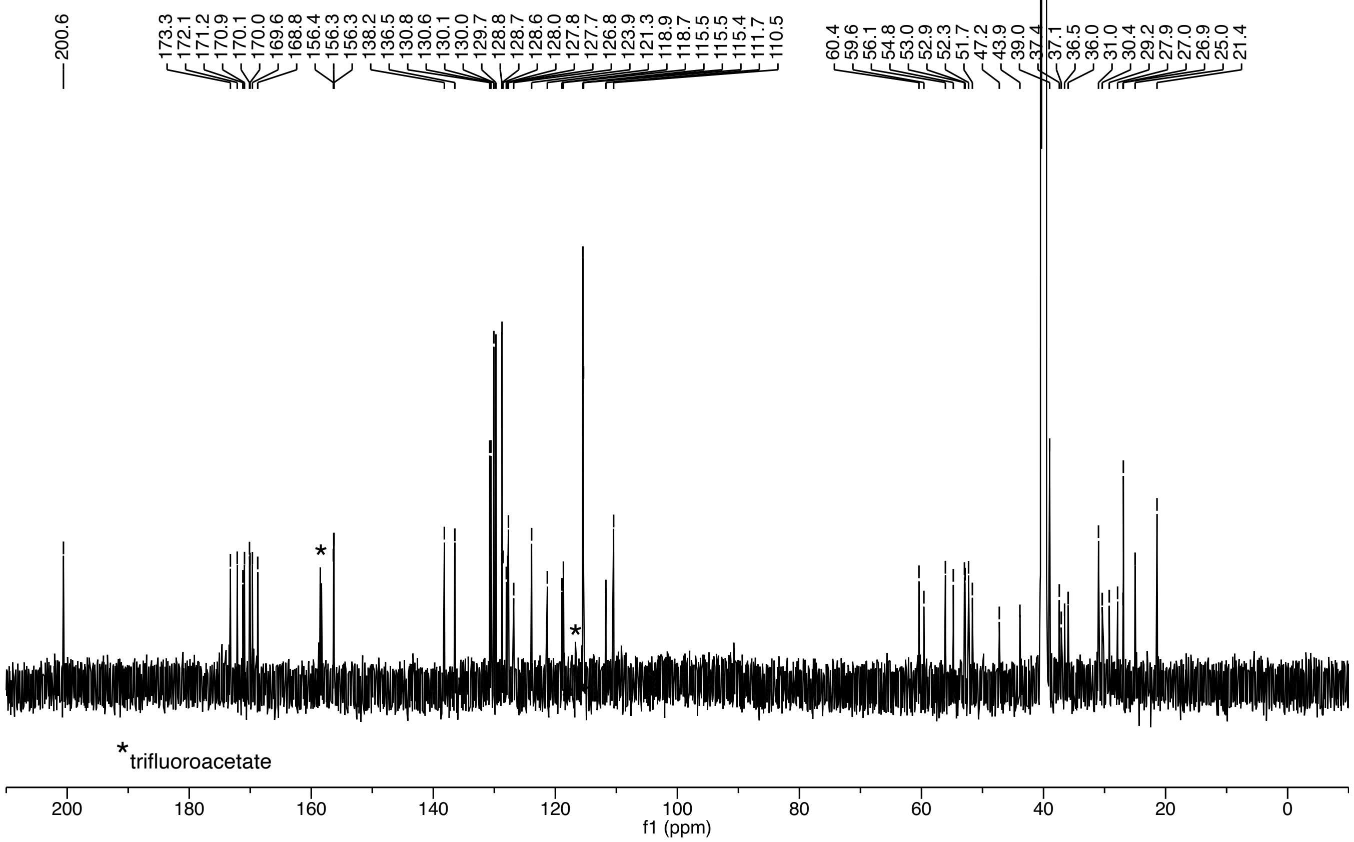

***S. simulans* AIP-III (25)**

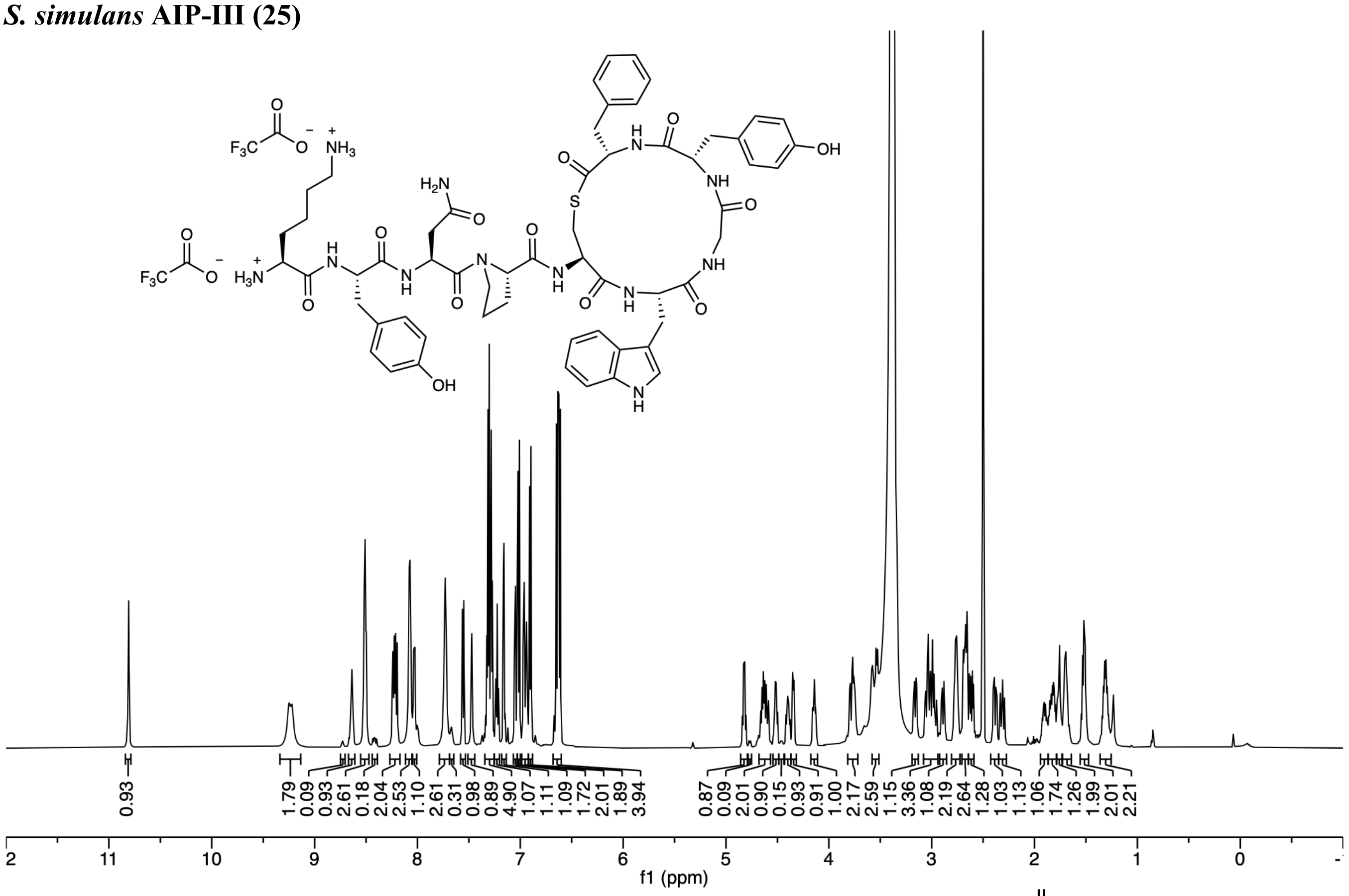

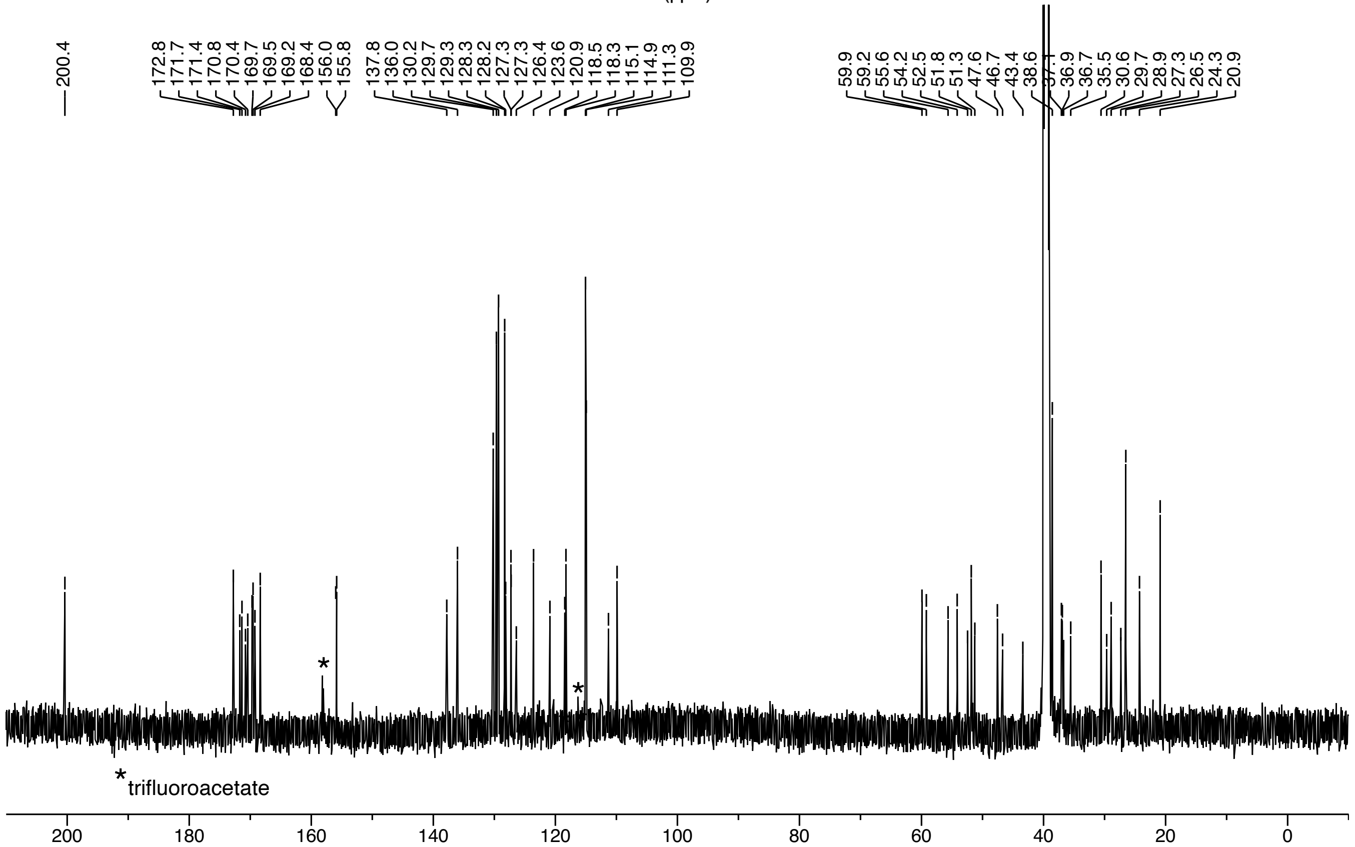
